# A Hymenoptera-restricted gene mediating ant castes co-opts deeply conserved machinery to control organ size

**DOI:** 10.64898/2026.08.31.748196

**Authors:** Ruyan Li, Sha Song, Xiafang Zhang, Rasmus Stenbak Larsen, Yanmei Qi, Yu Li, Fuqiang Lin, Joel Vizueta, Zijun Xiong, Jie Zhao, Dashuang Zuo, Wei Dai, Jixuan Zheng, Bitao Qiu, Yue Shi, Zijian Huang, Jiao Wang, Hao Ran, Qiye Li, Jacobus J. Boomsma, Xianjue Ma, Weiwei Liu, Guojie Zhang

**Author notes:** These authors contributed equally: Ruyan Li, Sha Song, Xiafang Zhang. Corresponding authors: Xianjue Ma, Weiwei Liu or Guojie Zhang.

## Abstract

Lineage-specific genes are widespread and have been implicated as phenotypic innovation inducers ^1^, but how they acquire complex developmental functions remains poorly understood. Ant queens and workers develop dramatically different organ sizes from identical genomes under juvenile hormone (JH) control ^2^, yet the molecular effectors translating JH signalling into caste-specific organ growth remain unknown. Here we identify *torch*, a Hymenoptera-restricted gene, as the most consistently gyne-biased and JH-responsive gene across 68 ant species. Knockdown of *torch* in virgin queens of *Monomorium pharaonis* produces a worker-like, multi-organ growth-restricted phenotype. Mechanistically, *torch* harbours an E-box-like motif activated by the JH receptor Gce–Tai and acts as a GA-repeat-binding transcription factor that regulates Hippo signalling, the deeply conserved organ-size control pathway in animals. Expressing *torch* heterologously in mice and a growth-restricted *Drosophila* background shows that the gene retained its general growth-promoting activity across more than 700 million years of animal evolution in lineages that lack the gene, establishing that its function is mediated through conserved rather than ant-specific machinery. A lineage-specific gene can therefore acquire complex morphogenetic function by co-opting ancient organ-size circuitry, providing a general route by which novel genes can drive phenotypic innovation.

## Introduction

Whether new biological functions evolve by novel biochemistry or by recycling already existing mechanisms is a long-standing puzzle ^3–5^. Novel genes, taxonomically restricted to particular clades, have been conjectured to play important roles in lineage-specific adaptation ^1,6^ but how they acquire complex developmental functions remains poorly understood. Strikingly, lineage-specific genes sometimes retain activity when expressed in distantly related genomes that lack them, suggesting that their function depends on machinery shared across lineages. For example, the *Arabidopsis*-specific gene *QQS* enhances protein content at the expense of starch in distantly related crops ^7^, the bee-specific venom peptide apamin ^8^ mediates antimicrobial activity in *Drosophila* guts ^9^, and the human-specific genes *SP0535* and *SMIM45* promote cortical expansion in mice ^10,11^. These and related findings ^12–21^, indicate that lineage-specific genes may functionally be more deeply rooted than their restricted taxonomic distribution suggests.

Ant caste-differentiation is one of the most distinctive evolutionary innovations because a single genome produces two dramatically different adult female phenotypes, gynes and workers, potentially differing by orders of magnitude in body size ^22^. Gynes (virgin future queens), have fully developed wings and flight muscles, expanded brains, prominent compound eyes and ocelli, large fat bodies and functional reproductive organs, whereas workers have reduced or vestigial versions of these traits ^2,23^. Inseminated queens also live much longer than workers ^24^, a divergence driven by juvenile hormone (JH) titers that bifurcate larval development towards gyne or worker phenotypes during critical time-windows ^25–36^. Yet despite five decades of work establishing JH as the master endocrine regulator of caste differentiation, the molecular effectors that translate hormonal signals into divergent multi-organ growth programs are unknown. The Hippo pathway is a conserved organ size control network across animals ^37,38^ and represents a strong candidate for mediating these effects. Hippo may integrate endocrine signals during insect appendage growth ^39^ but what extends JH signaling into Hippo-mediated organ size control remains unidentified. Because a substantial fraction of gyne and worker biased transcripts in ants derive from lineage-restricted genes ^40,41^, we hypothesized that pre-imaginal caste differentiation represents an evolutionary innovation by itself, based on lineage-specific gene recruitment into an ancient growth-control network.

Here we identify *torch*, acronym for Taxon-restricted Organ-size Regulator Co-opting Hippo, a Hymenoptera-restricted gene with no detectable homologs outside this insect order, as a key regulator required for general queen-specific organ growth in the ant *Monomorium pharaonis*. We show that *torch* is transcriptionally induced by JH signaling and in turn activates the Hippo pathway to drive coordinated growth across multiple gyne-specific organs, revealing how a lineage-specific gene can be recruited into an ancient, non-social hormonal cascade to produce caste-specific morphology in ants. We then test this co-option model by expressing *torch* in Hippo-sensitized *Drosophila* backgrounds and in mice, directly demonstrating that its morphogenetic function operates through deeply conserved machinery rather than ant-specific biochemistry.

## Results

### *torch*: a Hymenoptera-specific JH responsive gene with conserved gyne-biased expression

To identify conserved caste-specific genetic signatures, we analyzed adult transcriptomes from 68 ant species spanning 46 genera and seven subfamilies ^23^, capturing much of the extant phylogenetic ant diversity (Fig. 1a). Across these lineages, *torch* was the most consistently gyne-biased gene, upregulated relative to workers in 57 of the 65 species possessing a detectable ortholog (Fig. 1b; Supplementary Dataset 1), a pattern retained across the deep formicoid-poneroid split (Extended Data Fig. 1a) and the five major ant subfamilies covered (Extended Data Fig. 1b). This distribution implies that expression bias was maintained for more than 150 million years of ant evolution ^23^. Furthermore, *torch* expression correlates with the extent of queen–worker caste dimorphism (PGLS *P* = 0.0008; Extended Data Fig. 1c), consistent with the gene being overexpressed in gynes relative to workers from the second larval instar onwards in both *M. pharaonis* with a single worker caste and in *Acromyrmex echinatior*, a leaf-cutting ant with 2-3 discrete worker subcaste ^42^ (Fig. 1c-d). Expression of *torch* was also elevated in larger soldiers relative to minor workers of *Carebara diversa* (Extended Data Fig. 1d) and in major workers relative to minor workers in *Camponotus floridanus* (log_2_ fold change = 2.9, adjusted *P* = 0.005) ^43^. The *torch* gene thus emerged as a candidate general organ-size regulator whose expression tracks larger caste phenotypes across the Formicidae.

**Fig. 1.**
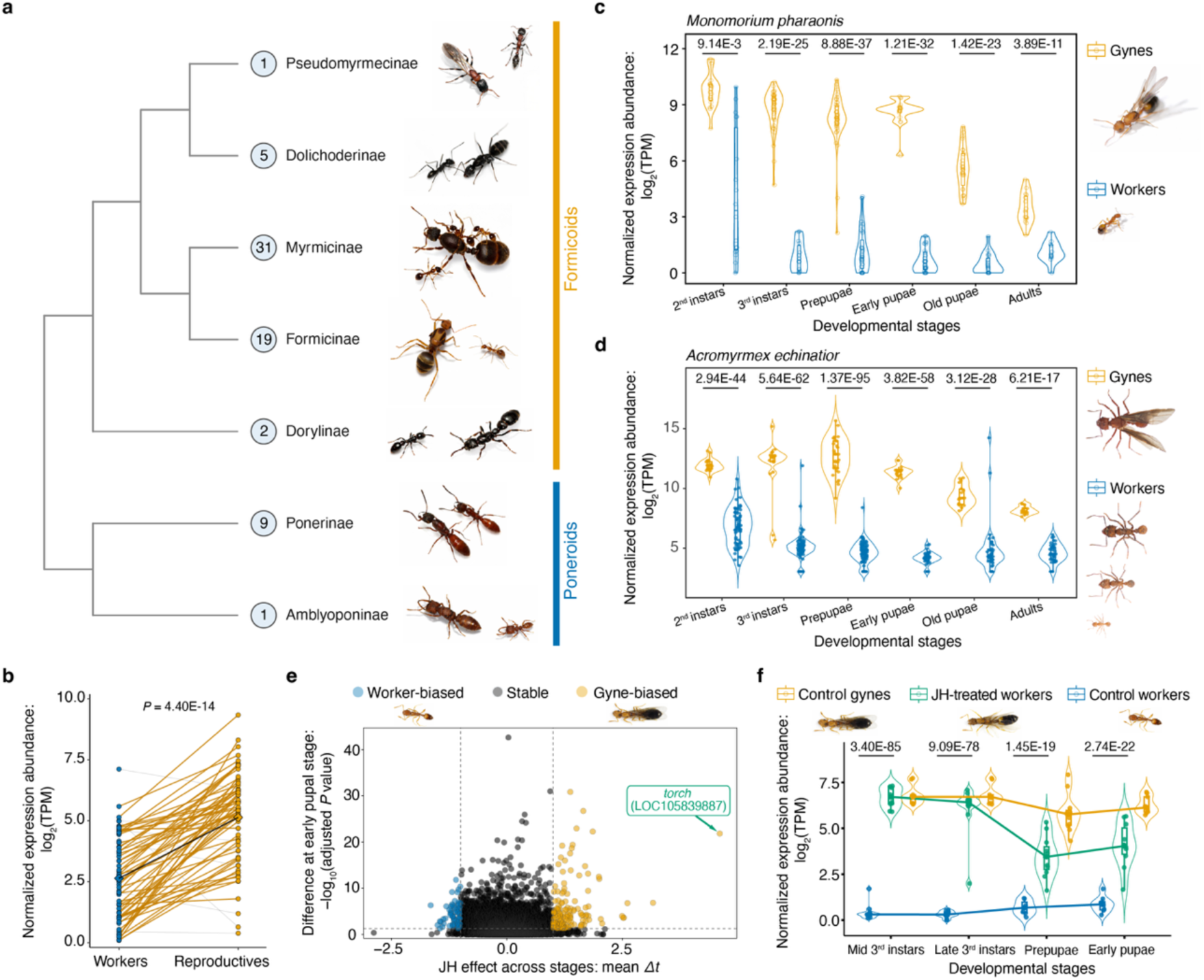
The *torch* gene as a Hymenoptera-restricted, JH–inducible gene with conserved gyne-biased expression across the ants. **(a)** Phylogenetic coverage of the adult transcriptomes analyzed across seven ant subfamilies (numbers of species analyzed in circles) with representative images (top to bottom): GAGA-0365 (*Tetraponera rufonigra*), GAGA-0363 (*Iridomyrmex anceps*), GAGA-0407 (*Kartidris nyos*), GAGA-0199 (*Anoplolepis gracilipes*), GAGA-0379 (*Cerapachys sulcinodis*), GAGA-0408 (*Buniapone amblyops*), and GAGA-0404 (*Mystrium camillae*). **(b)** Gyne-biased over-expression of *torch* across 65 ant species based on mean normalized TPM values averaged across replicates, with diamonds and black line representing the averages of the yellow lines (two-sided paired *t*-test, *t* = 9.64, df = 64). **(c-d)** Developmental transcriptomes for *torch* showing consistent gyne-biased expression relative to workers from second larval instar to adults in *Monomorium pharaonis* (c) and *Acromyrmex echinatior* (d) after pooling data from small, medium and large workers ^42^. **(e)** Meta-analysis volcano plot for all genes expressed across the five (pooled) developmental stages of *M. pharaonis*, identifying *torch* (LOC105839887) as the most strongly JH-responsive gene with by far the largest mean *Δt* value for differential expression across all five developmental stages together with a high −log_10_ adjusted *P* value for JH-treated workers versus control workers in the early pupal stage (y-axis). See Supplementary Dataset 3 and ^2^ for original data and Methods for analysis details. **(f)** Normalized RNA-seq developmental time course of *torch* expression in a JH treatment assay of *M. pharaonis* worker larvae using the same published data as in the e-panel, with JH-treated workers showing *torch* expression approaching levels in control gynes and exceeding those in control workers across developmental stages. For comparisons using RNA-seq data (c, d, and f), statistical significance is reported as DESeq2-derived adjusted *P* values.

Comparative genomic analyses revealed that *torch* has no detectable homologs outside the Hymenoptera (Extended Data Fig. 1e). The ant stem lineage experienced strong positive selection on the *torch* coding sequence and a pronounced structural truncation so the ant orthologs uniformly lack a C-terminal segment present in bee and wasp orthologs (Extended Data Fig. 1e-h; Supplementary Dataset 2). This combination of adaptive sequence evolution and structural shortening at the origin of ant-specific caste differentiation suggests that *torch* underwent functional remodeling when the ant-specific caste system emerged as part of an irreversible major transition in evolution (MTE) to superorganismal organization ^23^. The *torch* gene also resides within a conserved microsynteny block maintained across the Formicidae (Supplementary Fig. 1), consistent with cis-regulatory preservation underlying conserved caste-biased expression ^44^.

We previously showed that JH treatment of worker larvae induces gyne-like organ enlargement in multiple tissues, including imaginal disc-derived organs, muscle, fat body, and brain ^2^, so the organ-size correlated expression pattern of *torch* led us to hypothesize that the gene functions as a downstream effector of JH signaling. We therefore compared transcriptomes from JH-treated versus control workers of *M. pharaonis* ^2^ and identified *torch* as the most JH-responsive gene in the entire transcriptome across developmental stages (Fig. 1e; Supplementary Dataset 3). JH treatment elevated *torch* expression in workers to levels comparable to those naturally observed in gyne-destined larvae (Fig. 1f). RNA-FISH confirmed elevated *torch* signaling in the fat bodies of JH-treated worker larvae (Extended Data Fig. 1i), corroborating *torch*’s candidacy as effector of JH signaling towards caste-differentiated organ growth.

In panel f, the comparison is between JH-treated workers and control workers. Box plots show the median, 25th and 75th percentiles, outermost values (whiskers), and individual data points, a convention used throughout the manuscript.

### Loss of Torch redirects gyne development toward a worker-like trajectory

To test whether Torch is functionally necessary for the enhanced organ growth that distinguishes gynes from workers, we depleted *torch* during gyne development using splice-blocking vivo-morpholinos (vivo-MOs), cell-permeable antisense oligonucleotides that enable post-embryonic loss-of-function assays ^45^ (Extended Data Fig. 2a–c; see Supplementary Information Text for detailed procedures). This revealed that *torch*-depleted gyne larvae had markedly smaller adipocytes and lipid droplets 24 h after vivo-MO injection (Fig. 2a–c), phenocopying worker-like attenuation of fat-body growth ^2^. Knockdown also advanced the onset of metamorphosis by ∼1 day compared with controls (*P* = 0.008; Extended Data Fig. 2d–e), consistent with *torch* promoting late-larval organ growth while restraining precocious metamorphosis.

**Fig. 2.**
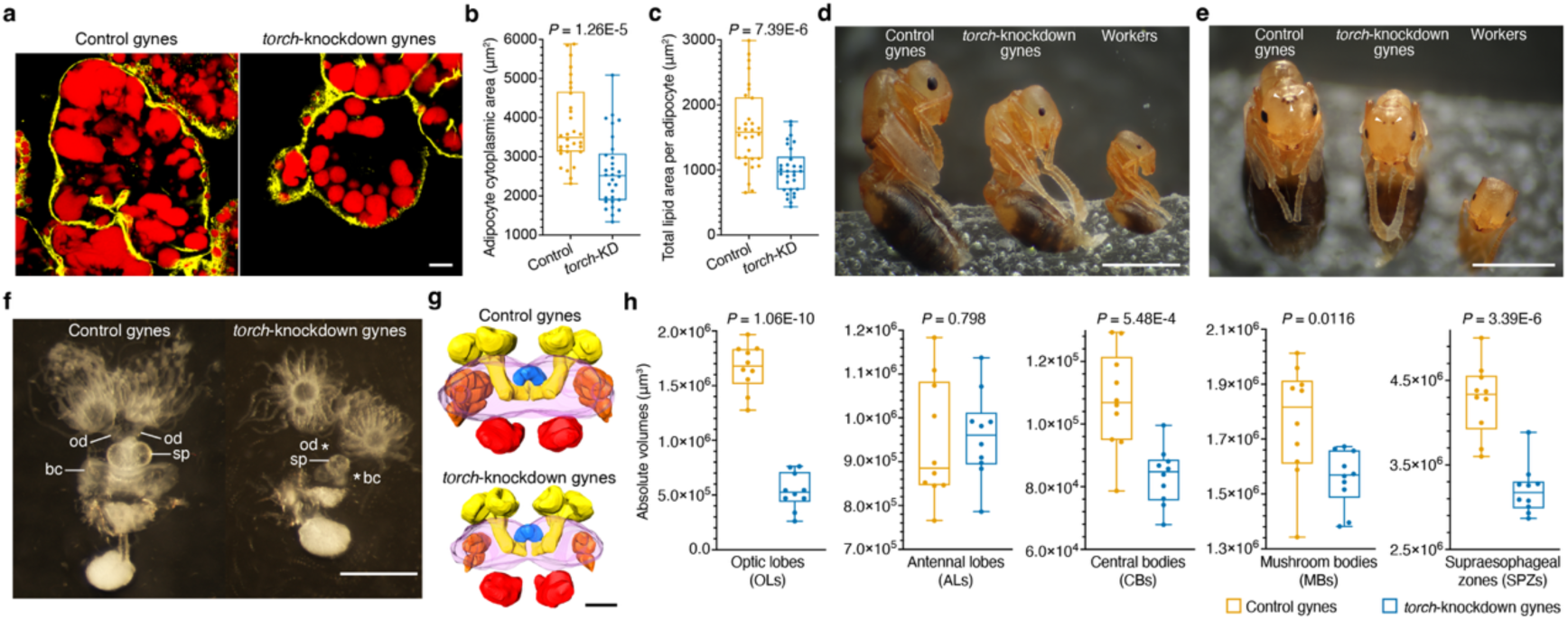
*Torch* knockdown inhibits gyne organ growth. **(a)** Late third-instar larval adipose tissue 24 h after vivo-MO injection, stained with HCS LipidTOX™ Red (neutral lipids; red) and Alexa Fluor™ 546-conjugated phalloidin (F-actin; yellow), were markedly reduced. **(b-c)** Concomitant quantitative reduction of cytoplasmic area (b) and total lipid area (c) of adipocytes in third-instar gyne larvae 24 h after *torch*-knockdown; *n* = 30 adipocytes per group. Statistical significances were assessed using two-sided Student’s unpaired *t*-tests: cytoplasmic area, *t* = 4.78, df = 58; total lipid area, *t* = 4.92, df = 58. Control denotes control gynes, and *torch*-KD denotes *torch*-knockdown gynes. **(d-e)** Representative lateral and top-down morphology of day-12 pupae of control gynes, *torch*-knockdown gynes and workers. White arrowheads (e) highlight ocelli in control and *torch*-knockdown gynes which are absent in workers. **(f)** Dissected female reproductive tracts from day-12 pupae with control gynes showing normal oviducts (od), spermatheca (sp) and bursa copulatrix (bc) whereas *torch*-knockdown gynes have a shrunken spermatheca and no bursa copulatrix and oviducts (asterisks). **(g)** Representative 3D reconstructions of day-12 pupal brains from control and *torch*-knockdown gynes, generated from confocal optical sections of whole-mount heads (Extended Data Fig. 2o). **(h)** Volumes of major brain neuropils quantified from 3D brain reconstructions, including optic lobes, antennal lobes, central body, mushroom bodies and supraesophageal zone. For bilateral neuropils, including optic lobes, antennal lobes and mushroom bodies, paired left–right volumes were summed per brain before statistical analysis. Neuropil volumes were significantly reduced after *torch* knockdown, except for antennal lobes; *n* = 10 reconstructed brains per group. Statistical significances were assessed using two-sided Student’s unpaired *t*-tests: optic lobes, *t* = 13.21, df = 18; antennal lobes, *t* = 0.2593, df = 18; central body, *t* = 4.192, df = 18; mushroom bodies, *t* = 2.811, df = 18; supraesophageal zone, *t* = 6.598, df = 18. Scale bars, 10 μm (a), 1 mm (d, e), 500 μm (f) and 100 μm (g).

Because major expansion of the imaginal-disc-derived organs, flight muscles and brain neuropils occurs during the prepupal and pupal stages ^46–49^, we asked whether *torch* is necessary for this growth burst towards larger adult organ size. By the final day of pupation (day 12), *torch*-depleted gynes exhibited a 10% reduction in body size and coordinated size reductions in the imaginal-disc-derived structures: forewings (50%), head capsule (8%), antennal scapes (10%), compound eyes (55%), ocelli (80%), and the flight-muscle-containing mesosoma (30%). Internal reproductive structures were similarly affected: the bursa copulatrix and oviducts did not develop, and the spermatheca was reduced by 75% (all *P* < 0.0001; Fig. 2d–f; Extended Data Fig. 2f– m). In contrast, ovaries remained prominently present in *torch*-depleted gynes (Fig. 2f), consistent with germline canalization starting unusually early in this ant species ^2,50^, i.e. well before the late third-instar stage at which *torch* knockdown was initiated (Extended Data Fig. 2n).

Torch depletion also profoundly impaired brain development. Three-dimensional brain reconstructions of day-12 pupal brains revealed significant reductions in neuropil volume in the optic lobes, central body, mushroom bodies, and supraesophageal zone (Fig. 2g–h; Extended Data Fig. 2o). Beyond size reduction, Torch depletion also disrupted caste-specific neural patterning: a gyne-specific serotonergic neuron cluster in the medulla, absent but JH-inducible in workers, was markedly reduced in *torch*-depleted gynes (Extended Data Fig. 2p–q), showing that *torch* is essential for quantitative and qualitative differentiation of the gyne phenotype, particularly during the pre-metamorphic growth burst.

Depletion of *torch* in developing gynes thus imposed a coordinated worker-like trajectory by reducing larval adipocytes and lipid droplets and accelerating metamorphosis, culminating in reduced pupal size, shrunken imaginal disc–derived organs, smaller brain neuropils and depleted gyne-specific neuronal populations. These effects closely mirror, in reverse, the enhanced multi-organ growth and delayed metamorphosis that JH treatment can induce in worker-destined larvae ^2^, consistent with Torch being a JH-controlled effector of the gyne specific developmental program.

### JH–Gce–Tai directly activates *torch* to drive caste-differentiated organ growth

The striking mirror image between *torch*-depletion phenotypes in gynes and JH-enlarged worker phenotypes, combined with *torch* being the most JH-responsive gene across the developmental stages (Fig. 1e-f), suggests that *torch* indeed functions as a critical JH-induced effector driving coordinated caste-differentiated organ growth. We therefore investigated whether JH directly activates *torch* transcription through the conserved Gce/Tai receptor complex ^51–53^, maintained in *M. pharaonis* as single-copy *gce* and *tai* genes (Extended Data Fig. 3a; Supplementary Dataset 4). We disrupted Gce and Tai function, both individually and in combination, using splice-blocking vivo-MOs injected into late third instar gyne larvae (Extended Data Fig. 3b-j; Supplementary Dataset 5; see Supplementary Information Text for detailed procedures). Impaired Gce– Tai function consistently reduced *torch* expression by more than 50% in larvae and by more than 90% in prepupae (Fig. 3a; Extended Data Fig. 3k; Supplementary Dataset 6). Combined with JH-induced *torch* activation and the multi-organ phenotypic effects of *torch* knockdown (Fig. 2; Extended Data Fig. 2), these results confirm Torch’s role as JH-controlled effector of caste-differentiated organ growth.

**Fig. 3.**
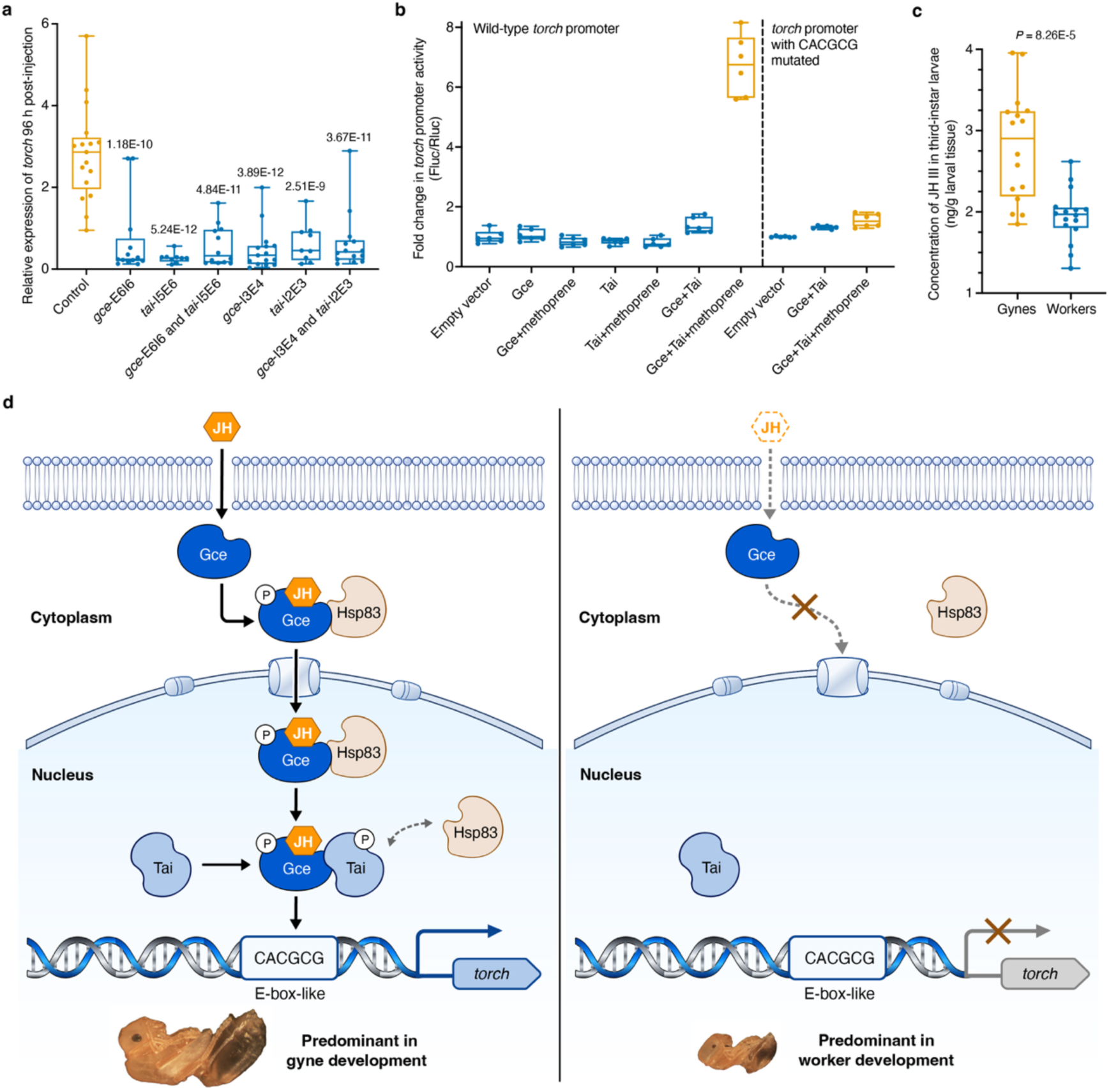
JH–Gce–Tai signaling directly activates *torch* to connect caste-biased JH titers to organ-growth. **(a)** ddPCR in prepupal gynes at 96 h post-injection showed that loss-of-function perturbation of *gce* or *tai*, induced by two independent splice-blocking vivo-MOs per gene (*gce*-I3E4 and *gce*-E6I6; *tai*-I2E3 and *tai*-I5E6), as well as combined *gce* + *tai* perturbation, significantly reduced *torch* transcript abundance relative to controls. These splice alterations are predicted to compromise Gce or Tai protein function. See Supplementary Information Text for vivo-MO validation details. Sample sizes were *n* = 17, 15, 13, 9, 9, 14 and 12 for control, *gce*-I3E4, *gce*-E6I6, *tai*-I2E3, *tai*-I5E6, *gce*-I3E4 + *tai*-I2E3 and *gce*-E6I6 + *tai*-I5E6, respectively. One-way ANOVA: *F*(6, 82) = 20.47, *P* = 1.69E-14. Dunnett-adjusted *P* values for comparisons with the control group are indicated above each perturbation group. **(b)** Dual-luciferase reporter assays in *D. melanogaster* S2 cells. Left panel, methoprene-dependent activation of the wild-type *torch* promoter reporter appears to require co-expression of *M. pharaonis* Gce and Tai. However, mutation of the CACGCG E-box–like motif largely abolishes methoprene-dependent activation by Gce–Tai, consistent with this site being the key JH–responsive receptor in the *torch* promoter. Fold changes were calculated from normalized Fluc/Rluc (Firefly/Renilla luciferase) ratios relative to empty-vector controls. **(c)** JH III concentrations in whole third-instar larvae measured by targeted LC–MS, with gyne larvae showing significantly higher JH III levels than worker larvae, consistent with a gyne-biased endocrine state underlying sustained *torch* expression; *n* = 16 gynes and *n* = 16 workers; two-sided Student’s unpaired *t* = 4.550, df = 30. **(d)** Proposed model (adapted from ^63^) for how *torch* integrates or abolishes JH–Gce–Tai signaling to drive gyne–worker caste differentiation in *M. pharaonis*. In its ligand-bound state, JH promotes formation of phosphorylated transcriptionally active Gce–Tai heterodimers that bind JH response elements and activate target-gene transcription, whereas assembly of this binding complex is strongly reduced in the absence of JH – a regulatory mechanism conserved across diverse insect lineages ^51,52^.

To verify whether *torch* is a direct transcriptional target of the Gce–Tai complex, we mapped *torch* transcription start sites (TSSs) by 5’ rapid amplification of cDNA ends (5’ RACE), which identified two closely spaced TSSs (Extended Data Fig. 3l-m; Supplementary Dataset 7). Inspection of the proximal promoter region (−3 kb to +1 kb) revealed a single E-box–like motif (CACGCG) located ∼650 bp upstream of the TSSs (Extended Data Fig. 3n), a canonical binding site for the JH receptor complex ^51,54–57^. To test whether the ant JH receptor complex indeed activates the *torch* promoter through this motif, we used *Drosophila melanogaster* S2 cells depleted of endogenous JH-receptor components by dsRNAs (Extended Data Fig. 3o-p) ^55–57^. Co-expression of *M. pharaonis* Gce and Tai with a *torch*-promoter luciferase reporter showed that JH analog methoprene increased reporter activity 7-fold over the baseline, but only when both ant factors were present (Fig. 3b, left panel). Furthermore, mutation of the CACGCG motif abolished methoprene-dependent transcription, directly linking Gce–Tai responsiveness to this E-box–like site (Fig. 3b, right panel). These results establish *torch* as a direct transcriptional target of the *M. pharaonis* JH receptor complex and identify CACGCG as the candidate cis-regulatory innovation through which *torch* could be recruited into the conserved JH-dependent transcriptional control system to mediate gyne-worker caste differentiation.

If Torch functions as a direct JH effector, sustained gyne-biased *torch* expression should correlate with the upstream caste difference in JH titers, consistent both with our previous work showing that exogenous JH treatment redirects worker larvae toward gyne-like development ^2,42^ and with reports that elevated larval JH titers influence gyne metamorphosis across multiple social insect lineages ^58–62^. To test this directly, we quantified JH III levels in third-instar larvae, prepupae and early pupae by targeted liquid chromatography–mass spectrometry (LC–MS) and found significantly higher JH III concentrations in gynes than in workers for all three stages (>1.5-fold each; all *P* < 0.001; Fig. 3c; Extended Data Fig. 3q-r). These combined JH-titer data thus provide a parsimonious explanation for the sustained caste-biased *torch* expression observed across the larval-pupal transition in *M. pharaonis* (Fig. 3d).

### Torch accumulates in nuclei of proliferating progenitors

To better understand Torch’s function, we generated a custom rabbit polyclonal antibody and examined its protein expression and subcellular localization. Torch has only one recognizable feature, an N-terminal Sec/SPI signal peptide predicted to be cleaved during maturation (Extended Data Fig. 4a-b). Western blotting indeed detected a single band at ∼18 kDa, matching the predicted molecular mass of mature Torch after signal-peptide cleavage and supporting antibody specificity (Extended Data Fig. 4c). Immunofluorescence revealed prominent nuclear accumulation of Torch across gyne ovaries, wing imaginal discs, fat body and brain (Fig. 4a; Extended Data Fig. 4d-g), the organs that exhibit gyne-specific enlargement ^2^ and whose joint expansion was significantly reduced by *torch* knockdown during gyne development (Fig. 2; Extended Data Fig. 2). This multi-organ nuclear localization at the protein level directly parallels the gyne-biased *torch* mRNA expression in the same organs (Extended Data Fig. 1i and 4h), consistent with Torch acting as a nuclear regulator of organ growth.

**Fig. 4.**
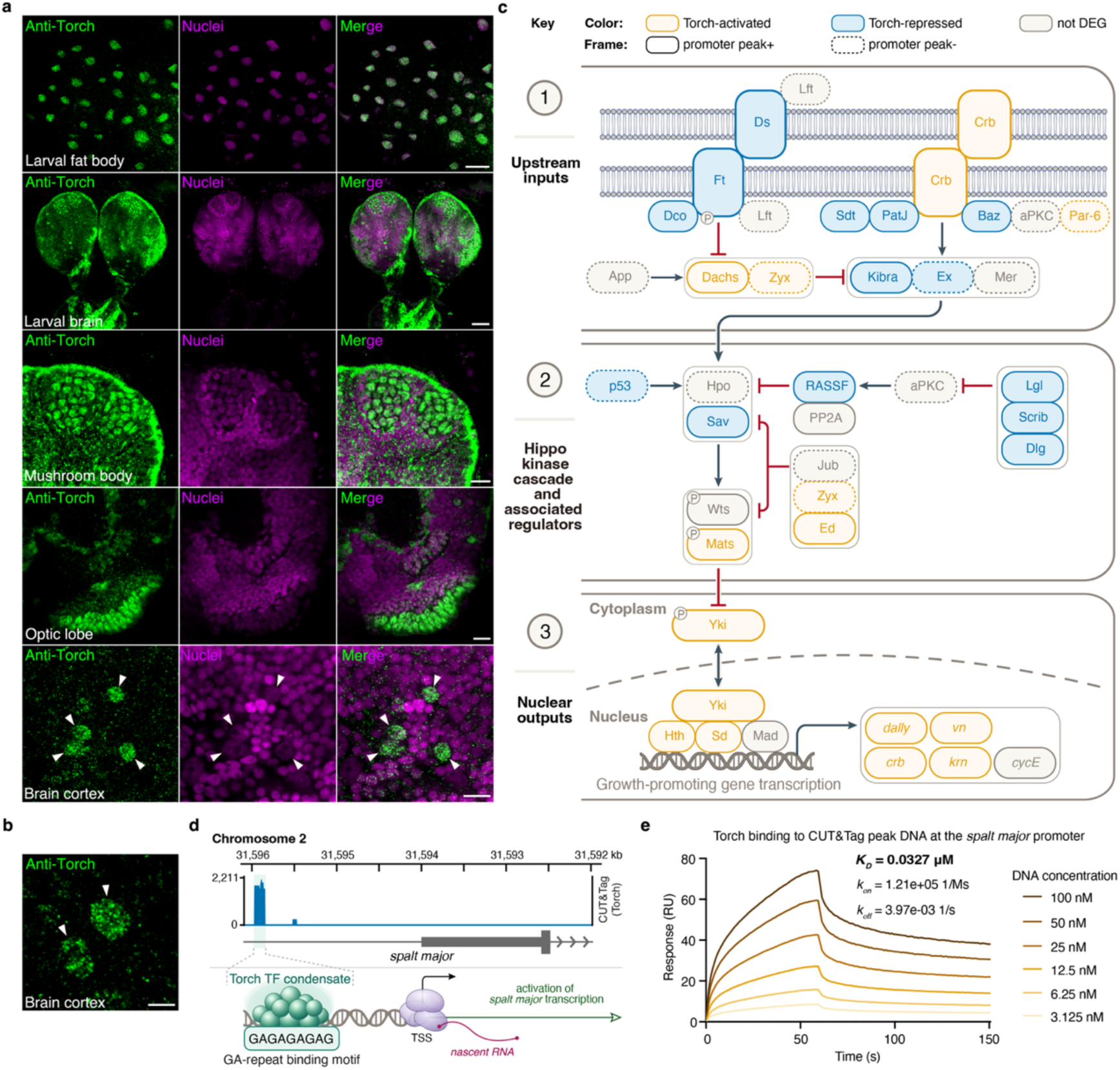
*Torch* connects Hippo-associated transcription to direct activation of *spalt major*. **(a)** Immunofluorescence for Torch (green) in fat bodies and brains of third-instar gyne larvae with counterstained nuclei (magenta). Top row: nuclear enrichment of *torch* in fat bodies. Second row: brain cross-sections showing widespread *torch* expression. Third to fifth row: higher-magnification of mushroom bodies, optic lobes and the superficial brain cortex highlighting strong nuclear accumulation of *torch* in cells with enlarged, weakly DNA-stained nuclei relative to neighboring cells. White arrowheads mark cells with neuroblast-like features and pronounced nuclear Torch abundance. Scale bars, from top to bottom, represent 50, 30, 10, 10, and 10 μm, respectively. **(b)** Higher-magnification view of two neuroblasts (arrowheads) in the superficial brain cortex of third-instar gyne larvae, stained with anti-Torch antibody. Scale bar, 5 μm. **(c)** Schematic of the Hippo signaling pathway in *M. pharaonis*. The pathway was reconstructed from *M. pharaonis* protein-coding genes with identifiable homologs of *Drosophila* Hippo pathway components; only detected Hippo-pathway homologs are shown. The schematic depicts two major upstream inputs, the Ft/Ds and Crb branches, which converge on the Hippo kinase cascade and associated regulators to control Yorkie (Yki)-dependent pro-proliferative transcriptional outputs. Hippo-associated DEGs identified across three developmental stages are integrated into a single pathway-level view. Orange boxes indicate genes positively regulated by Torch which are downregulated DEGs after *torch* knockdown, whereas blue boxes indicate genes negatively regulated by Torch which are upregulated DEGs after *torch* knockdown. Solid boxes denote direct targets with promoter-proximal Torch CUT&Tag peaks, whereas dashed boxes denote indirect targets lacking such peaks. Solid arrows and inhibitory bars indicate regulatory relationships inferred from the conserved *Drosophila* Hippo pathway architecture and do not imply that every interaction has been biochemically validated in *M. pharaonis*. Diagram design adapted from the KEGG fly Hippo signaling pathway map (ko04391) and ^38,126,127^. **(d)** Torch occupancy at a GA-repeat-containing regulatory region upstream of *spalt major*. Upper panel: Genome-browser view of Torch CUT&Tag signal in developing gynes across the *spalt major* promoter region. A prominent promoter-proximal Torch peak on chromosome 2, chr2:31,595,843–31,595,962, is located approximately 1.9 kb upstream of the mapped *spalt major* TSS and contains the 9-bp core GA-repeat sequence GAGAGAGAG. Lower panel: Proposed model in which GA-repeat DNA promotes assembly of a DNA-engaged Torch transcription-factor condensate at the *spalt major* promoter to facilitate transcription. Abbreviations: TF, transcription factor; TSS, transcription start site. **(e)** SPR assay showing Torch binding to the Torch CUT&Tag peak DNA fragment from the *spalt major* promoter, corresponding to the peak shown in panel d. RU, resonance units.

To understand how Torch promotes coordinated organ growth at the cellular level, we focused on the developing brain, which exhibits pronounced caste-specific size differences ^2,64^ and contains well-characterized proliferating progenitors (neuroblasts) whose identity can be validated with established *D. melanogaster* markers ^65,66^. Within developing gyne brains, Torch was strongly enriched in the mushroom bodies, optic lobes and superficial brain cortex (Fig. 4a; Extended Data Fig. 4i), regions known to harbor proliferating neural progenitors in developing *D. melanogaster* brains ^67–69^. Torch-positive nuclei in these regions were consistently enlarged and exhibited weak DNA staining compared with surrounding cells, a cytological signature characteristic of neuroblasts, the neural stem cells that drive brain growth through asymmetric divisions ^70^. These cells also contained prominent DNA-negative nucleoli, consistent with active ribosome biogenesis supporting neuroblast divisions ^71^. To establish the identity of these Torch-expressing cells, we performed triple labeling experiments combining anti-Torch immunofluorescence with RNA-FISH for two independent markers: *Proliferating Cell Nuclear Antigen* (*PCNA*; orthologue LOC105838901), a universal marker of cycling cells ^65^, and *deadpan* (orthologue LOC105832670), a well-established neuroblast marker in *D. melanogaster* ^66^ (Extended Data Fig. 4j; Supplementary Dataset 4). Torch, *PCNA*, and *deadpan* signals showed extensive co-localization predominantly in mushroom bodies, optic lobes, and superficial brain cortex (Extended Data Fig. 4k).

Quantitative analysis revealed that 100% of the Torch-positive nuclei were *PCNA*-positive and 83% were also *deadpan*-positive (*n* = 1,385 Torch-positive nuclei) (Extended Data Fig. 4l), identifying Torch-expressing cells as proliferating neuroblasts. To directly assess proliferative activity, we performed EdU (5-ethynyl-2’-deoxyuridine) incorporation assays, which label cells actively synthesizing DNA during the S-phase^72^. EdU labeling confirmed robust DNA synthesis in Torch-enriched brain regions (Extended Data Fig. 4m) with 60% of the Torch-positive nuclei incorporating EdU (*n* = 1,232 Torch-positive nuclei) (Extended Data Fig. 4l). The lower EdU overlap compared to *PCNA* is expected, as *PCNA* marks cells throughout G1/S/G2 ^65^, whereas EdU specifically labels the S-phase ^72–74^. Together, these data established that Torch accumulates in the nuclei of proliferating progenitor cells in developing gyne brains. Given that *torch*-knockdown significantly reduced neuropil volume (Fig. 2g–h), the accumulation of Torch in proliferating neuroblasts suggests that *torch* expression is required for neuroblast proliferation and the ensuing neuropil expansion that characterizes gyne brains.

We next asked whether this progenitor-associated localization extends beyond the gyne brains. In fat bodies from third-instar gyne larvae, where Torch exhibits similar nuclear enrichment (Fig. 4a; Extended Data Fig. 4f), 100% of Torch-positive nuclei were *PCNA*-positive (*n* = 867 Torch-positive nuclei; Extended Data Fig. 4n-o), identifying that proliferating fat-body progenitors also express Torch. Combined with the reduced adipocyte size and lipid-droplet area observed after *torch* knockdown (Fig. 2a–c), this indicates that Torch promotes fat-body growth by expanding fat-body progenitor number in addition to increasing cell size and lipid storage. Together with the brain neuroblast data, these findings support a model in which Torch acts in proliferating progenitors across developing gyne tissues to coordinate the multi-organ enlargement that defines the developing gyne caste.

### Torch engages Ran and GA-rich regulatory DNA as transcription factor

The strong nuclear accumulation of Torch in proliferating progenitors raised the mechanistic question of how a protein with no recognizable nuclear-localization or chromatin-regulatory domains could gain access to the nucleus and function there. To identify candidate cofactors for Torch nuclear entry and activity, we immunoprecipitated endogenous Torch from gyne-brood lysates and analyzed the associated proteins by qualitative MS across three biological replicates. This showed that Ran, a GTP-binding nuclear protein, was the only reproducibly detectable Torch-associated factor (Extended Data Fig. 5a-b; Supplementary Dataset 8). Independent *in vitro* pull-down assays using the same tissue lysates confirmed the Torch-Ran association (Extended Data Fig. 5c-d; Supplementary Dataset 8), and quantitative MS showed >10-fold Ran enrichment in both Torch co-immunoprecipitated and pull-down samples relative to controls (Extended Data Fig. 5e-f), corroborating the specific interaction between Torch and Ran in *Monomorium* gyne tissues.

We next asked whether Torch directly binds Ran, a deeply conserved small GTPase central to nucleocytoplasmic transport across metazoans ^75^ such that the *M. pharaonis* Ran orthologue shares 91% amino-acid identity with human Ran (Extended Data Fig. 5g). In reciprocal *in vitro* immunoprecipitation assays, Torch formed robust complexes with both *M. pharaonis* and human Ran (Extended Data Fig. 5h-j), indicating direct binding through a conserved Ran interface. Quantitative measurements by surface plasmon resonance (SPR), microscale thermophoresis (MST), and isothermal titration calorimetry (ITC) yielded micromolar dissociation constants (*K_D_* ≈ 1-9 μM) for Torch binding to both *M. pharaonis* and human Ran (Extended Data Fig. 5k-p). Thus, a Hymenoptera-restricted protein has acquired the capacity to engage the deeply conserved Ran GTPase, providing a plausible route by which Torch could connect to nuclear-entry machinery once selection for pre-imaginal caste differentiation became consistent in the ancestral ants.

Having identified Ran as a candidate partner for Torch nuclear access, we next asked how Torch is organized within the nucleus. High-resolution imaging of ant tissue stained with anti-Torch antibody showed that nuclear Torch was not diffusely distributed but instead formed discrete condensates (Fig. 4b; Extended Data Fig. 4f), reminiscent of transcription-factor condensates formed through liquid-liquid phase separation (LLPS) ^76–78^. Purified Torch intrinsically formed dynamic condensates *in vitro*, i.e. assembled into spherical droplets that fused upon contact and exhibited partial fluorescence recovery after photobleaching (FRAP), indicating liquid-like assemblies with internal molecular mobility (Extended Data Fig. 6a-f). Condensates were further disrupted by 1,6-hexanediol, a compound commonly used to perturb LLPS assemblies by weakening hydrophobic interactions, consistent with a phase-separation (Extended Data Fig. 6g-i). We therefore asked whether nuclear Torch directly engages chromatin. CUT&Tag profiling identified 24,089 Torch-binding sites in gyne broods (5,071 genes), with 32% falling within promoter regions (−3 kb/+1 kb of the TSS) (Extended Data Fig. 6j; Supplementary Dataset 9), indicating a broad repertoire of potential direct targets. Despite lower *torch* expression (Fig. 1c), worker broods showed more Torch-bound sites (57,119) but weaker promoter enrichment (∼20%; 7,881 genes) (Extended Data Fig. 6j; Supplementary Dataset 9), suggesting a more distal occupancy landscape that may preserve developmental competence to a lesser extent. This distribution may help explain that organ growth could be rapidly activated by JH to reprogram the less developmentally canalized experimental worker trajectory toward gyne development ^2,42^. By contrast, the gyne landscape contained fewer peaks but stronger promoter enrichment, consistent with a more deeply canalized developmental program ^42^ in which Torch occupies a focused promoter-proximal target set to explicitly stabilize organ-growth programs in gynes.

De novo motif discovery from Torch-bound peaks identified a GA-repeat consensus as the most enriched motif, with strong peak-centered enrichment in both gyne and worker datasets (Supplementary Dataset 9). This motif matched the canonical GAGA element bound by the *D. melanogaster* Trithorax-like protein ^79^ and mammalian ZBTB3/ZBTB7B protein ^80–82^ (Extended Data Fig. 6k; Supplementary Dataset 9). All showed transcription factors recognizing GA-dinucleotide repeats and facilitating transcriptional activation by engaging nucleosomal DNA and recruiting chromatin-remodeling machinery. To validate direct DNA binding, we performed independent biophysical assays using purified Torch protein. SPR, MST, and ITC all demonstrated specific binding to the GAGA oligonucleotide with *K_D_* values of ∼0.5–1.3 µM (Extended Data Fig. 6l-n). Disruption of the GA repeats abolished binding (Extended Data Fig. 6o), confirming sequence-specific recognition. Consistent with the GAGAG sequence being a consensus binding element for the *Drosophila* GAGA factor Trithorax-like ^83,84^, SPR analysis showed that Torch binds to the GAGAG pentanucleotide with a *K_D_* of 81.2 µM (Extended Data Fig. 6p), demonstrating convergent evolution of GA-repeat consensus recognition despite the absence of sequence homology between Torch and Trithorax-like. Together, these results support Torch as a GAGA factor–like pioneer transcription factor that directly initiates caste-biased transcriptional programs during bifurcated preimaginal development.

Because transcription-factor condensates are shaped by protein and DNA partners ^85^, we asked whether Ran and GA-repeat DNA also finetune Torch condensate behavior. GA-repeat DNA indeed enlarged Torch condensates and reduced Torch dynamic mobility within condensates in FRAP assays (Extended Data Fig. 7a-b), consistent with motif DNA stabilizing transcription-factor condensates at cognate regulatory sites to support transcription-factor function ^86^. In contrast, *M. pharaonis* or human Ran reduced Torch condensate size without detectably changing Torch’s dynamic mobility (Extended Data Fig. 7c-f). Thus, GA-rich DNA stabilizes enlarged, chromatin-engaged Torch assemblies, whereas Ran constrains condensate scale, suggesting that Torch activity depends on finetuning condensates within an optimal range for proper transcriptional regulation ^87^. Torch thus engages the deeply conserved Ran machinery and GA-rich DNA as regulators of condensate behavior, explaining molecularly how a newly evolved protein could acquire nuclear regulatory functionality.

Finally, we asked whether Ran binding and GA-repeat recognition are general properties of hymenopteran Torch proteins or specializations refined in ants. *In vitro* pull-down assays using brood lysates from three non-ant Hymenoptera with superorganismal preimaginal caste differentiation, the honeybee *Apis mellifera*, the bumble bee *Bombus terrestris*, and the hornet *Vespa velutina* ^88–90^, recovered Ran from honeybee lysates but not from *B. terrestris* or *V. velutina* lysates (Extended Data Fig. 7g; Supplementary Dataset 10). SPR further showed that only the *A. mellifera* Torch ortholog bound GA-repeat oligonucleotides, albeit with an affinity ∼100 fold lower than *M. pharaonis* Torch; the *B. terrestris* and *V. velutina* orthologs showed no detectable binding (Extended Data Fig. 7h-j). These results indicate that the honey bee Torch ortholog, but not the bumble bee or hornet orthologs, retains a Torch mechanism resembling that of ants with Ran engagement and GA-repeat DNA binding, while the exceptionally high-affinity of GA-repeat recognition in *M. pharaonis* Torch could have been enhanced by adaptive sequence evolution and structural shortening on the ant stem lineage (Extended Data Fig. 1e). Because ants and honey bees independently evolved superorganismal colonies with a perennial lifecycle, this raises the possibility that Torch-based nuclear regulatory mechanisms may have evolved convergently.

### Torch activates organ growth through *spalt major* and Hippo signaling

To define the transcriptional programs through which Torch coordinates organ growth, we performed RNA-seq on individual Torch-depleted and control gynes for late third-instar larvae 24 h post-injection, prepupae 96 h after and early pupae 7 days after larval injection (white pupae lacking visible eye pigmentation; Extended Data Fig. 8a-c; Supplementary Dataset 11). Gene ontology enrichment analysis revealed coordinated disruption of organogenesis programs that mirror the multi-organ size defects caused by *torch* knockdown (Supplementary Dataset 12). In third-instar larvae, downregulated genes were enriched for lipid metabolism and biosynthetic processes (Extended Data Fig. 8d), consistent with impaired adipocyte growth (Fig. 2a-c). Concurrently, *torch* knockdown elicited a precocious metamorphic signature 24 h after injection, because canonical metamorphic markers (*br*, *Eip75B*, and *EcR*) ^91–93^ were among the upregulated differentially expressed genes (DEGs; adjusted *P* < 0.05) (Supplementary Dataset 11). These upregulated genes were enriched for organ-morphogenesis terms (Extended Data Fig. 8d), providing a transcriptomic proxy for the accelerated onset of metamorphosis ^94^ in larvae (Extended Data Fig. 2d-e). Across the prepupal and early pupal stages, downregulated genes were consistently enriched for imaginal-disc and compound eye morphogenesis, wing formation, neuronal and muscle development, neuroblast proliferation, and mitotic cell-cycle programs (Extended Data Fig. 8e-f), consistent with the developmental failure of many caste-differentiated gyne organs following *torch* depletion (Fig. 2d-h; Extended Data Fig. 2f-q).

KEGG pathway analysis identified Hippo signaling pathways as the most consistently enriched across all three developmental stages (Extended Data Fig. 8g; Supplementary Dataset 12). Hippo signaling is one of the most deeply conserved mechanisms for organ size control in the animal kingdom ^37,38^ so we asked which Hippo components are under direct Torch control to promote organ-size expansion towards gyne-typical dimensions. Intersecting stage-specific DEGs after *torch* knockdown (Supplementary Dataset 11) with genes carrying promoter-proximal Torch CUT&Tag occupancy in gyne brood (Supplementary Dataset 9) identified direct Torch-regulated targets spanning multiple nodes in the Hippo pathway (Fig. 4c). Knockdown of *torch* in gyne larvae directly upregulated growth-restrictive Hippo regulators, including *ds*, *ft*, *dco*, *sdt*, *patj*, *baz*, *kibra*, *lgl*, *scrib*, and *dlg*, and directly downregulated the growth-promoting components (*ed* and *hth* in prepupae; *yki, sd, dachs* and downstream pro-growth output genes *dally*, *vn*, *crb*, and *krn* in early pupae). These coordinated transcriptional shifts align with the organ-size reductions in larvae and pupae after *torch* depletion and indicate that Hippo signaling is recurrently engaged by Torch to drive gyne organ-size expansion across the larval–pupal transition.

To pinpoint direct core targets through which Torch connects caste-specific development and JH responsiveness to the growth pathways highlighted above, we applied a stringent four-way integration that retained only genes meeting the following combined criteria: (i) DEGs following *torch* depletion, (ii) caste-biased DEGs between gynes and control workers ^2^, (iii) JH-responsive DEGs between JH-treated and control workers ^2^, and (iv) direct binding by Torch at promoters via CUT&Tag occupancy in gyne broods (Supplementary Fig. 2). This filtering yielded high-confidence direct targets at each developmental stage, enriched for regulators controlling imaginal disc ^95–100^, flight muscle ^101–103^, fat body ^104^, eye ^105–109^ and nervous system development ^103,110–114^, precisely matching the pleiotropic organ-growth defects caused by Torch depletion.

Focusing on prepupal and early pupal stages, the window during which caste-differentiated organs achieve their final form, we asked which direct Torch targets are most likely to drive coordinated organ growth. The *spalt major* gene emerged as the only annotated transcription factor consistently recovered as a direct Torch-activated target in both developmental stages (Supplementary Fig. 2c, e). Expression of *spalt major* was also significantly reduced upon Gce perturbation in prepupae (Extended Data Fig. 8h; Supplementary Dataset 6), consistent with a JH–Gce–Tai→ *torch*→ *spalt major* regulatory cascade. Spalt major is a conserved master regulator that promotes growth and differentiation across the same organs that exhibit caste-specific enlargement in gynes ^115–121^. Spalt major promotes growth by upregulating the Hippo-pathway pro-growth effectors *bantam* in insects ^116,122,123^ and YAP in mammals ^124^, connecting Spalt major to Hippo signaling across metazoans. Together, these analyses suggest that the Hymenoptera-specific pioneer transcription factor Torch engages ancient Hippo signaling at multiple levels, directly regulating Hippo components and downstream targets while integrating into the pre-existing Spalt major–Hippo regulatory network to drive caste-differentiated organ expansion. Spalt major’s wing-promoting function is conserved across diverse insects, including aphids ^125^ and locusts ^116^, and has also been implicated in ant wing polyphenism ^33^, supporting its importance for Torch-mediated organ size control in *Monomorium* gynes.

To validate direct Torch binding to the *spalt major* promoter, we first mapped *spalt major* transcription start sites (TSSs) by 5’-RACE using RNA from gyne prepupae, identifying two TSSs near the NCBI-annotated positions (Extended Data Fig. 8i-j; Supplementary Dataset 7). Torch CUT&Tag then revealed multiple peaks within the *spalt major* promoter in gyne brood, including a prominent peak strongly enriched for the GA-repeat motif (FIMO *P* < 0.0001) (Fig. 4d; Supplementary Dataset 9), consistent with GA-repeat-scaffolded Torch condensate assembly (Extended Data Fig. 7a-b). An electrophoretic mobility-shift assay (EMSA) with recombinant Torch then showed dose-dependent, sequence-specific binding to the *spalt major* promoter fragment (Extended Data Fig. 8k). Finally, quantitative biophysical measurements by SPR, MST, and ITC confirmed high-affinity binding (*K_D_* ≈ 0.01–0.09 µM; Fig. 4e; Extended Data Fig. 8l-m) with no detectable interaction at negative-control fragments (Extended Data Fig. 8n).

### Heterologous *torch* expression promotes latent growth in fruit flies

Because Torch engages the conserved Ran GTPase, the Hippo pathway, and a GA-repeat binding mode reminiscent of *Drosophila* GAGA factor Trithorax-like, its growth-promoting function appears to rely on conserved machinery rather than ant-specific biochemistry. We tested this contention by expressing *torch* in *D. melanogaster*, whose common ancestor with ants lived ∼350 million years ago ^128,129^. Using the UAS– GAL4 system, we expressed *torch* in female tissues with caste-differentiated growth: compound eyes, wings, mushroom bodies, and fat body. The *torch* coding sequence alone failed to efficiently target fruit fly nuclei (Extended Data Fig. 9a) as happens in *M. pharaonis* (Fig. 4a), suggesting that fly importins do not recognize ant nuclear-import signals. However, appending a C-terminal 3× nuclear localization signal (3×NLS; nucleoplasmin–cMyc–cMyc) ^130^ restored strong nuclear localization (Extended Data Fig. 9b), indicating that Torch’s native Ran-mediated import does not fully transfer to *Drosophila*, consistent with cross-species portability concerning chromatin-binding and signaling output, not the import logistics.

Eye-specific expression of *torch-3×NLS* driven by *GMR-GAL4* did not alter eye size in a wild-type *Drosophila* background (Fig. 5a-b), indicating that introducing Torch into flies cannot induce organ overgrowth by itself, consistent with fly organs normally developing to their genetic maximum leaving no room for an exogenous regulator. Ant workers, by contrast, are in a growth-restricted state that Torch alleviates during gyne development. We therefore hypothesized that Torch relaxes growth-restriction, thus being inactive in saturated wild-type tissue but becoming active when Hippo signaling is artificially elevated in a worker-like growth-restricted state. To test this, we co-expressed *torch-3×NLS* with *kibra*, an upstream Hippo pathway activator known to restrict organ growth ^37,131–133^. In this background, *torch-3×NLS* significantly increased eye area (Fig. 5a-b) and head length (Extended Data Fig. 9c) of files compared to *kibra*-alone controls, supporting Torch as a Hippo-restriction de-repressor.

**Fig. 5.**
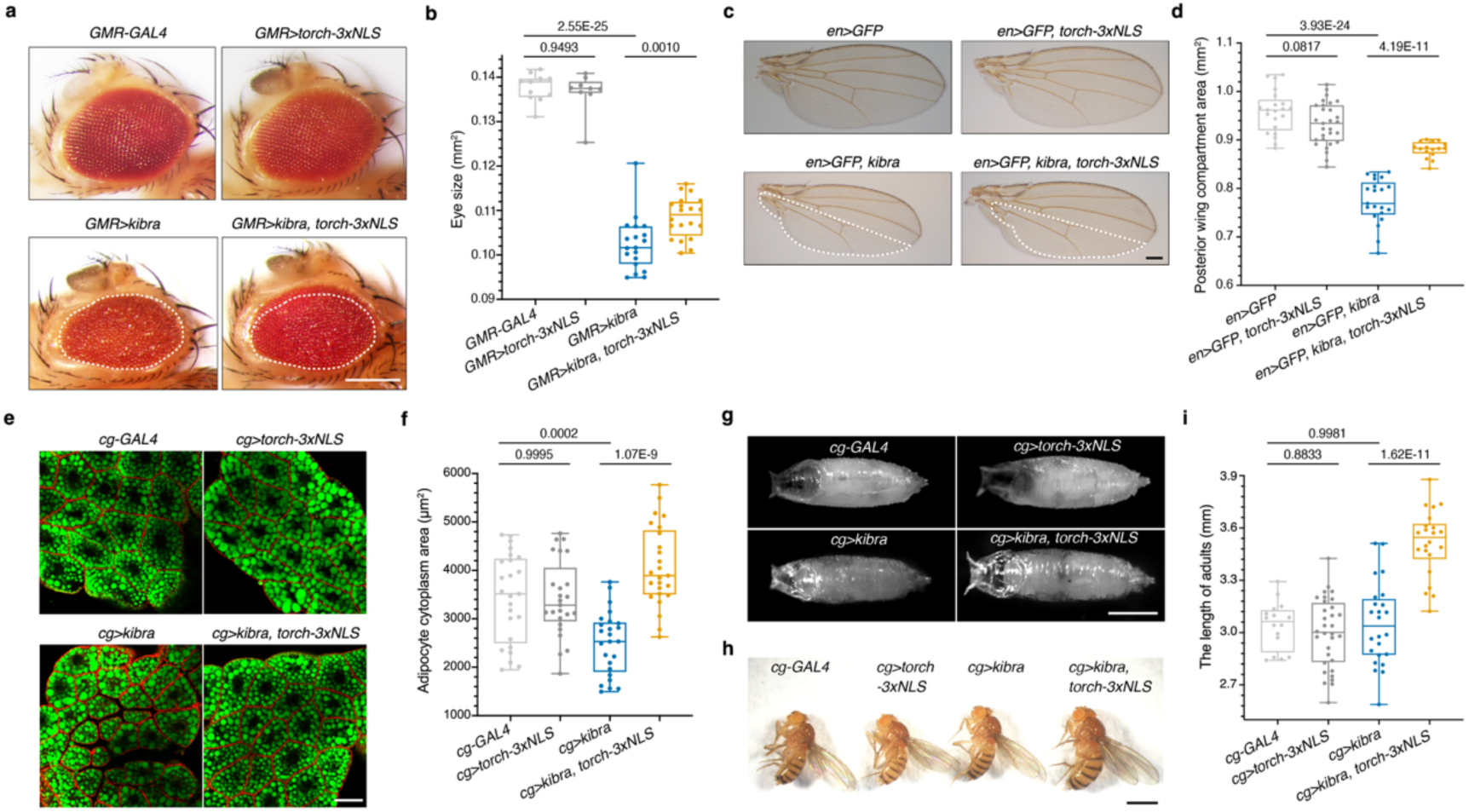
Heterologous Torch promotes organ growth in *D. melanogaster*. (a–b) Eyes (*GMR-GAL4*): Representative adult eyes (a) and eye area (b) – *torch-3×NLS* does not enlarge wild-type eyes but partially rescues *kibra*-induced eye-size reduction under Hippo activated conditions. **(c–d)** Wings (*en-GAL4*): Representative adult wings (c) and posterior wing compartment area (d) – *torch-3×NLS* expression increases posterior wing compartment size only in a *kibra*-expression (Hippo-activated) background. See Extended Data Fig. 9d-e for additional quantifications. White dashed outlines in panels a and c indicate the eye area or posterior wing compartment area of the representative *kibra*-only assay and are copied onto the corresponding *kibra, torch-3×NLS* sample as a visual reference for the partial rescue. **(e–f)** Fat body (*cg-GAL4*): Late non-wandering third-instar larval adipose tissue stained with BODIPY™ 493/503 (neutral lipid droplets; green) and Alexa Fluor™ 555–conjugated phalloidin (F-actin; red) – Representative confocal images (e) and quantification of adipocyte cytoplasmic area (f) show partial rescue of *kibra*-induced adipocyte shrinkage by *torch-3×NLS*. **(g–i)** Body length (*cg-GAL4*): Representative pupae (g), adults (h) and adult body-length measurements (i) showing increased length upon *torch-3×NLS* expression in a *kibra*-expression (Hippo-activated) background. *P* values of significances based on one-way ANOVA followed by Šídák-adjusted planned pairwise comparisons are indicated in the figure. Each data point represents an individual fly in panels b, d, and i, and an individual adipocyte in panel f. Sample sizes follow the left-to-right genotype order in each quantification panel: (b) *n* = 11, 9, 19 and 20; *F*(3, 55) = 188.88, *P* = 6.26E-29; (d) *n* = 21, 29, 21 and 16; *F*(3, 83) = 86.10, *P* = 2.11E-25; (f) *n* = 25 per group; *F*(3, 96) = 16.55, *P* = 9.56E-9; (i) *n* = 16, 32, 24 and 22; *F*(3, 90) = 34.51, *P* = 6.13E-15. Scale bars, 200 μm (a, c), 50 μm (e) and 1 mm (g, h).

A parallel pattern emerged across other tissues enlarged in gynes. Posterior-compartment-specific expression driven by *engrailed-GAL4* (*en-GAL4*) showed that *torch-3×NLS* rescued *kibra*-induced reduction in posterior wing compartment area (Fig. 5c-d; Extended Data Fig. 9d-g). However, the mushroom body experiment with *OK107-GAL4* provided the most rigorous test because neither *torch-3×NLS* nor *kibra* alone perturbed adult α– or β-lobe widths while co-expression significantly enlarged both lobes (Extended Data Fig. 9h-j). This indicates that Torch-driven growth relies specifically on the presence of an active growth-restriction signal and not on Torch’s intrinsic activity in normal tissue. In fat bodies, *cg-GAL4*–driven *kibra* expression reduced adipocyte and lipid-droplet size in late non-wandering third instar fly larvae, but *torch-3×NLS* partially rescued both (Fig. 5e-f; Extended Data Fig. 9k). Strikingly, fat-body co-expression of *torch-3×NLS* and *kibra* also increased body length in late third-instar larvae, pupae and adults relative to single manipulations or to wild-type controls (Fig. 5g-i; Extended Data Fig. 9l-n), suggesting a non-cell-autonomous, systemic growth effect potentially mediated by secreted factors downstream of fat-body Torch activity. This cross-species functionality of Torch, despite the absence of *torch* orthologs outside the Hymenoptera, confirms that Torch’s growth-promoting function is based on conserved rather than ant-specific machinery.

### Heterologous *torch* expression promotes mammalian cell proliferation and organ growth

To test whether Torch’s molecular properties could also be recapitulated in mammalian cells, we expressed 3×NLS-tagged Torch in mammalian cell lines, with the NLS tag used to maximize nuclear access in this heterologous system. In U2OS cells, Torch– GFP–3×NLS formed discrete nuclear condensates that exhibited rapid ∼62% FRAP recovery; half-time ∼6.5 s (Fig. 6a), comparable to *in vitro* reconstitution assays (Extended Data Fig. 6b-i) and to reported transcription factor condensates ^134–137^. Bimolecular fluorescence complementation detected human Ran–Torch–3×NLS association in HEK293T nuclei (Extended Data Fig. 10a), indicating that the Torch– Ran partnership identified in ants could be recovered in this mammalian cellular context. Finally, CUT&Tag profiling of genome-integrated Torch–3×NLS-expressing HEK293T cells, followed by de novo motif discovery, recovered AGAGA among the most enriched motifs within Torch-bound peaks (Extended Data Fig. 10b; Supplementary Dataset 13), consistent with the GA-repeat preference observed in ant chromatin and *in vitro* binding assays (Extended Data Fig. 6k-p). Thus, Torch can exploit mammalian Ran association, dynamic nuclear condensates and GA-repeat chromatin recognition despite deep evolutionary divergence

**Fig. 6.**
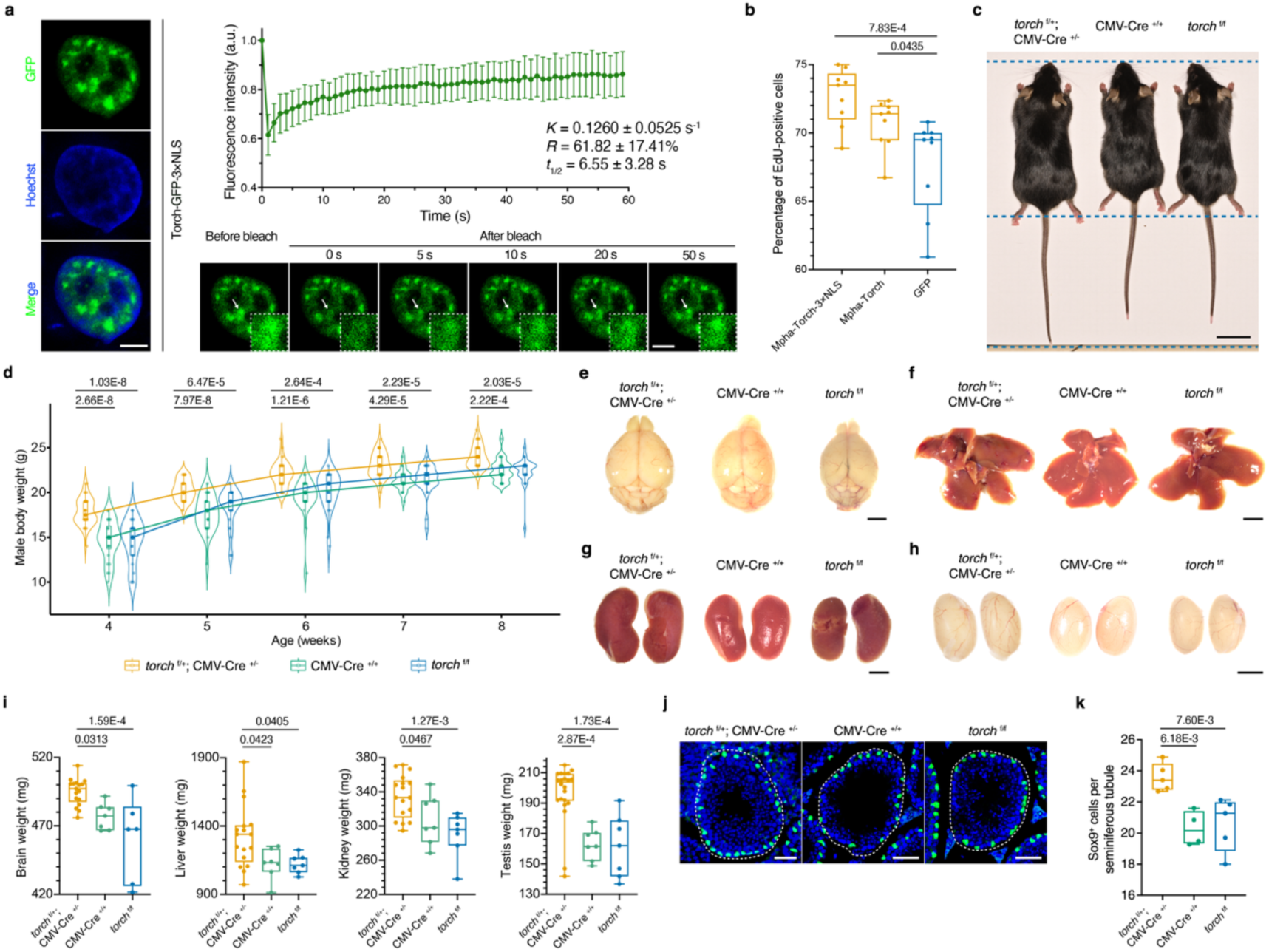
Torch induces mammalian overgrowth phenotypes. **(a)** Representative live-cell images and FRAP analysis of Torch–GFP–3×NLS nuclear condensates in U2OS cells. Left, U2OS cell expressing Torch–GFP–3×NLS with Hoechst-stained nuclei. Top right, corrected fluorescence recovery curve over 60 s after photobleaching after correcting bleached condensates intensities with an unbleached reference condensate in the same field of view and normalizing to the mean pre-bleach intensity. Bottom right, representative time-lapse images before and after photobleaching; arrows mark the bleached condensate, and boxed regions show magnified views of the bleached condensate. Means ± SD; *n* = 6 condensates. **(b)** Flow-cytometric quantification of EdU incorporation in HEK293T cell lines carrying UbC promoter–driven transgenes encoding GFP as a negative control, *M. pharaonis* Torch without an exogenous NLS, or Torch–3×NLS. Mpha, *M. pharaonis*. *n* = 9 independent experiments per group. **(c)** Representative images of two-month-old male mice showing increased overall body size in CMV-Cre;R26-LSL-*torch* mice compared with either of the control genotypes (CMV-Cre only and R26-LSL-*torch* only). Horizontal dashed lines mark the nose-tip, anus and tail-end levels. **(d)** Body weights of male CMV-Cre;R26-LSL-*torch* mice from 4 to 8 weeks of age compared with control genotypes. **(e–h)** Representative images of brain (e), liver (f), kidney (g) and testes (h) from two-month-old male mice, illustrating the larger organ size of CMV-Cre;R26-LSL-*torch* mice relative to both control genotypes. **(i)** Organ weights in two-month-old male mice, showing increased brain, liver, kidney and testes mass in CMV-Cre;R26-LSL-*torch* mice relative to both control genotypes. **(j–k)** Testes from two-month-old males stained for SOX9 to mark Sertoli cells (j) and quantification of SOX9⁺ Sertoli cells per seminiferous tubule (k) in CMV-Cre;R26-LSL-*torch* mice versus controls. In panel j, nuclei are in blue and anti-SOX9 immunostaining marking Sertoli cells in green. Dashed lines indicate seminiferous tubule boundaries. Each point in panels d, i and k represents one mouse; lines in d connect group medians.

We next tested Torch’s growth-promoting activity using genome-integrated HEK293T lines expressing His-tagged Torch, either unmodified or fused to a 3×NLS. Anti-His immunofluorescence confirmed expression: unmodified Torch was distributed between the nucleus and cytoplasm, whereas Torch–3×NLS remained predominantly nuclear with both forming discrete condensates (Extended Data Fig. 10c), consistent with the condensate behaviors observed in U2OS cells, in ant tissues and *in vitro* (Fig. 4b and 6a; Extended Data Fig. 4f and 6b-i). We then quantified DNA synthesis by EdU incorporation using complementary imaging and flow-cytometry assays. Both Torch and Torch–3×NLS increased the fraction of EdU-positive HEK293T cells relative to GFP controls, with enforced nuclear targeting producing the strongest increase (Fig. 6b; Extended Data Fig. 10d–f). Torch expression is thus sufficient to promote S-phase entry in mammalian cells, and this pro-proliferative cellular output is enabled by nuclear access, consistent with the gene’s transcriptional regulatory function in ants.

Mammalian organs develop over an extended postnatal window with substantial inter-individual variation, providing more developmental leeway for detectable perturbed excess growth in a wild-type genetic background than in the tightly canalized fly imaginal discs. To test whether Torch’s growth-promoting outputs extended to mammalian organs *in vivo*, we generated a Rosa26 (R26)-targeted, Cre-dependent *Mus musculus* LSL (loxP-STOP-loxP)-*torch* knock-in line. Crossing this line with a ubiquitous CMV-Cre driver induced systemic Torch expression in mice (Extended Data Fig. 10g; Supplementary Dataset 14). Male CMV-Cre;R26-LSL-*torch* mice exhibited increased overall body size, as reflected by significantly higher body weight than controls (CMV-Cre only and R26-LSL-*torch* only) from one to two months of age (Fig. 6c-d). At two months, these male CMV-Cre;R26-LSL-*torch* mice showed enlargement of brains, livers, kidneys, and testes (Fig. 6e-i), but no significant changes in spleen or heart size (Extended Data Fig. 10h-i). The growth-promoting effect of Torch was sex-specific with female transgenic mice showing no sustained body-weight increase, apart from a transient elevation at one month of age (Extended Data Fig. 10j), and no organ-weight differences at two months (Extended Data Fig. 10k-o). This sex-specificity can likely be explained by sex-specific regulation of mammalian organ growth, including stronger postnatal GH/IGF1 signaling ^138,139^ and sex-biased Hippo-pathway activity ^140–142^ in males. This may provide a more growth-permissive substrate for Torch’s de-repressor activity, although the molecular basis is undermined. The organ-level phenotypes in male mice aligned with Torch’s growth-enhancing outputs in insect brains and fat bodies (functionally analogous to the mammalian liver) (Fig. 2a-c, 2g-h, 5e-f; Extended Data Fig. 9k), suggesting that Torch induces at least in part functionally analogous growth programs across animal Orders.

Activation of Torch expression by germline-specific Vasa-Cre also robustly increased testis mass and testis-to-body weight ratio (Extended Data Fig. 10p-r) in mice, demonstrating tissue-autonomous testicular growth. To connect organ-level phenotypes to cellular outputs, we quantified SOX9^+^ Sertoli support cells per seminiferous tubule within the testis ^143,144^ and found increased Sox9^+^ cell numbers in both CMV-Cre– driven (Fig. 6j-k) and Vasa-Cre driven (Extended Data Fig. 10s-t) *torch*-expressing testes relative to their respective controls, corroborating Torch’s pro-proliferative activity observed in mammalian cell cultures (Fig. 6b; Extended Data Fig. 10d–f).

For panels d, i and k, genotypes are ordered from left to right as CMV-Cre;R26-LSL-*torch*, CMV-Cre-only and R26-LSL-*torch*-only. Sample sizes in this order were: panel d, *n* = 42/24/25, 42/24/28, 41/24/27, 44/22/28 and 41/22/28 at 4–8 weeks, respectively; panel i, *n* = 17/7/6 for brain, 17/7/7 for liver and kidney, and 22/7/7 for testes; panel k, *n* = 5/4/5. Statistical significance in panels b, d, i and k was assessed by one-way ANOVA followed by Dunnett’s multiple-comparisons tests against GFP (b) or the two mouse control genotypes (d, i and k), with adjustment applied separately at each age in panel d; the resulting Dunnett-adjusted *P* values are shown in the corresponding panels. ANOVA results were: panel b, *F*(2, 24) = 8.51, *P* = 1.61E-3; panel d, *F*(2, 88) = 29.70, *P* = 1.39E-10 (4 weeks), *F*(2, 91) = 20.54, *P* = 4.36E-8 (5 weeks), *F(*2, 89) = 16.65, *P* = 7.21E-7 (6 weeks), *F*(2, 91) = 15.34, *P* = 1.81E-6 (7 weeks), and *F*(2, 88) = 14.00, *P* = 5.24E-6 (8 weeks); panel i, *F*(2, 27) = 11.66, *P* = 2.23E-4 (brain), *F*(2, 28) = 4.591, *P* = 0.0189 (liver), *F*(2, 28) = 8.246, *P* = 1.53E-3 (kidney), and *F*(2, 33) = 15.49, *P* = 1.80E-5 (testes); panel k, *F*(2, 11) = 9.178, *P* = 4.52E-3. Scale bars, 5 μm (a), 2 cm (c), 3 mm (e–h) and 50 μm (j).

### Torch engages Hippo-associated growth programs across metazoan lineages

The ability of Torch to promote growth in *Drosophila*, HEK293T cells and mice raised the question of whether these heterologous phenotypes reflect engagement of the same ancient organ-size circuitry that Torch regulates in ant gynes. We therefore profiled transcriptomes across the heterologous contexts in which Torch’s growth-enhancing activity was validated (Supplementary Dataset 14). In *Drosophila*, we compared *GMR-GAL4* female heads co-expressing *kibra* and *torch-3×NLS* with *GMR>kibra* controls, the context in which Torch partially rescued Kibra-induced eye-size restriction. In mammalian cells, we profiled HEK293T lines expressing either nuclear-targeted Torch–3×NLS or unmodified Torch relative to Torch-negative controls. *In vivo*, we profiled brains, kidneys, livers and testes from *torch*-expressing male mice relative to non-expressing controls. KEGG pathway analysis revealed recurrent enrichment of Hippo signaling (*q* < 0.05) in all heterologous datasets except mouse liver and testis, where tissue-specific transcriptional programs may have masked coordinated Hippo-pathway gene-expression changes (Supplementary Dataset 15). Thus, in almost all cases Torch engaged conserved Hippo organ-size control machinery, mirroring the endogenous ant gyne system where Torch depletion perturbed Hippo signaling during caste-specific organ growth (Fig. 4c; Extended Data Fig. 8g).

To identify the direct targets through which Torch engages Hippo signaling outside ants, we generated Torch CUT&Tag profiles in the heterologous contexts where Hippo signaling was recurrently enriched (Supplementary Dataset 13). In each context, genes with promoter-proximal Torch peaks were intersected with matched RNA-seq DEGs (Supplementary Dataset 14), defining direct Torch-regulated genes. The cross-lineage overlap recovered nine conserved direct Torch targets, all of which were consistently downregulated (Supplementary Dataset 14). Among these, *WWC1* is the human ortholog of *Drosophila kibra*. Thus, Torch-mediated repression of *WWC1*/ *kibra* is conserved across these heterologous systems (Extended Data Fig. 10u) and matches the regulatory direction observed in ant gynes, where *kibra* is among the direct Torch-repressed Hippo regulators (Fig. 4c). Moreover, in all tested samples Torch CUT&Tag peaks at the *WWC1*/ *kibra* promoter were consistently enriched for the Torch-binding GA-repeat motif (FIMO *P* < 0.002 in all cases; Supplementary Dataset 13), supporting conserved Torch occupancy at GA-rich regulatory DNA near this growth-restrictive Hippo node.

## Discussion

Our findings reveal how a Hymenoptera-restricted gene, *torch*, became integrated into the ancestral JH pathway to drive coordinated organ growth during pre-imaginal caste differentiation in ants. The mechanism operates both at the cis-regulatory level where *torch* acquired a CACGCG element placing it under direct JH-Gce-Tai control, and at the protein level. Here, Torch partners with Ran GTPase, binds GA-repeat motifs, and modulates Hippo signalling, both by directly regulating pathway components and by activating *spalt major*, a master regulator that itself engages the Hippo size-control machinery. JH’s ancestral anti-metamorphic gatekeeping function ^145^ and its derived multi-organ growth-promoting output ^2^ have thus become unified in a single effector gene *torch*, providing the molecular substrate through which the conserved JH pathway could evolve into an ant caste-differentiation regulator. Torch’s heterologous activity in *Drosophila* (in Hippo-sensitized backgrounds) and mice (in male wild-type background) supports a de-repressor explanation in which Torch releases latent growth potential otherwise held in check by active anti-growth signalling. At the molecular level, Torch engages Hippo-pathway components across ant, fly, mammalian-cell, and mouse models, and the conserved mechanistic embedding of Torch allows accurate predictions of where Torch’s activity should and should not be detectable. Torch is consistently expressed in a gyne-biased manner across ant subfamilies, with the few exceptions almost certainly representing secondary modifications, which supports Torch’s role as key molecular effector of coordinated caste-differentiated organ growth programs associated with the MTE origin to superorganismal colonies with deeply canalized pre-imaginal caste differentiation more than 150 million years ago ^2,42^. That honeybee Torch retains Ran binding and GA-repeat recognition while bumble bee and hornet orthologs do not, raises important questions about the presence or absence of possible convergent mechanisms when the corbiculate bees and vespine wasps realized their independent MTEs to pre-imaginal caste differentiation.

More generally, our results bear on the long-standing question of whether new biological functions arise by novel biochemistry or by novel connectivity that modifies existing pathways. Torch’s functional retention across ∼700 million years of animal evolution ^146^ without the gene being present supports the second scenario. Lineage-specific genes have previously been shown to sometimes acquire developmentally relevant regulatory connections to deeply conserved machinery ^6,7,147–156^, but our study is the first to fully mechanistically dissect this process and to validate its functional relevance across distant lineages. The genetic basis of major phenotypic innovations may thus be more amenable to evolutionary change than previously thought.

## Methods

### Ant colony maintenance and caste and sex identification

Laboratory colonies of *Monomorium pharaonis* of known pedigree were maintained at the University of Copenhagen (Denmark) and the Kunming Institute of Zoology, Chinese Academy of Sciences (China), under previously described conditions ^2^. Colonies were housed at 27 ± 1 °C and 50% relative humidity in Fluon-coated plastic boxes (21 × 17 × 15 cm), with cotton-plugged plastic tubes provided as nesting sites. Colonies were fed twice weekly with frozen house crickets (*Acheta domesticus*) and sucrose agar containing 12.5% sucrose and 1.25% agar. Water was supplied in cotton-plugged tubes and replaced weekly. To obtain reproductive brood, queenless subcolonies were established by transferring workers and mixed-stage brood from stock colonies into Fluon-coated Petri dishes and removing all queens, because reproductive brood is normally suppressed in queenright colonies ^157^. Queenless subcolonies were maintained under the same conditions.

Caste and sex were assigned according to the developmental criteria established for *M. pharaonis* ^50^. From the second larval instar onward, worker-destined and reproductive-destined brood were distinguished by external morphology. Live third-instar reproductive larvae were sexed non-destructively under a bright-field microscope based on genital imaginal-disc morphology ^50^. Larvae identified as gynes or males were reared separately, allowing their sex to be tracked through the prepupal stage. After pupation, gynes and males were distinguished directly from their external morphology. For reproductive larvae collected for dissection, sex was confirmed by examining the gonads: individuals with paired testes were classified as males, whereas individuals without testes were classified as gynes ^50^.

### Comparative and developmental analyses of *torch* expression in ants

Adult whole-body RNA-seq data from reproductive females (gynes or queens) and workers of 68 ant species spanning 46 genera and seven subfamilies were obtained from the published Global Ant Genomics Alliance (GAGA) dataset and the public datasets incorporated therein ^23^; sample collection, RNA extraction, sequencing, species relationships and genome annotations were described previously ^23^. Raw reads were quality-filtered using SOAPnuke v2.1.5 and aligned to the corresponding annotated genomes using STAR v2.7.2b. Gene-level counts and transcripts per million (TPM) were quantified using StringTie v2.1.5 and aggregated by GAGA orthogroup, with abundances summed across member genes for multicopy orthogroups. For GAGA species lacking biological replication, three pseudo-replicates were generated from the raw reads using seqtk v1.4, as described previously ^23^. *torch* orthologues were retrieved from the published OrthoFinder v2.5.4 orthology map across 163 ant genomes, in which *torch* corresponds to hierarchical orthogroup N0.HOG0005048. Seven species lacked a detectable *torch* orthologue, including three of the 68 species with adult caste transcriptomes, leaving 65 species for expression comparisons. Within each species and caste, TPM values were averaged across the available biological replicates or, where applicable, across the three pseudo-replicates generated from a single sample, yielding one species-level mean TPM value for reproductive females and workers. These paired values were compared across all species and within the major ant clades and subfamilies using two-sided paired *t*-tests. Cross-species consistency of caste-biased expression was summarized as the number of species showing a positive reproductive-female-versus-worker log₂ fold change. Species-specific log₂ fold changes and *P* values were estimated from raw orthogroup counts using edgeR v4.0.16 after low-count filtering and TMM normalization, using the model gene count ∼ colony + caste and workers as the reference. *P* values were adjusted using the Benjamini–Hochberg method. *torch* orthologue assignments, log₂ fold changes and adjusted *P* values are provided in Supplementary Dataset 1.

To test whether the degree of reproductive-biased *torch* expression was associated with caste dimorphism, phylogenetic generalized least-squares (PGLS) regression was performed across 27 ant species for which both adult caste transcriptomes and queen– worker dimorphism scores were available; the 27-species phylogeny, including branch lengths, was pruned from the GAGA species tree ^23^. Reproductive bias was quantified for each species as the log_2_ fold change in *torch* expression between the caste-specific mean TPM values of reproductive female (gynes or queens) and workers. The model was fitted by maximum likelihood using the gls function in the R package *nlme*, with phylogenetic covariance specified using a Pagel’s λ correlation structure implemented by corPagel in the R package *ape*.

Published developmental RNA-seq data for *Monomorium pharaonis* and *Acromyrmex echinatior* ^42^ and developmental juvenile hormone (JH)-treatment data for *M. pharaonis* ^2^ were reanalyzed from the original gene-level counts. Counts were normalized and differential expression was analyzed using DESeq2 as described below. For developmental caste comparisons, gynes and workers were compared separately at each stage; for *A. echinatior*, samples from the small, medium and large worker subcastes were treated as a single worker group. DESeq2-normalized torch expression values and adjusted *P* values were used for the developmental profiles in Fig. 1c, d and f. For the JH dataset, *t* statistics comparing gynes with JH-treated workers and control workers were calculated as defined previously ^2^ at five developmental stages. At each stage, *Δt* was calculated as *t*_gyne x JH-treated worker_ − *t*_gyne x control worker_. The mean *Δt* across the five stages and the adjusted *P* value for JH-treated versus control workers at the early pupal stage were used as the x and y variables, respectively, in Fig. 1e.

### Taxonomic distribution and microsynteny of *torch*

The taxonomic distribution of *torch* was assessed from the GAGA orthology dataset spanning ant and outgroup genomes ^23^ and further examined by searching the *M. pharaonis* Torch protein sequence (XP_036149432.1) against NCBI protein and genome databases using BLASTP and TBLASTN, respectively, with Hymenoptera excluded and an *E*-value threshold of 0.05. No matches were detected outside Hymenoptera, supporting the classification of *torch* as Hymenoptera-restricted within current taxonomic sampling. Protein families and conserved domains were queried using InterPro ^158^, and the N-terminal signal peptide and cleavage site were predicted using SignalP 6.0 with the eukaryotic model ^159^. Microsynteny blocks surrounding *torch* across Hymenoptera and Formicidae were inferred using orthogroups computed with SYNPHONI ^160^. Genomes were scanned for collinear neighborhoods anchored by the *torch* orthologue, and conserved clusters were defined by the retention of gene order and composition. Detailed procedures followed the GAGA study ^23^.

### Molecular evolutionary and positive-selection analyses of *torch*

Orthologous *torch* coding sequences were aligned directly at the codon level using PRANK v170427 with the –codon –F parameters ^161^. Corresponding amino-acid sequences were aligned separately using MAFFT L-INS-i v7.525 and filtered with trimAl v1.4.rev22 using the –gappyout option ^162^. A maximum-likelihood gene tree was inferred from the filtered protein alignment using IQ-TREE2 v2.2.2.6 ^163^, with ModelFinder for substitution-model selection and 1,000 ultrafast bootstrap replicates (– m MFP –B 1000). The PRANK codon alignment and inferred gene tree were analyzed in HyPhy; branch-specific *d*N, *d*S and ω (*d*N/*d*S) values were estimated using the standard FitMG94 model in local mode (–type local) ^164^. Episodic diversifying selection on individual branches was tested using the adaptive branch-site random-effects likelihood (aBSREL) model ^165^. Branches were considered to show evidence of positive selection when the nominal (uncorrected) *P* value was <0.01, irrespective of the branch-wide ω estimated using FitMG94, because aBSREL tests whether a proportion of sites on a branch evolved under positive selection even when the overall branch-wide ω is <1.

### *Carebara diversa* ant rearing and sampling

*Carebara diversa* colonies were collected from Qinzhou, Guangxi, China, and maintained at 27 °C and 65% relative humidity under a 12-h light/12-h dark cycle. Colonies were housed in plaster-lined branched test-tube nests supplied with water at the distal ends; the tubes were wrapped in red paper and placed in plaster-lined boxes. Colonies were fed every 2 d with mealworms and a solid agar diet containing 30 g sucrose, 3 g agar and one multivitamin tablet per 250 ml distilled water. Worker and soldier larvae were distinguished by body size, head width and setal morphology. Second-instar workers were smaller than soldiers (mean body length, 680 versus 855 μm; mean head width, 196 versus 213 μm) and bore dense, unbranched setae, whereas soldiers bore dense, curled, bifurcate setae. Third-instar workers were likewise smaller (mean body length, 1,229 versus 1,885 μm; mean head width, 262 versus 283 μm) and bore dense, curled, bifurcate rather than the predominantly three-to five-branched setae of soldiers. Workers and soldiers were sampled at the second instar, on day 1 of the third instar, and at the prepupal and pupal stages. To obtain later third-instar time points, day-1 larvae were transferred to subcolonies and collected 5 or 9 d after the start of the instar. Five biological replicates were collected per caste at each stage, with each replicate comprising a single individual. Individuals were snap-frozen in liquid nitrogen and stored at −80 °C until RNA extraction. Total RNA was extracted separately from each individual using TRIzol Reagent as described below. Library construction, sequencing and read processing were performed as described below, and filtered reads were aligned to the C. diversa ASM4854224v1 reference genome (GenBank accession GCA_048542245.1) and the corresponding gene annotation published in the GAGA study ^23^.

### Methoprene treatment of worker larvae

JH-treated workers used for RNA-FISH and brain immunostaining were generated as described previously ^2^. Briefly, early third-instar worker larvae measuring 0.85–1.15 mm were fed methoprene (5 mg ml⁻¹ in 10% ethanol) on six occasions, whereas control larvae received 10% ethanol alone. Samples were collected at the indicated developmental stages for tissue staining.

### Quantification of *torch* expression abundance by droplet digital PCR (ddPCR)

Total RNA was reverse transcribed with the iScript™ cDNA Synthesis Kit (Bio-Rad, 1708891) following the manufacturer’s instructions. Transcript abundance was quantified by ddPCR with QX200™ ddPCR™ EvaGreen Supermix (Bio-Rad, 1864034) on a QX200 Droplet Digital PCR System (Bio-Rad). Expression values were normalized to *NADH* (*NADH-ubiquinone oxidoreductase subunit 8*), previously validated as a reference gene in ants ^166^. ddPCR primer sequences (5’–3’) were TATTCTACCTTGGTGCACCG (*torch* forward) and CCCACAATAGTGTACGTCAC (*torch* reverse), and AGGTATTTCGAGAACCAGCG (*NADH* forward) and GTCCTTCGAGATCCATCTGC (*NADH* reverse).

### Custom rabbit anti-Torch polyclonal antibody production

A recombinant fragment of *M. pharaonis* Torch comprising residues 25–183 (XP_036149432.1) was cloned into the pGEX-4T-AB1 vector and expressed in *Escherichia coli* Rosetta as a GST– and His-tagged fusion protein following induction with 0.8 mM IPTG for 4 h at 37 °C. The purified recombinant protein was used by ABclonal Technology to immunize a New Zealand rabbit. The rabbit received an initial immunization with 0.3 mg antigen in complete Freund’s adjuvant, followed by four booster immunizations with 0.15 mg antigen in incomplete Freund’s adjuvant. Serum was collected after the final immunization, antibody responses were evaluated by ELISA, and the antiserum was affinity-purified against the recombinant antigen. Antibody specificity was subsequently validated by western blotting of *M. pharaonis* gyne-brood lysates (Extended Data Fig. 4c).

### Western blotting of ant lysates

Two independent biological replicates of whole-body gyne-brood lysates were prepared, each by pooling one larva, one prepupa and one early pupa. Samples were homogenized with a metal bead in 100 μl lysis buffer containing Laemmli sample buffer (Bio-Rad, 1610737), β-mercaptoethanol, Tris-Glycine SDS buffer (Thermo Fisher, 28362) and protease inhibitors using a TissueLyser, heated at 95 °C for 5 min and centrifuged at maximum speed for 5 min. Equal volumes of clarified lysate (20 μl per lane) were separated on Mini-PROTEAN TGX Any kD precast gels (Bio-Rad, 4569033) in 1× Tris/Tricine/SDS running buffer (Bio-Rad, 1610744) at 150 V for approximately 35 min. An Odyssey One-Color Protein Molecular Weight Marker (LI-COR, 928-40000) was included. Proteins were transferred for approximately 7 min to 0.2-μm nitrocellulose membranes using the Trans-Blot Turbo system and transfer packs (Bio-Rad, 1704158). Membranes were blocked for 1 h at room temperature in Intercept (PBS) Blocking Buffer (LI-COR, 927-70001) and incubated overnight at 4 °C with custom rabbit anti-Torch (1:2,000) and mouse anti-β-actin (ABclonal, AC004; 1:2,000) diluted in blocking buffer containing 0.2% Tween 20. After three 15-min washes in PBST (PBS containing 0.1% Tween 20), membranes were incubated for 40 min at room temperature in the dark with IRDye 680RD goat anti-rabbit IgG (LI-COR, 926-68071) and IRDye 800CW goat anti-mouse IgG (LI-COR, 926-32210), each diluted 1:10,000. Membranes were washed three times for 15 min in PBST, rinsed in PBS and imaged using a LI-COR Odyssey imaging system.

### Hybridization Chain Reaction RNA fluorescence *in situ* hybridization (HCR**™** RNA-FISH), HCR**™** Immunofluorescence (HCR**™** IF), and combined assays in ants

Detailed protocols for specimen preparation, staining, imaging, and image analysis are provided in ^2^. Briefly, RNA-FISH probes for all targets in this study (*torch*, *PCNA*, *deadpan*) and all associated HCR™ reagents were obtained from Molecular Instruments, Inc. and used according to the manufacturer’s instructions. Images were captured on a Leica STELLARIS 8 inverted confocal laser scanning microscope, provided by the Center for Advanced Bioimaging (CAB) Denmark at the University of Copenhagen. Primary antibodies were used as follows: rabbit polyclonal anti-Torch (custom; 1:2000 in antibody buffer; overnight at 4 °C) and rabbit polyclonal anti-serotonin (ImmunoStar, 20080; 1:2000 in antibody buffer; overnight at 4 °C). Nuclei were counterstained with SYTOX™ Deep Red (Thermo Fisher, S11381) by adding it to the hairpin solution at 1:1000 and incubating the samples overnight. For the anti-serotonin brain staining, Alexa Fluor™ 546 phalloidin (Thermo Fisher, A22283; 1:400 in antibody buffer) was included as a counterstain and co-incubated with the secondary antibody for 3 hours at room temperature. Where indicated, HCR™ RNA-FISH and HCR™ IF were performed sequentially on the same specimens. Quantification and statistics: the area of medullary serotonergic neurons was measured in Fiji/ImageJ ^167^ from the single focal plane that exhibited the maximal two-dimensional extent of the serotonin signal. Positive cells (anti-Torch, *PCNA*, *deadpan*) were counted manually using the Cell Counter plugin in Fiji/ImageJ from the representative focal plane for each specimen. For each sample, three independent counts were performed and averaged.

### Combined HCR**™** IF and EdU labeling in dissected ant brains

Proliferating (S-phase) cells in late third-instar gyne larval brains were detected with the Click-iT® Plus EdU Imaging Kit (Alexa Fluor® 594 picolyl azide; Thermo Fisher, C10639), following a published protocol ^168^ and the manufacturer’s instructions. Briefly, brains were rapidly dissected in 0.1 M PBS and transferred to an Eppendorf tube containing EdU diluted in Grace’s Insect Medium (Thermo Fisher, 11595030). Tissues were incubated for 1 hour at room temperature to allow sufficient EdU incorporation. Specimens were then fixed and permeabilized as described for HCR™ IF, and the Click-iT® Plus reaction was performed according to the kit protocol. Following EdU detection, HCR™ IF with the anti-Torch antibody was carried out as described above. Positive cells (anti-Torch, EdU) were counted manually from the representative focal plane using the Cell Counter plugin in Fiji/ImageJ ^167^.

### Orthology assessment and phylogenetic analysis

We assessed sequence conservation and inferred phylogenetic relationships for *gce*, *tai*, *PCNA*, and *deadpan* across ants (Formicidae), their closest outgroup Apoidea, and *Drosophila melanogaster*. Taxon sampling, accession lists, tree files and multiple-sequence alignments are provided in Supplementary Dataset 4. Detailed procedural steps followed published protocols ^2^.

### Co-immunoprecipitation coupled to discovery and targeted LC–MS/MS

Whole-body lysates were prepared from *M. pharaonis* gyne broods comprising pooled third-instar larvae, prepupae and early pupae. Three independent biological replicates were performed using an anti-Torch antibody for Co-IP followed by qualitative MS and two replicates for quantitative MS; normal IgG served as the matched negative control. Clarified lysates were incubated with pre-equilibrated agarose beads and anti-Torch (or IgG) overnight at 4 °C, followed by washes and heat elution in SDS buffer for downstream proteomics. Eluates were briefly resolved by SDS-PAGE, gel regions were excised, and proteins were reduced, alkylated, and digested with trypsin; peptides were extracted, dried, and reconstituted for LC-MS/MS.

For discovery proteomics by data-dependent acquisition (DDA) LC-MS/MS, peptides were analyzed on an EASY-nano LC 1000 coupled to a Q Exactive™ HF mass spectrometer using a C18 column (75 µm × 25 cm) at 300 nL min⁻¹ and a 60-min binary gradient (water/0.1% formic acid; acetonitrile/0.1% formic acid). Raw DDA files were processed in PEAKS Studio X+ (Bioinformatics Solutions Inc., Waterloo, Canada) against a *M. pharaonis* protein database (assembly GCF_013373865.1 ASM1337386v2) with trypsin specificity. Search tolerances and modifications are as follows: precursor 7 ppm; fragment 0.02 Da; fixed carbamidomethyl-C; variable Met-oxidation, N/Q-deamidation, protein N-term acetylation. Identification thresholds were set at peptide and protein −10lg*P* ≥ 15, requiring at least one unique peptide per protein. Protein lists from anti-Torch IPs were compared to IgG controls to remove background.

To validate Ran as a putative Torch-interacting protein, targeted prm-PASEF analysis was performed on an UltiMate 3000 LC system coupled to a timsTOF Pro 2 mass spectrometer (Bruker Daltonics). A scheduled assay targeting peptides from Torch and Ran was generated in SpectroDive 11 (Biognosys) from a DDA-derived spectral library. Peptides were separated using a 30-min gradient at 400 nl min^-1^ and acquired with a 100-ms accumulation time, an *m/z* range of 349–1,229 and an ion-mobility range of 0.6–1.6 V s cm^-2^ (1/*K*_0_). Data were analyzed in SpectroDive 11; precursors with *q* ≤ 0.01 were quantified from MS2 fragment-ion chromatograms and manually inspected. Relative enrichment was assessed by comparing two independent anti-Torch Co-IP biological replicates with the IgG control.

### GST-pull down coupled to discovery and targeted LC-MS/MS

A recombinant GST-tagged fragment of protein Torch (amino acids 25-183; accession number XP_036149432.1) was cloned into a modified pGEX-6P-1 vector to generate an N-terminal GST–6xHis fusion. Recombinant GST–Torch was expressed in *E. coli* BL21(DE3) by induction with 0.2 mM IPTG for 6 h at 37 °C. Cells were disrupted by sonication in urea-containing buffer, and GST–Torch was purified sequentially by Ni– NTA affinity, Q-anion-exchange and Superdex 75 size-exclusion chromatography. Protein-containing fractions were pooled, dialysed against 50 mM Tris–HCl (pH 8.0), 300 mM NaCl and 10% glycerol, concentrated, filtered through a 0.45-μm membrane, aliquoted and stored at −80 °C. Protein purity (>95%) was verified by SDS–PAGE.

For the GST pull-down assay, whole-body lysates from *M. pharaonis* gyne broods comprising pooled third-instar larvae, prepupae and early pupae were incubated with GST-Torch immobilized on glutathione-agarose beads. A beads-only sample without GST-Torch was included as a negative control. After extensive washing, bound proteins were eluted from the beads with SDS-PAGE sample buffer. The eluates were resolved by SDS-PAGE, and proteins were in-gel digested with trypsin to generate peptides for liquid chromatography–tandem mass spectrometry (LC-MS/MS) analysis. Both DDA and targeted PRM were performed under the same LC-MS/MS settings and data analysis workflow described in the co-immunoprecipitation (Co-IP) section.

### Reciprocal IPs and Western blot validation of the direct Torch-Ran interaction

A recombinant His-tagged *M. pharaonis* Ran fragment (residues 1-217; accession number XP_012540431.1) was expressed in *E. coli* BL21(DE3) and purified to homogeneity using the same pipeline established for GST-tagged Torch. Recombinant full-length human His-tagged Ran protein (residues 1–216) was purchased from NewEast Biosciences (catalog no. 10112). To assess direct binding between GST-Torch and His-Ran, we performed reciprocal pull-downs (reciprocal IPs) with purified proteins. For the GST-directed assay, GST-Torch was immobilized on glutathione agarose, incubated with equimolar His-Ran, extensively washed, and eluted in SDS sample buffer, following the established GST pull-down workflow. The converse assay captured His-Ran on anti-His resin, incubated with GST-Torch, washed, and eluted. Eluates from both directions were resolved by SDS-PAGE and analyzed by Western blot using anti-GST and anti-His antibodies to detect bait recovery and prey co-precipitation, respectively; input and flow-through fractions were retained where indicated.

### Expression and His pull-down proteomics of non-ant hymenopteran Torch orthologues

Signal-peptide-deleted fragments of *Apis mellifera* Torch (residues 25–396; XP_006557478.1; LOC100577537) and *Bombus terrestris* Torch (residues 25–492; XP_003396730.2; LOC100645125) were cloned into pET-28a and pET-28a-SUMO expression vectors, respectively. Full-length *Vespa velutina* Torch (residues 1–325; XP_047363137.1; LOC124954358) was cloned into pCold TF expression vector. All three His-tagged recombinant proteins were expressed in *E. coli* BL21(DE3) and purified using the same procedure described above for recombinant proteins expressed in *E. coli*. Each recombinant protein was used as bait in His pull-down assays with worker-brood lysates comprising pooled final-instar larvae, prepupae and early pupae from the corresponding species; one bait pull-down and one matched beads-only control lacking His–Torch were performed per species. Bound proteins were trypsin-digested, C18-desalted and analyzed by positive-ion LC–MS/MS on an Orbitrap Exploris 480. Spectra were searched against the NCBI RefSeq proteomes of *A. mellifera* GCF_003254395.2 (Amel_HAv3.1), *B. terrestris* GCF_910591885.1 (iyBomTerr1.2) and *V. velutina* GCF_912470025.1 (iVesVel2.1), and proteins detected exclusively in the corresponding His–Torch pull-down were considered candidate interactors.

### Baculovirus/insect-cell expression of His-tagged Torch

His-tagged Torch recombinant protein was produced by ProteoGenix (Schiltigheim, France) using a baculovirus–insect cell expression system. Briefly, a codon-optimized cDNA encoding Torch (amino acids 25-183; accession number XP_036149432.1) with a C-terminal 6×His tag was synthesized and subcloned into a proprietary baculovirus transfer vector (pTXBac1). Recombinant bacmids were generated in *E. coli* DH10Bac, transfected into *Spodoptera frugiperda* (*Sf*) cells to produce P1 virus, and amplified to P2. Expression screens in *Sf* and *Trichoplusia ni* (*Tn*) cells across multiplicity-of-infection and time courses identified robust production; signal was detected in culture medium and cell extracts by SDS-PAGE and anti-His immunoblot. Optimal expression was achieved in *Tn* cells approximately 72 h post-infection. Conditioned medium from *Tn* cultures grown under these optimized conditions was purified by immobilized metal affinity chromatography via the His tag.

### Biophysical binding assays

Interactions between purified Torch proteins and Ran or DNA were quantified by surface plasmon resonance (SPR), microscale thermophoresis (MST), and isothermal titration calorimetry (ITC). For SPR, proteins were immobilized on a CM5 sensor chip by EDC/NHS amine coupling using a Biacore 8K. Human or *M. pharaonis* Ran was immobilized for the Torch–Ran assays, whereas Torch proteins were immobilized for the DNA-binding assays. The corresponding Torch or DNA analytes were injected over ligand and reference flow cells at the concentrations indicated in the figures and at a flow rate of 30 µl min⁻¹ in PBS-P+ (pH 7.4) containing 5% DMSO. The 150-s measurement comprised approximately 60 s of association and 90 s of dissociation, followed by regeneration with 10 mM glycine–HCl (pH 2.0) for 5 min between concentration cycles. Reference– and solvent-corrected sensorgrams were globally fitted to a 1:1 Langmuir binding model in Biacore Insight to obtain *k_on_*, *k_off_* and dissociation constant *K_D_*, where binding was detected. For MST, Torch was labeled with RED-NHS dye (NanoTemper) for 30 min at room temperature, and excess dye was removed by desalting into 50 mM HEPES (pH 7.4) containing 0.05% Tween-20. Sixteen two-fold dilutions of Ran or DNA were mixed 1:1 with labeled Torch, incubated for 20 min at room temperature, and loaded into Monolith NT.115 capillaries (NanoTemper) for thermophoresis. *K_D_* values were calculated using NanoTemper analysis software. For ITC, samples were degassed for 10 min and analyzed using a Nano ITC microcalorimeter (TA Instruments). Human or *M. pharaonis* Ran was loaded into the sample cell and Torch into the injection syringe; for DNA measurements, Torch and DNA were loaded into the cell and syringe, respectively. Twenty injections separated by 200 s were performed at 25 °C, and binding isotherms were fitted in NanoAnalyze to determine *K_D_*.

### In vitro condensate formation, fluorescence recovery after photobleaching (FRAP) and turbidity assays

Purified proteins were diluted to the indicated concentrations in reaction buffer containing 20 mM HEPES (pH 7.5) and 150 mM NaCl. Except for matched no-PEG controls included only in the concentration-series imaging, all condensate assays contained 10% PEG 8000. *M. pharaonis* Torch–AF488 was tested at 10, 20, 40, 80 and 160 μM in glass-bottom 384-well plates after incubation at 4 °C overnight. The PEG-containing samples were used to quantify condensate number and area fraction (total condensate area divided by field area) from five independent fields per concentration using Analyze Particles in Fiji/ImageJ. Fusion of condensates formed from 160 μM Torch–AF488 was monitored by time-lapse imaging after 0.5–1 h incubation. For 1,6-hexanediol treatment, condensates formed from 80 μM Torch–AF488 over 0.5–1 h were treated with 0 or 10% 1,6-hexanediol for 20 min before imaging and quantification from five independent fields per condition.

To assess the effects of DNA and Ran, 40 μM Torch–AF488 was combined with 0.2, 1.0 or 2.0 μM Cy3-labelled GA-repeat DNA, with DNA-only samples at the same concentrations included as controls. Separately, 80 μM Torch–AF488 was combined with 5.5, 11 or 22 μM AF594-labelled *M. pharaonis* or human Ran. Samples were incubated at 4 °C overnight before imaging. For turbidity assays, absorbance at 600 nm was measured for 80 μM Torch alone or combined with 22 μM *M. pharaonis* or human Ran. Fluorescence and differential interference contrast images were acquired using a Zeiss LSM 980 Airyscan 2 confocal microscope equipped with a 63×/1.4-NA oil-immersion objective. FRAP was performed after 0.5–1 h incubation on condensates containing 160 μM Torch alone, 40 μM Torch plus 2 μM GA-repeat DNA, or 80 μM Torch plus 22 μM *M. pharaonis* Ran. A region within each condensate was photobleached, and fluorescence was recorded every 5 s for 240 s. Fluorescence intensities were background-subtracted, corrected using an unbleached reference condensate and normalized to the mean pre-bleach intensity. For co-condensates, recovery was quantified separately for each fluorescent component. Recovery curves, half-times and fractional recoveries were obtained as described below for U2OS-cell FRAP.

### Vivo-MO synthesis and larval microinjection

Vivo-morpholino antisense oligonucleotides (Vivo-MOs) targeting *torch*, *gce*, and *tai*, along with negative-control Vivo-MOs, were designed and synthesized by Gene Tools, LLC. The Vivo-MO sequences (5’–3’) and their sense-strand pre-mRNA target splice-junction sequences (the bracketed region denotes the binding site) were listed below; uppercase letters denote exonic sequences, whereas lowercase letters denote intronic sequences: *torch*-E2I2, ATCTTTCCTTGACTTACCTGACTTT, CGAATTTAGATTTTTGT[AAAGTCAGgtaagtcaaggaaagat]aaaacatt; *torch*-I3E4, ATCTGCAAAATGGAACAAACATCTT, aa[aagatgtttgttccattttgcagAT]TACCATTTATTCTACCTTGGTGC; *gce*-I3E4, TTCCAACTGCTGTAAGTAATGCACA, tacttatct[tgtgcattacttacagCAGTTGGAA]GTGGCACGAAGAATTT; *gce*-E6I6, CGTACCGAAAGATGAAGGCTCACCT, AACGGGAAGAGAGAAAGGCAACG[AGgtgagccttcatctttcggtacg]tc; *tai*-I2E3, GCACTTGTTACTGCAAACAATTGAA, aattttacct[ttcaattgtttgcagTAACAAGTGC]CTTAACGAAAAAAGA; and *tai*-I5E6, CAGCCTGGAATATGCAAACTCAAGT, aaat[acttgagtttgcatattccagGCTG]GACTAGCGAGCCACAGCCTCA. The commercially available negative-control Vivo-MO (CCTCTTACCTCAGTTACAATTTATA) targets a human β-globin intron mutation associated with β-thalassaemia.

Oligonucleotides were ordered at the 400-nmol scale and resuspended in 800 μL RNase-free water to yield 0.5 mM stock solutions. For each of *torch*, *gce* and *tai*, two independent vivo-MOs targeting distinct exon–intron junctions (see Supplementary Information Text) were injected into late third-instar *M. pharaonis* gyne larvae measuring 2.1–2.3 mm in body length, a critical window for caste-specific organ differentiation ^2^. Microinjection needles were pulled from filamented borosilicate-glass capillaries (WPI, 1B100F-4) using a P-97 Flaming/Brown micropipette puller (Sutter Instrument) and mounted on a Narishige IM-400 microinjector. Injections were performed under an Olympus SZX9 stereomicroscope equipped with a custom-built x– y translation stage. Each larva received approximately 0.05 µl of the 0.5 mM vivo-MO stock solution, with matched negative-control injections performed in parallel for each experimental batch. Successful delivery was confirmed by visible transient distension of the larval body cavity. Following injection, larvae were examined for entry into the prepupal stage at 12-h intervals from 24 to 96 h post-injection, with prepupae identified as described previously ^50^. For each of five independent experiments per treatment, the cumulative proportion of prepupae was determined at each time point, and T50—the time at which 50% of larvae had entered the prepupal stage—was estimated by linear interpolation between adjacent observation times.

### Validation of Vivo-MO-mediated splice disruption

At 24 h post-injection, total RNA was extracted and reverse-transcribed into cDNA. Vivo-MO-induced splicing changes in *gce* and *tai* were assessed by PCR using gene-specific primer pairs positioned outside the targeted exons (*gce*: GGTCACAGCAGTCACGGTAA and TTATAGCGCCCGGAGACAAC; *tai*: GAATTTGCGGGTGGAGGAGA and CATTTTCGCTGCCGCATACA), together with exon-skipping-specific primer pairs for *gce*-I3E4 (ACGTACTACGCGAAACCACG and ACCGAAATATTGCTCCCCTTGT), *gce*-E6I6 (CATTAGTCGACAAGGGGAGCA and CATCCTTGTGCATGAAGGCG), *tai*-I2E3 (CGTGTGAATTGCAAGACCCG and CGTTACTGGGCAGAAGAGCA), and *tai*-I5E6 (GGAGAAGTGTCCTCGTCGAA and TCCGATTTGGACCGTATCGTT). All primer sequences are shown in the 5’–3’ orientation. PCR products corresponding to the wild-type and Vivo-MO-induced exon-skipped transcripts were excised from agarose gels, purified using the NucleoSpin Gel and PCR Clean-up Mini kit (MACHEREY-NAGEL; 740609.50), and subjected to Sanger sequencing.

### Morphometric measurements in *torch*-knockdown and control gyne pupae

Live pupae and dissected reproductive organs were imaged on a Leica MZ12.5 stereomicroscope equipped with a Leica DFC420 camera and companion software (Leica, Germany). Reproductive organs from 12-day-old pupae were dissected in PBS and mounted in a glass depression slide with a coverslip. Z-stacks were captured at multiple focal planes and combined using Zerene Stacker v1.04 (Zerene Systems, Richland, WA, USA). To assess developmental effects of *torch* knockdown, the following traits were quantified from captured images: body length (sum of mandible, head, mesosoma, petiole, postpetiole, and gaster lengths) from lateral views; wing length, mesosoma area, and compound eye area from lateral views; head width (across the eyes) and ocelli area from dorsal views; and antennal scape length from an oblique view that maximized scape projection. Spermatheca area was measured from dissected preparations. All lengths and areas were measured in Fiji/ImageJ ^167^.

### Quantification of lipid droplets and cytoplasmic area in gyne adipocytes

Late third-instar gyne larvae were collected 24 h post-injection from size-matched control gynes and *torch*-knockdown gynes. For each condition, fat bodies from three larvae were dissected, pooled before staining. Adipocytes were stained following a published protocol ^2^: lipid droplets with HCS LipidTOX™ Red Neutral Lipid Stain (1:1000 in PBSTw; H34476, Thermo Fisher) and F-actin to delineate cell boundaries with Alexa Fluor™ 546 Phalloidin (1:400 in PBSTw; A22283, Thermo Fisher). Specimens were imaged under identical acquisition settings across conditions. Image analysis followed the same published workflow ^2^, whereby LipidTOX-positive lipid droplets and phalloidin-defined cell areas were segmented using uniform parameters, and per-cell lipid-droplet and cytoplasmic areas were extracted for downstream statistical comparison.

### Volume quantification of gyne brain neuropils

Heads from 12-day-old (final day) pupae of control gynes and *torch*-knockdown gynes were processed for whole-mount preparation. Fixation, bleaching, confocal microscopy, and three-dimensional reconstruction in AMIRA v2021.2 (Mercury Computer Systems, Germany) followed a published protocol ^2^. For each condition, ten brains (total *n* = 20) were imaged under identical acquisition settings and reconstructed in parallel to minimize batch effects. Neuropil volumes were segmented in AMIRA and computed from voxel counts.

### Targeted LC–MS/MS quantification of juvenile hormone III

Juvenile hormone III (JH III) was quantified in whole-body *Monomorium pharaonis* gyne and worker broods at the third-instar larval, prepupal and early pupal stages by targeted LC–ESI–MS/MS using multiple-reaction monitoring, with modifications from established JH-detection methods ^169^. For each biological replicate, 10 mg of pooled brood material was homogenized for 5 min in 50 µl of 50% aqueous methanol, followed by addition of 150 µl acetonitrile. Samples were sonicated for 10 min, incubated at −20 °C for 1 h and centrifuged at 13,000 rpm for 10 min. JH III was analyzed on a Shimadzu LC-30AD UPLC system coupled to a SCIEX QTRAP 6500 mass spectrometer using a CORTECS T3 column (1.6 µm, 2.1 × 150 mm; Waters, 186008500). Multiple-reaction monitoring was performed in positive-ion mode using *m/z* 267.1 → 189.1 for quantification and 267.1 → 147.1 and 267.1 → 107.1 as qualifier transitions. JH III was quantified against an external calibration curve generated from an analytical standard (GLPBio, GC43934); concentrations were corrected using the spike recovery and normalized to sample wet mass.

### 5’ RACE mapping of transcription-start sites (TSSs) for *torch* and *spalt major*

The 5’ ends of *torch* and *spalt major* transcripts were characterized by 5’ rapid amplification of cDNA ends (5’ RACE) using whole-body RNA from third-instar gyne larvae and gyne prepupae, respectively. The known internal regions were first amplified using GTATTGACGAGCAAGATGAAC and CCTGAAGCAGATTCGGCGT for *spalt major*, and ATGTGGAGCGCTTTATTGAAATCGT and AAAAACTTAGAAAAGTCATATAACGGTTC for *torch*. First-strand cDNA was synthesized using the gene-specific primers CGGTGACGTTTGCTGATGATG and TCCGTCGTTGACGATTGCAT for *spalt major*, and AACTATGTTATGTGCAATCGGGTAT and TGGAGAAATCTTCATCGGTGC for *torch*. Following RNase H treatment and TdT-mediated C-tailing using a 5’-RACE kit (Sangon Biotech, B605102), two rounds of touchdown nested PCR were performed using TTGATTCTGAAGGGTGCACAG followed by AGAAATGCCAGGTCTTGTAGC for *spalt major*, and TCGGTGGTATGTAAACACATTCATA followed by CTGATGTAGCTTTGTCATTTATGGAT for *torch*, together with the kit-provided adapter primers. Amplicons were gel-purified and cloned into pMD18-T. The cloned 5’-RACE products were subjected to Sanger sequencing using universal M13 primers flanking the pMD18-T insertion site. The resulting reads were assembled to define the 5’ UTRs, and the most upstream nucleotide supported by the cloned 5’-RACE sequences was designated as the candidate TSS for each gene. All primer sequences are shown in the 5’–3’ orientation.

### Electrophoretic mobility shift assays (EMSA)

EMSA were performed with baculovirus/insect-cell-expressed Torch recombinant protein and 5’-FAM-labeled DNA probes from the *spalt major* promoter (chr2:31,595,843–31,595,962; NC_050468.1). Probes were generated by PCR with a 5’-FAM forward primer, verified on 2% agarose, and purified with kit (B110093, Sangon Biotech). EMSA reactions were performed in a total volume of 20 μl containing 50 ng FAM-labelled DNA probe, 2 μg salmon sperm DNA, 0–10 μg purified Torch protein and 1x binding buffer comprising 10 mM Tris-HCl (pH 7.5), 50 mM KCl and 1 mM DTT. Competition reactions contained a 100-fold molar excess of unlabeled cognate or negative-control DNA. Reactions were incubated at 25°C for 30 min. Samples were electrophoresed on 6–8% native PAGE at 60 V for 30 min then 170 V for 30 min at low temperature, and bands were visualized on a fluorescence gel imager (Furi Technology).

### Generation of UAS transgenic *Drosophila melanogaster* lines carrying *torch* or *torch-3×NLS* insertions

UAS transgenic *Drosophila melanogaster* lines were generated by Fungene Biotechnology Co., Ltd. A full-length coding sequence of *torch* (residues 1-183; accession number XP_036149432.1), either untagged or fused C-terminally to a 3×nuclear-localization signal (NLS; nucleoplasmin-cMyc-cMyc), was codon-optimized for *D. melanogaster* and synthesized. The 3×NLS cassette comprised one nucleoplasmin NLS and two c-Myc NLS motifs, both widely used to drive robust nuclear import in metazoan cells ^130^. The inserts were cloned into a UAS-attB donor plasmid to permit GAL4-dependent expression after site-specific integration. Constructs were integrated by φC31 recombination at attP40 (2L, 25C6) or attP2 (3L, 68A4) ^170^; surviving P0 adults were crossed to w1118, and F1 transformants were identified by eye-color or 3×P3-RFP markers, balanced and verified by junction PCR. For experiments, UAS lines were crossed to the indicated GAL4 drivers. Stocks used were *w*^1118^ (Bloomington Drosophila Stock Center, 5905); *GMR-Gal4^S^* and *en-Gal4* (gifts from Lei Xue, Tongji University); *UAS-kibra* (gift from Duojia Pan, University of Texas Southwestern Medical Center) ^171^; *cg-Gal4, UAS-GFP* (gift from Chenxi Wu, North China University of Science and Technology) ^172^; *OK107-Gal4* (gift from Yongfeng Jin, Zhejiang University) ^173^; and the custom *UAS-torch* and *UAS-torch-3×NLS* lines generated by Fungene Biotechnology.

### *Drosophila* husbandry and morphometric analyses

*Drosophila melanogaster* stocks and crosses were maintained on standard cornmeal– yeast food at 25 °C under a 12-h light–12-h dark cycle, as described previously ^174^. The indicated UAS lines were crossed to the corresponding GAL4 drivers. All late third-instar larvae, late pupae and adults analyzed in this study were female. Adult heads, dissected wings, and intact larvae, pupae and adults were imaged in bright-field mode using an Olympus SZX16 stereomicroscope and cellSens Dimension software (Olympus). Morphometric measurements were obtained from calibrated images using Fiji/ImageJ.

### *Drosophila melanogaster* tissue immunofluorescence

Female late non-wandering third-instar larvae were immobilized on dissection pads in PBS, and wing imaginal discs and fat bodies were dissected using fine forceps. For each fat-body staining preparation, fat bodies from three larvae were pooled after dissection and processed together. Virgin adult females were immobilized similarly, and brains were dissected in PBS. Tissues were fixed in 4% paraformaldehyde for 15 min at room temperature, washed three times for 20 min in 0.1% PBSTx, and blocked in 5% donkey serum for 1 h. Adult mushroom bodies were labeled overnight at 4 °C with mouse anti-Fasciclin II (1D4, DSHB; 1:200), whereas larval fat bodies were labeled under the same conditions using the same custom rabbit anti-Torch antibody used for the ant samples (1:1000). After three washes with 0.1% PBSTx, samples were incubated for 2 h at room temperature in the dark with Alexa Fluor Plus 555-conjugated anti-mouse (Invitrogen, A32727; 1:400) or anti-rabbit (Invitrogen, A32732; 1:400) secondary antibody, as appropriate. For fat-body analyses, F-actin was labeled with Alexa Fluor 488– conjugated phalloidin (Cell Signaling Technology, 8878S; 1:800) for 15 min at room temperature. In separate neutral-lipid staining experiments, lipid droplets were labeled with BODIPY 493/503 (Thermo Fisher Scientific, D3922; 1:1000) together with Alexa Fluor 555–conjugated phalloidin (Cell Signaling Technology, 8953S; 1:800) for 15 min at room temperature. Nuclei were counterstained with DAPI. For larval wing discs, native GFP fluorescence was imaged directly in the GFP-marked *en-Gal4* genetic background. Samples were mounted VECTASHIELD Vibrance Antifade Mounting Medium with DAPI (Vector Laboratories, H-1800), and slides were stored at 4 °C until imaging. Images were acquired using a Zeiss LSM 980 inverted confocal laser-scanning microscope.

Microscopy-based measurements were performed in FIJI/ImageJ using identical acquisition settings and analysis parameters across experimental conditions. Fat-body adipocyte lipid-droplet and cytoplasmic areas were quantified from the confocal optical section showing the largest phalloidin-defined cytoplasmic area for each adipocyte, following our previously published protocol ^2^, whereas wing-disc GFP-positive area and mushroom-body lobe width were measured from maximum-intensity projections of the corresponding confocal z-stacks.

### dsRNA-mediated depletion and RT–qPCR validation in S2 cells

*Drosophila melanogaster* S2 cells (FBtc9000001), provided by the laboratory of Xianjue Ma (Westlake University), were cultured and transfected as described previously ^175^. DNA templates for double-stranded RNA (dsRNA) synthesis were amplified using gene-specific primers containing 5’ T7 promoter sequences. dsRNAs targeting endogenous *D. melanogaster gce* and *tai*, together with a dsRNA targeting GFP served as the negative control, were synthesized and purified using the MEGAscript RNAi Kit (Thermo Fisher, AM1626) according to the manufacturer’s instructions. To reduce endogenous JH-receptor activity, S2 cells were seeded in 12-well plates in 1 ml medium and simultaneously transfected with *gce*– and *tai*-targeting dsRNAs at a combined total of 20 μg per well using Effectene Transfection Reagent (QIAGEN, 301427). Cells were incubated for 24 h before reporter-plasmid transfection. Control cells were treated in parallel with 20 μg GFP-targeting dsRNA and processed on the same schedule. Owing to sequence similarity between *gce* and *met*, the gce-targeting dsRNA also reduced endogenous *met* expression. For the specificity analysis, S2 cells were transfected with pUASTattB *Drosophila* gene expression vectors encoding codon-optimized, full-length *Monomorium pharaonis* Gce (XP_036139069.1) and Tai (XP_036149276.1) proteins. Depletion of endogenous *D. melanogaster met*, *gce* and *tai*, and the absence of substantial effects of the dsRNAs on the expression of co-transfected *M. pharaonis gce* and *tai*, were verified by RT–qPCR.

Total RNA was extracted using TRIzol Reagent (Thermo Fisher Scientific, 15596018) and reverse-transcribed using the HiScript II 1st Strand cDNA Synthesis Kit (Vazyme, R211-01). Quantitative PCR was performed using Taq Pro Universal SYBR qPCR Master Mix (Vazyme, Q712-02) on a qTOWER384G Real-Time PCR System (Analytik Jena). Transcript abundance was normalized to *rp49*. Primer sequences are given in the 5’–3’ orientation. The dsRNA-template primers were *D. melanogaster gce*, forward TAATACGACTCACTATAGGGAGAAACTCCTCCACGAGCCACAGT and reverse TAATACGACTCACTATAGGGAGAAGAGGGCGTCGGCGATTGGTA; *tai*, forward TAATACGACTCACTATAGGGAGAAGCCCAAGTGACTTACCAGAT and reverse TAATACGACTCACTATAGGGAGAGCTGGTGCTGGTAGGCCAATT; and GFP, forward TAATACGACTCACTATAGGGAGAATGGTGAGCAAGGGCGAGGAGCTG and reverse TAATACGACTCACTATAGGGAGACTTGTACAGCTCGTCCATGCCGAGAG.

The RT–qPCR primers were: *D. melanogaster met*, forward GACGATGACGAGGATCACCC and reverse TTCTCGAGCGTCGTATCAGC; *gce*, forward TGTTCTGGGCAAGAGCGAAT and reverse CCGAGCTCTCGTTCTCATCC; *tai*, forward TGCGAGTGAGTATTGGGGAA and reverse CAATGTTGTTCAAGCCCGCA; *M. pharaonis gce*, forward CGATGATCGGTCTACGGCAA and reverse TTTCATCAGGCGGATGCCTT; *M. pharaonis tai*, forward TGAACCAGCAGGACGAAGAC and reverse GACTCCGCAAGAGAGGATCG; and *D. melanogaster rp49*, forward CCACCAGTCGGATCGATATGC and reverse CTCTTGAGAACGCAGGCGACC.

### Dual-luciferase reporter assays in S2 cells

A 4,000-bp genomic fragment spanning the −3-kb to +1-kb promoter region of *torch* (NC_050477.1:16,120,736–16,124,735) was synthesized and cloned immediately upstream of the firefly luciferase gene in pGL3-Basic vector. This fragment encompassed the two experimentally mapped *torch* transcription start sites at positions 16,123,750 and 16,123,782 and contained a single E-box-like motif (CACGCG) at positions 16,123,099–16,123,104. The mutant reporter was identical to the wild-type reporter except that CACGCG was replaced with GTGCGC. Following dsRNA pretreatment, cells in each well were co-transfected with 20 ng of the wild-type or mutant *torch* firefly luciferase reporter, 10 ng of a copia–Renilla luciferase internal-control plasmid, 5 ng of the Act5C–Gal4 driver plasmid and 5 ng each of the indicated pUASTattB expression constructs encoding Gce and/or Tai. Empty pUASTattB served as the expression-vector control. At 48 h after plasmid transfection, cells were treated with methoprene (MedChemExpress, HY-B1161) at a final concentration of 50 µg/ml, or an equivalent volume of DMSO for 12 h. Firefly and Renilla luciferase activities were measured using the Dual-Luciferase Reporter Assay System (Promega, E1910) on a GloMax 96 Microplate Luminometer (Promega). Firefly luciferase activity was normalized to Renilla luciferase activity, and normalized Fluc/Rluc ratios were expressed relative to the corresponding empty-vector control.

### Live-cell imaging and FRAP in U2OS cells

U2OS cells (Cell Bank of the Chinese Academy of Sciences, SCSP-5030), provided by the laboratory of Xianjue Ma (Westlake University), were cultured, transfected and imaged as described previously ^176^. Briefly, cells were maintained in high-glucose DMEM (Gibco, C11995500CP) supplemented with 10% heat-inactivated fetal bovine serum (ExCell Bio, FSP500) and 1% penicillin–streptomycin (Macklin, Q6532) at 37 °C in a humidified atmosphere containing 5% CO_2_. Cells were seeded into 20-mm glass-bottom culture dishes (NEST, 801001) one day before transfection and transiently transfected with 2 μg per dish of a CMV-driven, pLVX-derived expression plasmid using Lipofectamine LTX (Thermo Fisher, 15338100). The expression cassette encoded *M. pharaonis* Torch residues 25–183 fused in frame to GFP mStayGold (GenBank LC756333.1) and the 3×NLS cassette described above, with a GGGGSGGGGS linker separating each adjacent component. Cells were imaged after overnight expression. Before imaging, nuclei were stained with Hoechst 33342 (Thermo Fisher, H3570; 5 µg/ml) for 10 min in the dark.

Live-cell imaging and FRAP were performed using a Nikon 980 confocal microscope equipped with a 63× oil-immersion objective and a humidified stage-top incubator maintained at 37 °C and 5% CO₂. mStayGold was excited using a 488-nm laser, with an exposure time of 500 ms. A circular region of interest (ROI; 2 µm in diameter) within an individual nuclear Torch–mStayGold–3×NLS condensate was bleached using a 405-nm, 500-mW diode laser at 30% power with a dwell time of 20 ms. A single pre-bleach frame was acquired, and fluorescence recovery was recorded at 1 frame s⁻¹ for 60 s. Fluorescence intensities were extracted in Nikon NIS-Elements AR software from the bleached condensate and an unbleached reference condensate in the same field of view. At each time point, the fluorescence intensity of the bleached condensate was divided by that of the unbleached reference condensate, and the resulting ratio was normalized to its pre-bleach value. For curve fitting, the first post-bleach measurement was defined as *t* = 0. Recovery curves were fitted in GraphPad Prism v11.0.0 using a one-phase association model, *F*(*t*) = *F*_0_ + (*F*_∞_ − *F*_0_)(1 − *e^−kt^*), where *F*_0_ and *F*_∞_ denote the normalized fluorescence immediately after bleaching and at the fitted plateau, respectively. The recovery half-time was calculated as ln(2)/*k*, and fractional recovery as (*F*_∞_ − *F*_0_)/(1− *F*_0_).

### Bimolecular fluorescence complementation (BiFC) in HEK293T cells

BiFC was used to test whether Torch bearing a 3×NLS tag associates with human Ran in mammalian cells. In BiFC, two non-fluorescent fragments of yellow fluorescent protein Venus are fused to candidate proteins; interaction-driven proximity reconstitutes a fluorescent Venus, enabling visualization of protein-protein interactions in living cells ^177^. Torch (residues 25–183; accession number XP_036149432.1) fused to a C-terminal 3×NLS (nucleoplasmin-cMyc-cMyc; described above) was cloned into a pcDNA3.1 vector carrying the Venus C-fragment (VC155; residues 155–238), and human Ran (residues 1–216; accession number NP_006316.1) was cloned into the complementary Venus N-fragment vector (VN173; residues 1–173). Empty-fragment vectors were included.

The HEK293T cell line was obtained from the Kunming Cell Bank, Kunming Institute of Zoology in China. Cells were maintained in Dulbecco’s modified Eagle’s medium (DMEM) supplemented with 10% fetal bovine serum (FBS) at 37 °C in a 5% CO₂ and seeded on coverslips in 24-well plates (60-70% confluence at transfection). Cells were co-transfected with pcDNA3.1-VC155-Torch-3×NLS and pcDNA3.1-VN173-human Ran using Lipofectamine 2000 (DNA: reagent = 1:1; per-well mixture prepared in serum-free DMEM), alongside control combinations (VC155-empty + VN173-empty; VC155-Torch-3×NLS + VN173-empty; VC155-empty + VN173-human Ran). After 4-6 h, media were replaced with complete DMEM and cells were incubated for about 48 h. Cells were gently fixed in 4% paraformaldehyde for 10-15 min, rinsed in PBS, mounted with antifade medium, and were imaged by laser-scanning confocal microscopy (Leica TCS SP8) to detect yellow fluorescence together with transmitted-light images.

### Lentiviral generation of stable HEK293T cells expressing Torch or Torch-3×NLS

Human-codon-optimized sequences encoding His-tagged Torch (residues 25–183; accession number XP_036149432.1), either alone or fused at the C terminus to a 3×NLS cassette described above, were cloned downstream of the UbC promoter in a modified FUGW vector carrying a puromycin-resistance cassette for constitutive expression and selection. For lentiviral production, HEK293T cells grown in 10-cm dishes were transfected with a mixture containing 12.5 μg of total plasmid DNA (FUGW-*torch* or FUGW-*torch-3×NLS* expression construct, packaging plasmid psPAX2, and envelope plasmid pMD2.G at a ratio of 3:2:1) and 25 μL of Lipofectamine 3000 transfection reagent (Thermo Fisher, L3000015) per dish. The transfection was performed according to the manufacturer’s protocol. After 48 hours, the lentivirus-containing cell culture supernatant was collected. For stable expression of His-Torch or His-Torch-3×NLS, HEK293T cells seeded in a 6-well plate at approximately 50% confluence were incubated with a solution consisting of 1 mL of lentiviral supernatant and 1 mL of DMEM supplemented with 10% FBS. Following a 48-hour incubation at 37 °C in 5% CO₂, the medium was replaced with DMEM containing 2 μg/mL puromycin to select for cells stably expressing His-Torch or His-Torch-3×NLS. Selection was continued until no further cell death was observed, indicating successful establishment of stable expression. GFP was cloned into the same vector, and stable empty-vector and GFP-expressing HEK293T cells generated using the same procedure served as the negative control for the indicated assays.

### EdU incorporation assays in HEK293T cells

Stable HEK293T cell lines expressing His-tagged Torch, His-tagged Torch–3×NLS or GFP were pulsed with 10 µM EdU for 2 h and processed using the BeyoClick EdU Cell Proliferation Kit with AF555 (Beyotime, C0075S). Cells were fixed in 4% paraformaldehyde for 15 min, permeabilized with 0.3% PBSTx for 10 min and subjected to click labelling for 30 min at room temperature in the dark. For imaging, parallel cultures grown on poly-D-lysine-coated chamber slides were counterstained with DAPI, mounted and imaged on an Olympus FV4000 confocal microscope; the percentage of EdU-positive nuclei was quantified in Fiji/ImageJ. For flow cytometry, labelled cells were analyzed on a BD Influx cytometer, and data were processed in FlowJo™ v7.6 (BD Biosciences). No-EdU controls were used to define EdU-positive gates, which were applied uniformly across groups, and GFP-expressing cells served as the negative control.

### Immunofluorescence in HEK293T cells

Stable HEK293T cells expressing His-tagged Torch or His-tagged Torch–3×NLS were seeded on four-well poly-D-lysine-coated chamber slides (Corning, 354577) and fixed 24 h later in 4% paraformaldehyde for 10 min. Cells were permeabilized with 0.1% PBSTx for 10 min, blocked in 5% BSA and incubated overnight at 4 °C with His-Tag (D3I1O) rabbit monoclonal antibody (1:500; Cell Signaling Technology, 12698), followed by Alexa Fluor 488-conjugated anti-Rabbit secondary antibody (1:1000; Thermo Fisher Scientific, A11034) incubation for 1 h at 37 °C. Nuclei were counterstained with DAPI, and samples were mounted and imaged on an Olympus FV4000 confocal microscope.

### Mouse strains, husbandry and genotyping

To test Torch function in mammals, a Cre-inducible Rosa26 knock-in line carrying an LSL (loxP-STOP-loxP) cassette containing the full-length *M. pharaonis torch* coding sequence (encoding residues 1–183; XP_036149432.1) was generated on a C57BL/6J background by Shanghai Model Organisms Center, Inc. (hereafter R26-LSL-*torch*). CMV-Cre mice (C57BL/6JCya background; Cyagen Biosciences, stock no. C001055) were purchased from Cyagen Biosciences (Suzhou, China), and Vasa-Cre mice (FVB-Tg(Ddx4-cre)1Dcas/J; The Jackson Laboratory, stock no. 006954) were provided by Bingyu Mao. R26-LSL-*torch* mice were crossed with CMV-Cre or Vasa-Cre mice to generate systemic and germline-directed *torch*-expression cohorts, respectively. Double-positive CMV-Cre;R26-LSL-*torch* and Vasa-Cre;R26-LSL-*torch* mice constituted the experimental groups; where indicated, the corresponding Cre-only and R26-LSL-*torch*-only littermates served as controls. All mice were maintained under specific-pathogen-free conditions at the Kunming Institute of Zoology, Chinese Academy of Sciences, on a 12-h light/12-h dark cycle at 22 °C and 47% relative humidity, with sterilized food and water available ad libitum. Mouse sex was assigned at 1–2 months of age by external examination of the anogenital region, based on the longer anogenital distance in males and confirmed by visible nipples in females or scrotal testes in males. All animal procedures were performed in accordance with protocols approved by the Institutional Animal Care and Use Committee of the Kunming Institute of Zoology, Chinese Academy of Sciences (IACUC protocol no. IACUC-OE-2025-09-005). Cre transgenes were genotyped with forward primer 5’-GCCTGCATTACCGGTCGATGC-3’ and reverse primer 5’-CAGGGTGTTATAAGCAATCCC-3’, yielding a 500-bp amplicon. The R26-LSL-*torch* allele was genotyped with forward primer 5’-TTTGTCGCTGTGGGAAATGC-3’ and reverse primer 5’-AGAAATCTTCATCGGTGCACC-3’, yielding a 400-bp amplicon.

### Mouse western blotting

Liver tissue was collected from two-month-old male CMV-Cre;R26-LSL-*torch* and CMV-Cre-only mice. Total protein was extracted using RIPA lysis buffer supplemented with protease inhibitors, and protein concentrations were determined using the BCA Protein Assay Kit (Beyotime, P0012). Equal amounts of protein (20 μg per lane) were separated by SDS–PAGE and transferred to 0.22 μm PVDF membranes. Torch was detected using the same custom rabbit polyclonal anti-Torch antibody used for the *M. pharaonis* experiments (1:1000), and GAPDH was detected using a mouse monoclonal antibody (clone 1E6D9; Proteintech, 60004-1-Ig; 1:1,000) as a loading control. Membranes were subsequently incubated with HRP-labeled Goat Anti-Rabbit IgG(H+L) (Beyotime, P0948, 1:1000) or Goat Anti-Mouse IgG(H+L) (Beyotime, P0946, 1:1000), and signals were detected using enhanced chemiluminescence reagents (WBKLS0500, Millipore, USA) and Bio-Rad Gel Doc XR+ system.

### Mouse body and organ weight measurements

Body weights of male and female mice were recorded weekly from 4 to 8 weeks of age. At 2 months of age, mice were euthanized by an intraperitoneal overdose of sodium pentobarbital before tissue collection, and the brain, liver, kidneys, spleen and heart were dissected and weighed; testes were additionally collected from males. Organ-to-body-weight ratios were calculated where indicated.

### Mouse testis SOX9 immunofluorescence and Sertoli-cell counting

Two-month-old male mice were deeply anesthetized with sodium pentobarbital and transcardially perfused with PBS. Testes were dissected, fixed in 4% paraformaldehyde for 48 h and dehydrated sequentially in 75% ethanol for 4 h, 85% ethanol for 2 h, 90% ethanol for 2 h, 95% ethanol for 1 h and 100% ethanol for 1 h. Samples were cleared in a 1:1 ethanol:xylene mixture for 5–10 min and then twice in 100% xylene for 10 min each. Tissues were infiltrated with paraffin at 65 °C for 1 h, transferred through three fresh paraffin changes for 1 h each and embedded. Paraffin blocks were sectioned at 5 μm. Sections were dried at 65 °C for 3 h, deparaffinized three times for 30 min each in TO clearing agent (Guangzhou Zhongnan Chemical Instrument Co., Ltd.) and rehydrated through 100%, 95%, 75% and 35% ethanol (three changes per concentration, 2–5 min each), followed by two PBS washes. Heat-induced epitope retrieval was performed by boiling the sections for 15 min in 1× Tris–EDTA antigen-retrieval solution (pH 9.0; Beyotime, P0084), followed by cooling at room temperature for 15 min. Sections were washed in PBS, permeabilized in PBS containing 0.5% Triton X-100 for 30 min and blocked with 5% BSA for 1 h. Sertoli cells were labeled overnight at 4 °C with rabbit anti-SOX9 polyclonal antibody (1:200; Oasis Biofarm, OB-PRB049). After three PBS washes, sections were incubated for 1 h at room temperature in the dark with Alexa Fluor 488-conjugated anti-Rabbit secondary antibody (1:1000; Thermo Fisher Scientific, A11034), and counterstained with DAPI. Images were acquired using a TissueFAXS Q-FLUO-S imaging system (TissueGnostics) or an Olympus IXplore SpinSR10 microscope and processed with TissueFAXS software or cellSens Dimension v4.4, respectively. Identical acquisition settings were used for samples included in the same quantitative comparison. For each mouse, five randomly selected fields were imaged from SOX9-stained testis sections. SOX9-positive Sertoli-cell nuclei were manually counted in 25 randomly selected seminiferous-tubule cross-sections across these fields, and the mean number per tubule was calculated for each mouse and plotted in Fig. 6k.

### RNA extraction across experimental systems and microsatellite genotyping in ants

Immediately after collection or dissection, samples were snap-frozen in liquid nitrogen and subsequently homogenized for RNA extraction. For *M. pharaonis* brood, total RNA was extracted individually from whole third-instar larvae, prepupae, and pupae using the RNeasy Plus Micro Kit (QIAGEN) according to the manufacturer’s instructions. Samples were homogenized with 5-mm stainless-steel beads in Buffer RLT Plus supplemented with Reagent DX and β-mercaptoethanol. Genomic DNA retained on the gDNA Eliminator columns from third-instar larvae and prepupae was subsequently recovered by washing and eluting. To distinguish haploid males from diploid gynes among reproductive-destined larvae and prepupae, individuals were genotyped at five highly polymorphic nuclear microsatellite loci—Mp4, Mp8, Mph2, Mph23, and Mph9^178^. PCR fragment sizes were resolved on an Applied Biosystems 3130xl Genetic Analyzer, and alleles were called using GeneMapper v4.0 (Applied Biosystems). Total RNA from dissected adult tissues of *M. pharaonis* and *D. melanogaster*, *Carebara diversa* samples, cultured HEK-293T cells, and dissected mouse tissues was extracted using TRIzol Reagent (Thermo Fisher Scientific; 15596018) according to the manufacturer’s instructions. Following chloroform extraction and phase separation, RNA was precipitated from the aqueous phase with isopropanol, washed with 75% ethanol, briefly air-dried, and resuspended in nuclease-free water. Purified RNA was stored at −80 °C until further analysis.

### RNA sequencing and quality control

RNA quality control, complementary DNA (cDNA) library construction, and sequencing were performed at BGI (Shenzhen, China) and Novogene (Beijing, China). RNA integrity was assessed on an Agilent 2100 Bioanalyzer. Ambion ERCC RNA Spike-In Mix (Thermo Fisher, 4456740) was added prior to cDNA library construction. cDNA synthesis followed the Smart-seq2 workflow ^179^, followed by random fragmentation, PCR amplification, and size selection (150-350 bp) prior to circularization. Libraries were sequenced on the BGI DNBSEQ platform using a 100-nt paired-end protocol. Raw paired-end reads were subjected to quality and adapter trimming using Trim Galore (v0.6.10; RRID: SCR_011847), which uses Cutadapt for adapter removal ^180^. Filtered reads were aligned to the corresponding reference genome assemblies using HISAT2 (v2.2.1) ^181^. The resulting alignment files were converted from SAM to BAM format using SAMtools (v1.6) ^182^. Reads mapping to annotated genomic features were subsequently quantified using featureCounts (v2.0.6) ^183^.

### Differential expression, orthology mapping and functional enrichment analyses

Count normalization and differential expression analysis were performed using DESeq2 (v1.36.0) in R ^184^. Genes were designated as differentially expressed (DEGs) if they met a false discovery rate-adjusted *P* value < 0.05. For heterologous RNA-seq analyses, the exonic sequence of *M. pharaonis torch*, together with its corresponding gene annotation, was appended to each host reference genome and annotation before read alignment and gene-level quantification. Functional annotations of *M. pharaonis* genes were assigned from their *D. melanogaster* orthologues using the previously established OrthoFinder v2.5.4-based orthology map across 17 insect proteomes; when multiple *Drosophila* orthologues were identified, the most similar orthologue was retained ^42^. Orthologous relationships among *D. melanogaster*, *Homo sapiens* and *Mus musculus* genes were identified using the DRSC Integrative Ortholog Prediction Tool (DIOPT) Ortholog Finder. Functional enrichment was assessed by applying Generally Applicable Gene-set Enrichment (GAGE) ^185^ to gene-level *t*-statistics from the differential expression analyses (R package gage, v2.58), with same.dir = TRUE to identify pathways showing coherent up– or downregulation. Gene Ontology (GO) Biological Process terms ^186^ and Kyoto Encyclopedia of Genes and Genomes (KEGG) pathways ^187,188^ with nominal *P* value < 0.01 were retained for cross-system comparisons; FDR-adjusted *q* values are reported where indicated. For systems with both transcriptomic and CUT&Tag data, direct Torch targets were defined as the intersection of DEGs and genes carrying promoter-proximal Torch peaks (−3 kb to +1 kb relative to the transcription start site). Orthologous direct-target sets were subsequently intersected across species to identify conserved Torch-regulated genes.

### CUT&Tag profiling and data analysis

CUT&Tag was performed essentially as described previously ^189^. Nuclei were isolated from whole-body *Monomorium pharaonis* gyne and worker broods comprising mixed third-instar larvae, prepupae and early pupae (six biological replicates per caste); heads of female *Drosophila melanogaster* expressing *kibra* with or without *torch–3×NLS* under *GMR-GAL4* control (two biological replicates per group); and brains and kidneys from male CMV-Cre;R26-LSL-*torch* mice (three biological replicates per tissue) and CMV-Cre-only controls (two biological replicates per tissue). Stable HEK293T cells carrying a genome-integrated construct encoding His-tagged Torch–3×NLS (four biological replicates) and matched control cells carrying a genome-integrated empty-vector construct (two biological replicates) were analyzed in parallel. Nuclei from all tissue samples were isolated by homogenization in 1× HB buffer followed by filtration and density-gradient separation. The affinity-purified custom rabbit anti-Torch antibody described above was used for ant, fly and mouse samples, whereas HEK293T samples were incubated with a His-Tag (D3I1O) rabbit antibody (Cell Signaling Technology, 12698). A total of 2 μg primary antibody was used per CUT&Tag library. Cells or nuclei were immobilized on concanavalin A-coated beads, permeabilized with digitonin and sequentially incubated with primary antibody, secondary antibody and protein A–Tn5 transposome. Following Mg²⁺-activated tagmentation, libraries were PCR-amplified, purified using AMPure XP beads, assessed using an Agilent 2100 Bioanalyzer and qPCR, and sequenced on an Illumina NovaSeq platform using 150-bp paired-end reads.

Read quality was assessed using FastQC v.0.11.3. Raw reads were filtered using fastp v.0.20.0 (––length_required 15 –-n_base_limit 6) ^190^ and aligned using BWA v.0.7.12 (– k 32 –T 30 –t 4 –M) ^191^ to the corresponding reference genome: *M. pharaonis* ASM1337386v2 reference genome (RefSeq GCF_013373865.1), the *D. melanogaster* Release 6 reference genome (GenBank GCA_000001215.4; FlyBase annotation release 6.44, FB2022_01), the *H. sapiens* GRCh38.p14 primary assembly (GenBank GCA_000001405.29; Ensembl release 110) or the *M. musculus* GRCm39 primary assembly (GenBank GCA_000001635.9; Ensembl release 108), as appropriate. Uniquely mapped, properly paired reads with MAPQ ≥ 13 were retained, and PCR duplicates were removed. Peaks were called independently for each replicate using MACS2 v.2.1.0 (–q 0.05 –-call-summits –f BAMPE –-keep-dup all) ^192^. Genome-wide signal correlations between samples were evaluated by Pearson correlation using deepTools v.3.0.2 (––corMethod pearson) ^193^, which was also used to generate coverage tracks. For each ant caste, peaks from the six replicates were merged using BEDTools^194^ to generate separate gyne– and worker-brood union sets. For each heterologous system, peaks from the Torch-expressing replicates were merged, and any merged peak overlapping a peak detected in either Torch-negative control replicate was discarded. Peaks were annotated as promoter-associated (−3 kb to +1 kb relative to the transcription start site), exonic, intronic or intergenic using BEDTools. De novo motif discovery was performed on 500-bp sequences centered on peak summits using the MEME-ChIP web server in MEME Suite v.5.5.9 ^195^. CentriMo ^196^ was used to assess positional enrichment, Tomtom ^197^ to compare discovered motifs with reference motifs, and FIMO ^198^ to identify GA-repeat motif occurrences within selected promoter-associated peaks.

### Statistics and reproducibility

Statistical analyses were performed using GraphPad Prism v11.0.0 or R v4.4.2. Data presentation, sample sizes, statistical tests, degrees of freedom, and exact or multiplicity-adjusted *P* values are provided in the corresponding figures and legends. No formal statistical calculation was used to predetermine sample sizes; sample sizes were guided by previous studies and sample availability. No randomization or blinding procedures were used. No samples or data points were excluded from the analyses. All cultured cell lines were routinely tested for mycoplasma contamination and remained negative throughout the study.

## Data availability

The raw RNA-sequencing and CUT&Tag data generated in this study, together with the associated sample metadata, have been deposited in the NCBI repository (BioProject Accession Number: PRJNA1495256). The metadata specify the number of biological replicates per group, the biological material represented by each library (tissues or cultured cells, as applicable), experimental group assignments for the ant, *Drosophila*, HEK293T and mouse RNA-seq datasets, and the age and sex of the mice. The mass spectrometry proteomics data have been deposited to the ProteomeXchange Consortium via the PRIDE ^199^ partner repository with the dataset identifier PXD081382. These datasets will be released upon publication. Previously published transcriptomic datasets reanalyzed in this study comprised the multispecies adult gyne–worker transcriptomes generated by the *Global Ant Genomics Alliance* (*GAGA*) (NCBI BioProject PRJNA1172379) ^23^, the developmental JH-treatment transcriptomes of *M. pharaonis* (PRJNA1143187) ^2^, and the caste-specific developmental transcriptomes of *M. pharaonis* and *Acromyrmex echinatior* (PRJNA767561) ^42^.

## Code availability

No custom software or algorithms central to the conclusions of this study were developed. All computational analyses were performed using publicly available software and standard packages, as described in the Methods.

## Acknowledgements

We thank the Information Technology Center of Zhejiang University and China Mobile Zhejiang Co., Ltd. (Hangzhou Branch) for providing computational resources; the Center for Advanced Bioimaging Denmark at the University of Copenhagen for access to confocal microscopy and associated software; the Institutional Center for Shared Technologies and Facilities of the Kunming Institute of Zoology, Chinese Academy of Sciences, for support with confocal image acquisition and flow-cytometric analysis; and the staff of the National Research Facility for Phenotypic and Genetic Analysis of Model Animals (Primate Facility; https://cstr.cn/31137.02.NPRC) for technical support with data collection and analysis. We thank Siwei Liang for advice on data visualization; Alivia Lee Price, Guo Ding, Jinda Li and Cong Li for technical assistance; Zhuang Wu, Xiao Zhang, Ling Zhao and Li Ma for animal husbandry; and Bingyu Mao for providing Vasa-Cre mice.

## Author notes

These authors contributed equally: Ruyan Li, Sha Song, Xiafang Zhang.

## Author contributions

X.M., W.L. and G.Z. conceived and supervised the project. R.L., S.S., X.Z., Y.Q., Y.L. and D.Z. designed and performed experiments in ants, *Drosophila*, cultured cells and mice, and generated and analyzed data from RNA-seq, CUT&Tag, imaging, flow cytometry and mass spectrometry. R.L., F.L., J.V., Z.X., Jixuan Zheng, B.Q., Y.S. and Z.H. performed the comparative genomic, phylogenetic and integrative bioinformatic analyses. R.S.L., Y.L., Jie Zhao, W.D., J.W., H.R., Q.L., X.M., W.L. and G.Z. provided materials, resources and technical support. R.L., X.Z., J.J.B., X.M., W.L. and G.Z. wrote and revised the manuscript with input from all authors.

## Additional information

Supplementary information is available for this paper. Correspondence and requests for materials should be addressed to Xianjue Ma, Weiwei Liu or Guojie Zhang.

