## Supplementary Information for "A Hymenoptera-restricted gene mediating ant castes co-opts deeply conserved machinery to control organ size"

This file contains Supplementary Information Text describing the design and validation of splice-blocking vivo-morpholinos, two Supplementary Figures, ten Extended Data Figures, descriptions of Supplementary Datasets 1–15, and supplementary references supporting the study of Torch-mediated organ growth.

**Supplementary Information Text**

**Design and validation of splice-blocking vivo-morpholinos (vivo-MOs) targeting the *torch* pre-mRNA**

A splice-blocking vivo-MO binds across an exon-intron junction downstream of the start codon on an unspliced pre-mRNA and sterically blocks the spliceosome at that site [1]. If the junction is the first or last in the pre-mRNA, splice blocking forces intron inclusion by inserting an intron-derived segment into the protein; otherwise, splice blocking causes exon skipping when the target junction lies within the pre-mRNA, so the mature protein lacks that exon-encoded segment [2]. Either scenario typically compromises protein function. Moreover, when the skipped exon or retained intron is non-cassette (i.e. its length not being divisible by three), the downstream reading frame is shifted, introducing a premature termination codon (PTC) that triggers nonsense-mediated mRNA decay (NMD) and eliminates the transcript [3].

Guided by these principles, we designed two vivo-MOs to target the exon 2-intron 2 (E2I2) and intron 3-exon 4 (I3E4) boundaries of the *torch* pre-mRNA - junctions shared by both NCBI-annotated isoforms. E2I2-vivo-MO was expected to trigger skipping of exon 2 - which harbors the start codon, whereas I3E4-vivo-MO should enforce intron 3 inclusion; and because neither fragment's length is a multiple of three both manipulations should frameshift the coding sequence and introduce a PTC, triggering NMD (Extended Data Fig. 2a). This splice blocking followed by NMD was thus expected to eradicate nearly all *torch* mRNA.

Droplet digital PCR (ddPCR) confirmed that each vivo-MO reduced whole-body *torch* transcript levels by ~90% within 24 h of injection in late third-instar larvae, with depletion sustained through the prepupal stage at 96 h post-injection (Extended Data Fig. 2b-c; all $P < 0.0001$). Consistent with these ddPCR results, RNA-seq confirmed >90% depletion of *torch* transcripts from late third-instar larvae to early pupae, spanning 24 h to 7 d post-injection (Extended Data Fig. 8a-c; Supplementary Dataset 11; all adjusted $P < 0.0001$). Together, these data show that our vivo-MO targeting of *torch* pre-mRNA achieved robust and durable knockdown from the late larval stage through to pupal organ growth, the developmental window during which caste-differentiated organs undergo active expansion.

**Design and validation of splice-blocking vivo-MOs targeting the *gce* pre-mRNA and *tai* pre-mRNA of the JH receptor complex**

To test whether compromising the JH receptor Gce and its co-activator Tai reduced *torch* transcription, we designed two splice-blocking vivo-MOs targeting splice junctions for each of these genes. For the *gce* transcript (NCBI accession number: XM_036283176.1) the oligonucleotides span the intron 3-exon 4 (I3E4) and exon 6-intron 6 (E6I6) junctions (Extended Data Fig. 3b) while for the *tai* transcript (NCBI accession number: XM_036293388.1) they span intron 2-exon 3 (I2E3) and intron 5-exon 6 (I5E6) (Extended Data Fig. 3c). All four junctions are shared by every annotated isoform, and each targeted exon is non-cassette. Skipping these exons should therefore frameshift the coding sequence and introduce a PTC in downstream sequence followed by NMD as in the *torch* pre-mRNA.

However, contrary to that expectation, ddPCR showed no significant decline in *gce* or *tai* mRNA 24 h after injection with any of the vivo-MOs injected into late third-instar gynes when transcript abundance was quantified with primer pairs whose amplicons do not overlap with the targeted exons (Extended Data Fig. 3d). This observation concurs with growing evidence that a substantial subset of PTC-containing transcripts evade NMD through an array of mechanistically diverse and incompletely elucidated pathways [4,5]. However, PCR with a primer pair whose forward primer lies in the exon immediately upstream of the targeted exon and whose reverse primer lies in the exon immediately downstream - so that the amplicon spans the entire targeted exon (Extended Data Fig. 3e) - followed by agarose-gel electrophoresis produced an additional, lower-molecular-weight band in every treatment both 24 h (late third-instar larvae) and 96 h (prepupae) post-injection (Extended Data Fig. 3f). The appearance of this shorter product is consistent with efficient morpholino-induced exon skipping. Sanger sequencing confirmed complete or partial exon skipping in each case: *gce*-I3E4 removed the entire 83 bp exon 4, thereby frameshifting the downstream coding sequence and introducing a PTC; *gce*-E6I6 excised the terminal 162 bp of exon 6, deleting a 54-amino-acid segment from the mature protein despite preserving the reading frame; *tai*-I2E3 skipped the full 139 bp exon 3 causing a frameshift and introducing a PTC; and *tai*-I5E6 excised the last 2 bp from exon 5 and the first 155 bp of exon 6 (157 bp in total), likewise inducing a frameshift and introducing a PTC (Extended Data Fig. 3g-j; Supplementary Dataset 5). This kind of partial exon skipping likely reflects activation of cryptic splice sites and is a known response to morpholino treatment [6,7].

Thus, unlike the *torch* vivo-MOs that depleted *torch* mRNA almost entirely, the splice-blocking vivo-MOs primarily redirected *gce*/*tai* splicing and generated frameshifted/truncated products predicted to compromise Gce/Tai function, without markedly reducing total *gce*/*tai* transcript abundance, consistent with splice-switching morpholino effects reported in other model systems [6,8].

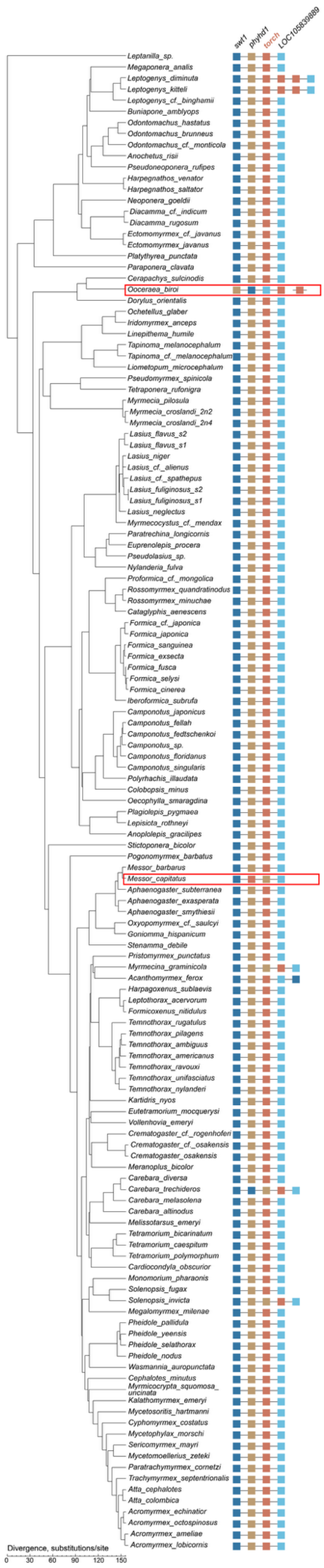
swf1
phyhd1
torch
LOC105839889
Leptanilla_sp.
Megaponera_analis
Leptogenys_diminuta
Leptogenys_kitteli
Leptogenys_cf._binghamii
Buniapone_amblyops
Odontomachus_hastatus
Odontomachus_brunneus
Odontomachus_cf._monticola
Anochetus_risii
Pseudoneoponera_rufipes
Harpegnathos_venator
Harpegnathos_saltator
Neoponera_goeldii
Diacamma_cf._indicum
Diacamma_rugosum
Ectomomyrmex_cf._javanus
Ectomomyrmex_javanus
Platythyrea_punctata
Paraponera_clavata
Cerapachys_sulcinodis
Ooceraea_biroi
Dorylus_orientalis
Ochetellus_glaber
Iridomyrmex_anceps
Linepithema_humile
Tapinoma_melanocephalum
Tapinoma_cf._melanocephalum
Liometopum_microcephalum
Pseudomyrmex_spinicola
Tetraponera_rufonigra
Myrmecia_pilosula
Myrmecia_croslandi_2n2
Myrmecia_croslandi_2n4
Lasius_flavus_s2
Lasius_flavus_s1
Lasius_niger
Lasius_cf._alienus
Lasius_cf._spathepus
Lasius_fuliginosus_s2
Lasius_fuliginosus_s1
Lasius_neglectus
Myrmecocystus_cf._mendax
Paratrechina_longicornis
Euprenolepis_procera
Pseudolasius_sp.
Nylanderia_fulva
Proformica_cf._mongolica
Rossomyrmex_quandratinodus
Rossomyrmex_minuchae
Cataglyphis_aenescens
Formica_cf._japonica
Formica_japonica
Formica_sanguinea
Formica_exsecta
Formica_fusca
Formica_selysi
Formica_cinerea
Iberoformica_subrufa
Camponotus_japonicus
Camponotus_fellah
Camponotus_fedtschenkoi
Camponotus_sp.
Camponotus_floridanus
Camponotus_singularis
Polyrhachis_illaudata
Colobopsis_minus
Oecophylla_smaragdina
Plagiolepis_pygmaea
Lepisiota_rothneyi
Anoplolepis_gracilipes
Stictoponera_bicolor
Pogonomyrmex_barbatus
Messor_barbarus
Messor_capitatus
Aphaenogaster_subterranea
Aphaenogaster_exasperata
Aphaenogaster_smythiesii
Oxyopomyrmex_cf._saulcyi
Goniomma_hispanicum
Stenamma_debile
Pristomyrmex_punctatus
Myrmecina_graminicola
Acanthomyrmex_ferox
Harpagoxenus_sublaevis
Leptothorax_acervorum
Formicoxenus_nitidulus
Temnothorax_rugatulus
Temnothorax_pilagens
Temnothorax_ambiguus
Temnothorax_americanus
Temnothorax_ravouxi
Temnothorax_unifasciatus
Temnothorax_nylanderi
Kartidris_nyos
Eutetramorium_mocquerysi
Vollenhovia_emeryi
Crematogaster_cf._rogenhoferi
Crematogaster_cf._osakensis
Crematogaster_osakensis
Meranoplus_bicolor
Carebara_diversa
Carebara_trechideros
Carebara_melasolena
Carebara_altinodus
Melissotarsus_emeryi
Tetramorium_bicarinatum
Tetramorium_caespitum
Tetramorium_polymorphum
Cardiocondyla_obscurior
Monomorium_pharaonis
Solenopsis_fugax
Solenopsis_invicta
Megalomyrmex_milenae
Pheidole_pallidula
Pheidole_yeensis
Pheidole_selathorax
Pheidole_nodus
Wasmannia_auropunctata
Cephalotes_minutus
Myrmicocrypta_squomosa_uncinata
Kalathomyrmex_emeryi
Mycetosoritis_hartmanni
Cyphomyrmex_costatus
Mycetophylax_morschi
Sericomyrmex_mayri
Mycetomoellerius_zeteki
Paratrachymyrmex_cornetzi
Trachymyrmex_septentrionalis
Atta_cephalotes
Atta_colombica
Acromyrmex_echinatior
Acromyrmex_octospinosus
Acromyrmex_ameliae
Acromyrmex_lobicornis
Divergence, substitutions/site
0
30
60
90
120
150

**Supplementary Fig. 1. Conserved microsynteny around *torch* across the GAGA ant genomes [9].** Synteny maps aligned to the gene phylogeny show a stable genomic neighborhood of *torch* across the 135 high-quality ant genomes with scaffold N50 >1 Mb, selected from the 163 ant genomes generated by the *Global Ant Genomics Alliance* (GAGA) [9]. Across these high-quality assemblies, *torch* resides in a highly conserved microsyntenic block, flanked by *swt1* and *phyhd1* on one side and with *LOC105839889* adjacent to *torch* at the other side, a gene order and orientation preserved in almost all extant ant species. The two exceptions, highlighted in red boxes, are the clonal raider ant *Ooceraea biroi* and *Messor capitatus*, both of which underwent a local inversion involving the *torch* locus. These species are among the few ants whose workers can reproduce by thelytokous parthenogenesis, producing diploid female offspring from unfertilized eggs [10,11], suggesting that aberrant *torch* regulation may be linked to this rare reproductive mode. Two other exceptions *Leptogenys diminuta* and *Leptogenys kitteli* harbor two additional *torch* copies within the same syntenic block, whereas *Ooceraea biroi* carries an additional copy embedded in a separate block and four other species (*Myrmecina graminicola*, *Acanthomyrmex ferox*, *Carebara trechideros*, *Solenopsis invicta*) have duplications of flanking genes. These lineage-specific rearrangements or duplications do not alter the general inference that the ancestral *torch* locus was embedded in a conserved microsyntenic neighborhood. Among the remaining 28 GAGA species excluded from this synteny panel because their assemblies did not meet our very high quality threshold, no detectable *torch* ortholog was identified in seven species: the three sampled Amblyoponinae species (*Stigmatomma sp.*, *Mystrium camillae*, and *Prionopelta kraepelini*), *Probolomyrmex sp.* in the Proceratiinae, *Dinoponera quadriceps* in the Ponerinae, and two Myrmicinae species (*Pheidole capellinii* and *Temnothorax longispinosus*) (Supplementary Dataset 1).

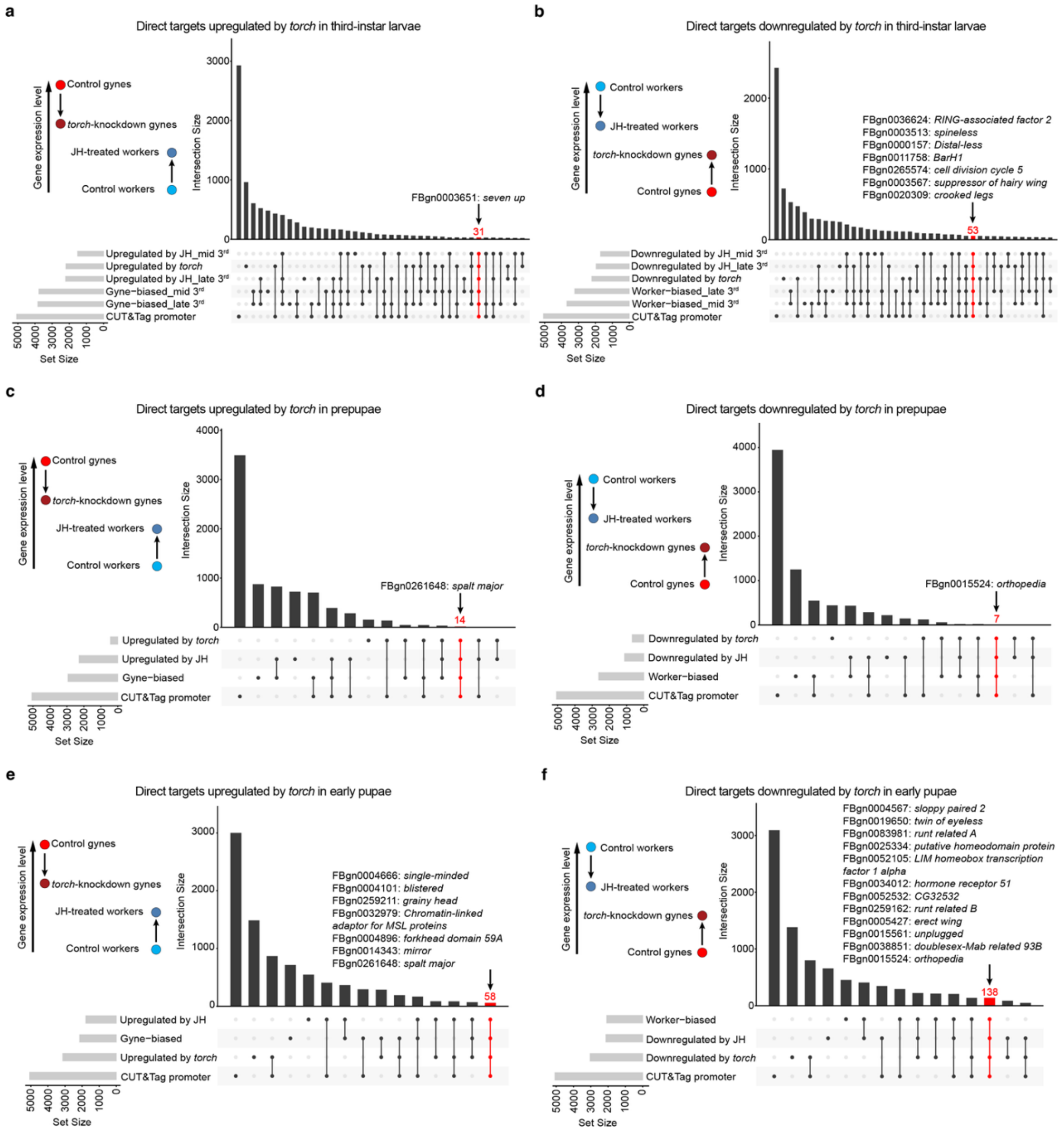
a
Direct targets upregulated by torch in third-instar larvae
Gene expression level
Control gynes
torch-knockdown gynes
JH-treated workers
Control workers
Intersection Size
FBgn0003651: seven up
31
Upregulated by JH_mid 3rd
Upregulated by torch
Upregulated by JH_late 3rd
Gyne−biased_mid 3rd
Gyne−biased_late 3rd
CUT&Tag promoter
Set Size
b
Direct targets downregulated by torch in third-instar larvae
Control workers
JH-treated workers
torch-knockdown gynes
Control gynes
FBgn0036624: RING-associated factor 2
FBgn0003513: spineless
FBgn0000157: Distal-less
FBgn0011758: BarH1
FBgn0265574: cell division cycle 5
FBgn0003567: suppressor of hairy wing
FBgn0020309: crooked legs
53
Downregulated by JH_mid 3rd
Downregulated by JH_late 3rd
Downregulated by torch
Worker−biased_late 3rd
Worker−biased_mid 3rd
CUT&Tag promoter
c
Direct targets upregulated by torch in prepupae
FBgn0261648: spalt major
14
Upregulated by torch
Upregulated by JH
Gyne−biased
CUT&Tag promoter
d
Direct targets downregulated by torch in prepupae
FBgn0015524: orthopedia
7
Downregulated by torch
Downregulated by JH
Worker−biased
CUT&Tag promoter
e
Direct targets upregulated by torch in early pupae
FBgn0004666: single-minded
FBgn0004101: blistered
FBgn0259211: grainy head
FBgn0032979: Chromatin-linked adaptor for MSL proteins
FBgn0004896: forkhead domain 59A
FBgn0014343: mirror
FBgn0261648: spalt major
58
Upregulated by JH
Gyne−biased
Upregulated by torch
CUT&Tag promoter
f
Direct targets downregulated by torch in early pupae
FBgn0004567: sloppy paired 2
FBgn0019650: twin of eyeless
FBgn0083981: runt related A
FBgn0025334: putative homeodomain protein factor 1 alpha
FBgn0052105: LIM homeobox transcription
FBgn0034012: hormone receptor 51
FBgn0052532: CG32532
FBgn0259162: runt related B
FBgn0005427: erect wing
FBgn0015561: unplugged
FBgn0038851: doublesex-Mab related 93B
FBgn0015524: orthopedia
138
Worker−biased
Downregulated by JH
Downregulated by torch
CUT&Tag promoter

**Supplementary Fig. 2. Stage-resolved core targets of *torch* defined by UpSet set intersections.** UpSet plots integrating gene sets of third-instar larvae (24 h post-injection; **a-b**), prepupae (96 h post-injection; **c-d**) and early pupae (7 d post-injection; **e-f**) to define direct targets of core *torch*-activation (a, c, e) and *torch*-repression (b, d, f). Sets comprise (i) genes positively or negatively regulated by *torch* (DEGs decreased or increased in *torch*-knockdown gynes versus control gynes), (ii) gyne- or worker-biased DEGs (gynes versus workers), (iii) JH-upregulated or JH-downregulated DEGs (JH-treated versus control workers), and (iv) genes with Torch CUT&Tag promoter peaks in gyne broods. We defined core *torch*-activated targets as genes whose promoters carried a Torch CUT&Tag peak and whose expression decreased upon *torch* knockdown and was also gyne-biased and JH-upregulated; conversely, core *torch*-repressed targets were defined as genes whose promoters carried a Torch CUT&Tag peak and whose expression increased upon *torch* knockdown and was also worker-biased and JH-downregulated. Dots denote set membership and bars indicate intersection size. Because the published third-instar larval data include both medium- and late-third-instar larvae [12], we retained only caste-biased and JH-responsive DEGs shared by both stages when defining core targets for third-instar larvae (a-b), thereby focusing on genes consistently regulated across the larval stages. Core targets defined by the four-way intersection are highlighted in red, with this intersection size labelled; and annotated transcription factors (based on *D. melanogaster* homologs) are listed as follows: third-instar larvae (upregulated by *torch*): *seven up*; third-instar larvae (downregulated by *torch*): *RING-associated factor 2*, *spineless*, *Distal-less*, *BarH1*, *cell division cycle 5*, *suppressor of hairy wing*, *crooked legs*; prepupae (upregulated by *torch*): *spalt major*; prepupae (downregulated by *torch*): *orthopedia*; early pupae (upregulated by *torch*): *single-minded*, *blistered*, *grainy head*, *Chromatin-linked adaptor for MSL proteins*, *forkhead domain 59A*, *mirror*, *spalt major*; early pupae (downregulated by *torch*): *sloppy paired 2*, *twin of eyeless*, *runt related A*, *putative homeodomain protein*, *LIM homeobox transcription factor 1 alpha*, *hormone receptor 51*, *CG32532*, *runt-related B*, *erect wing*, *unplugged*, *doublesex-Mab related 93B*, *orthopedia*. Differentially expressed genes (DEGs) were defined by an adjusted $P$ value $< 0.05$. Complete source lists are provided in Supplementary Dataset 3 for stage-resolved caste-biased DEGs (gynes versus control workers) and JH-responsive DEGs (JH-treated versus control workers) [12], Supplementary Dataset 9 for genes with Torch-bound promoters identified by CUT&Tag in gyne broods, and Supplementary Dataset 11 for stage-resolved DEGs following *torch* depletion in gynes.

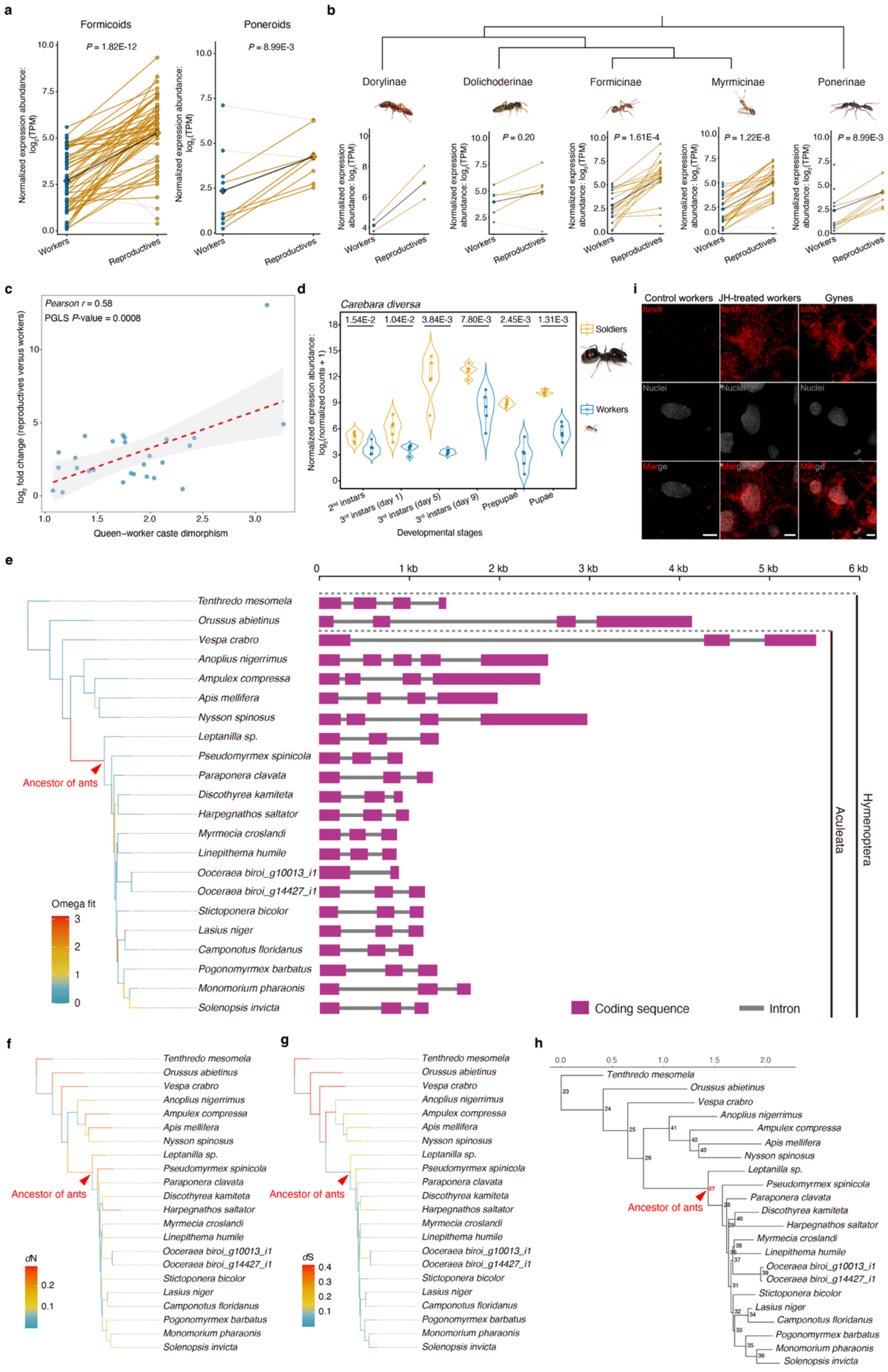

a
Formicoids
P = 1.82E-12
Poneroids
P = 8.99E-3
Normalized expression abundance: log2(TPM)
Workers
Reproductives
b
Dorylinae
Dolichoderinae
Formicinae
Myrmicinae
Ponerinae
P = 0.20
P = 1.61E-4
P = 1.22E-8
P = 8.99E-3
c
Pearson r = 0.58
PGLS P-value = 0.0008
log2 fold change (reproductives versus workers)
Queen–worker caste dimorphism
d
Carebara diversa
1.54E-2
1.04E-2
3.84E-3
7.80E-3
2.45E-3
1.31E-3
Soldiers
Workers
Normalized expression abundance: log2(normalized counts + 1)
2nd instars
3rd instars (day 1)
3rd instars (day 5)
3rd instars (day 9)
Prepupae
Pupae
Developmental stages
i
Control workers
JH-treated workers
Gynes
torch
Nuclei
Merge
e
Tenthredo mesomela
Orussus abietinus
Vespa crabro
Anoplius nigerrimus
Ampulex compressa
Apis mellifera
Nysson spinosus
Leptanilla sp.
Pseudomyrmex spinicola
Paraponera clavata
Discothyrea kamiteta
Harpegnathos saltator
Myrmecia croslandi
Linepithema humile
Ooceraea biroi_g10013_i1
Ooceraea biroi_g14427_i1
Stictoponera bicolor
Lasius niger
Camponotus floridanus
Pogonomyrmex barbatus
Monomorium pharaonis
Solenopsis invicta
Ancestor of ants
Omega fit
Coding sequence
Intron
Aculeata
Hymenoptera
f
dN
g
dS
h

**Extended Data Fig. 1. The *torch* gene shows phylogenetically conserved caste-biased expression, ant-lineage-specific coding-sequence remodeling, and juvenile hormone (JH) responsiveness. (a–b)** Reproductive (gyne/queen)-biased *torch* expression across the two main formicoid and poneroid clades of the ants (a) and across five major ant subfamilies (b), illustrated by the positive slopes of the black lines connecting paired reproductive and worker expression values. See Fig. 1b for additional legend details. Statistical significances were assessed using two-sided paired *t*-tests ($n = 56$, $t = 9.04$, df = 55 for Formicoids; $n = 9$, $t = 3.43$, df = 8 for Poneroids (Ponerinae); $n = 5$, $t = 1.52$, df = 4 for Dolichoderinae; $n = 19$, $t = 4.75$, df = 18 for Formicinae; $n = 29$, $t = 7.93$, df = 28 for Myrmicinae). No statistical test was performed for Dorylinae because of the small sample size. Besides the five subfamilies shown in panel b, Pseudomyrmecinae is represented by one species, *Tetraponera rufonigra*, which also shows reproductive-biased *torch* expression (Supplementary Dataset 1). In Dolichoderinae, the difference did not reach statistical significance ($P = 0.20$), but the across-species mean expression was higher in reproductives than in workers, indicating the same overall trend. In contrast, *torch* expression was higher in adult workers than reproductives for eight species, as illustrated by grey connecting lines: *Megaponera analis* and *Ectomomyrmex javanus* from the Ponerinae; *Liometopum microcephalum* from the Dolichoderinae; *Oecophylla smaragdina* from the Formicinae; and *Carebara lignata*, *Vollenhovia emeryi*, *Mycetophylax morschi*, and *Stenamma debile* from the Myrmicinae (Supplementary Dataset 1). Worker-biased *torch* expression in the clonally reproducing ant *V. emeryi* [13] was further supported by an independent study reporting a $\log_2$ fold change of −1.31 for queens versus workers, with an adjusted *P* value of $1.41 \times 10^{-6}$ [14]. Because these eight species with exceptional expression bias are phylogenetically scattered across the Formicidae rather than concentrated in the basal ant branches, they likely reflect independent, lineage-specific secondary modifications in *torch* regulation. Together with the predominant gyne-biased expression observed across most ant species, this distribution supports gyne-biased *torch* expression as the likely ancestral regulatory state in ants. **(c)** Phylogenetic generalized least squares (PGLS) regression across 27 ant species showing that *torch*'s reproductive female (gyne/queen)-biased expression (the $\log_2$ fold change in mean TPM between reproductive female and workers) in adult transcriptomes increases with queen–worker caste dimorphism using scores obtained from a published study [9]. Statistical significance was assessed using a two-sided *t*-test of the dimorphism coefficient ($t = 3.836$, df = 25, $P = 0.0008$). Points represent observed species values. The dashed line and shaded area show the ordinary least-squares trend line and 95% confidence interval for visualization. **(d)** In *Carebara diversa*, soldiers show consistently elevated expression of *torch* relative to workers across the larval and pupal stages based on developmental transcriptomes, differentiated by day after the start of the third instar under our rearing conditions. Expression values are shown as log2-transformed normalized counts. For each developmental stage, worker and soldier samples were compared using two-sided Welch's unpaired *t*-tests. *P* values were adjusted across the six developmental-stage comparisons using the Benjamini–Hochberg false discovery rate correction and are indicated in the figure. $n = 5$ biological replicates per caste per stage. Statistics were calculated as Workers minus Soldiers: 2$^{nd}$ instars, $t = -3.095$, df = 7.73; 3$^{rd}$ instars (day 1), $t = -3.970$, df = 5.53; 3$^{rd}$ instars (day 5), $t = -7.134$, df = 4.06; 3$^{rd}$ instars (day 9), $t = -4.706$, df = 5.04; Prepupae, $t = -8.175$, df = 4.38; Pupae, $t = -10.365$, df = 4.65. **(e–h)** Molecular evolutionary analyses of the *torch* coding sequence across hymenopteran orthologs. **(e)** Branch-site tests mapping positive selection (omega = 3.11, $P = 0.0026$; Supplementary Dataset 2) to the ancestral ant lineage, coincident with a shorter terminal coding exon in ants than in the more closely related non-ant hymenopteran lineages sampled (gene models towards the right showing coding sequence in magenta and introns in black). Branches in the gene tree with omegas higher than 3.5 were gray-shaded, highlighting an arbitrary threshold for denoting cases where branch-wise *d*S approaches zero so that omega = *d*N/*d*S increases toward infinity and becomes uninterpretable; such near-zero *d*S values typically arise on very short, low-divergence branches, so these large omega ratios should not be interpreted as evidence for positive selection [15,16]. **(f)** The matching *d*N tree of panel e where branch lengths are proportional to the estimated number of nonsynonymous substitutions per nonsynonymous site (*d*N) to highlight lineages with elevated rates of amino-acid change. **(g)** The same *d*S tree but with branch lengths proportional to the estimated number of synonymous substitutions per synonymous site (*d*S), providing a neutral baseline substitution-rate for comparison with panel f. **(h)** A tree topology with internal node labels to facilitate cross-referencing between in panels f-g and panel e. Node 27, representing the ancestral ant lineage (Supplementary Dataset 2), is highlighted in red. Scale bars indicate expected substitutions per site. The contrast between panels f and g facilitates visualizing branches where amino-acid changes (*d*N) were disproportionately high relative to synonymous changes (*d*S), consistent with the positive-selection signal in the ancestral ant lineage in panel e. **(i)** RNA-FISH for *torch* mRNA in late third-instar *M. pharaonis* larval adipocytes. The *torch* signal is shown in red and nuclei are counterstained in gray. JH-treated workers exhibited increased *torch* signaling relative to control workers, reaching levels comparable to gynes. Scale bars, 10 μm.

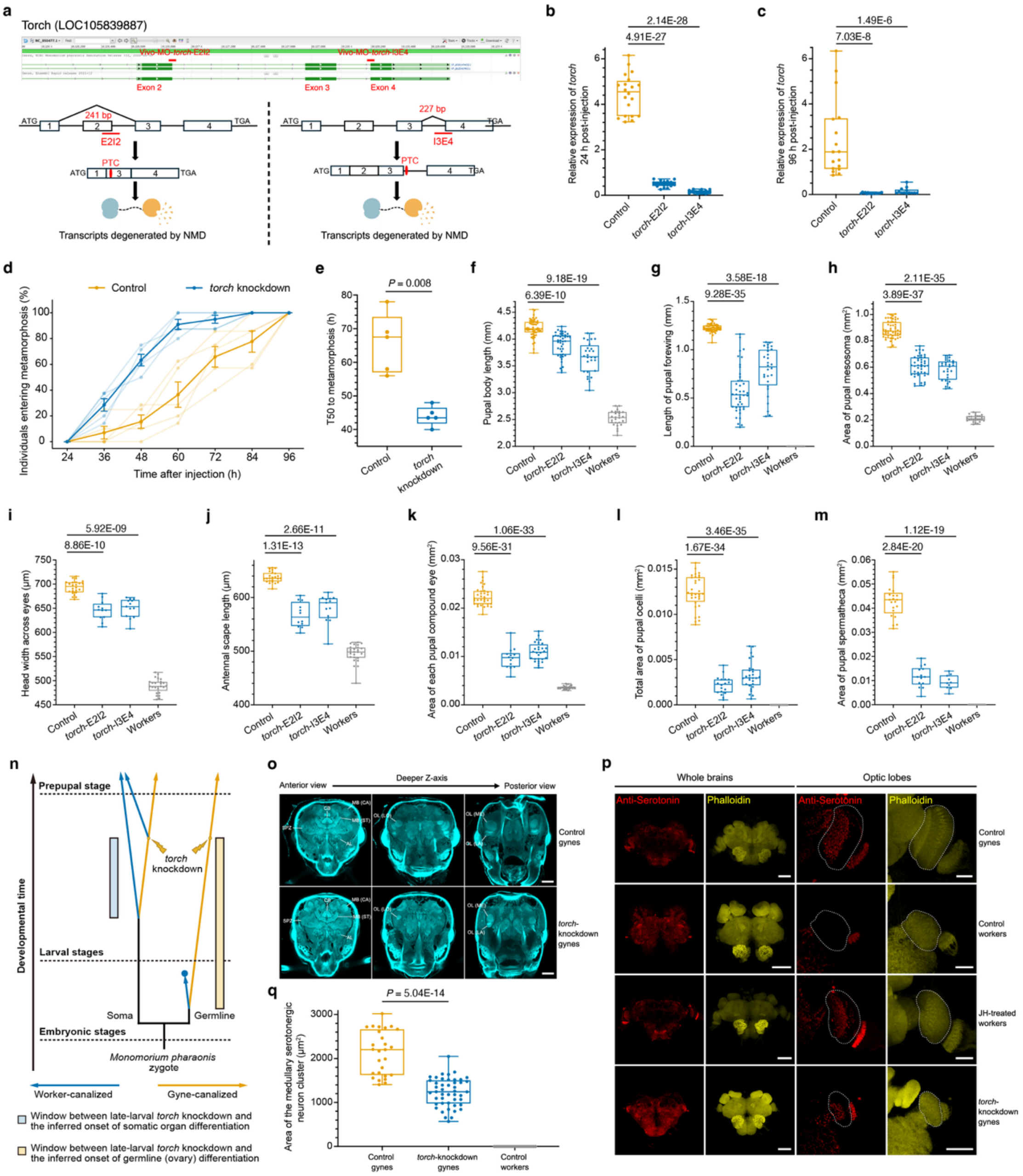

a
Torch (LOC105839887)
Vivo-MO-torch-E2I2
Vivo-MO-torch-I3E4
Exon 2
Exon 3
Exon 4
241 bp
227 bp
E2I2
I3E4
PTC
ATG
TGA
Transcripts degenerated by NMD
b
2.14E-28
4.91E-27
Relative expression of torch 24 h post-injection
Control
torch-E2I2
torch-I3E4
c
1.49E-6
7.03E-8
Relative expression of torch 96 h post-injection
d
Control
torch knockdown
Individuals entering metamorphosis (%)
Time after injection (h)
e
P = 0.008
T50 to metamorphosis (h)
torch knockdown
f
9.18E-19
6.39E-10
Pupal body length (mm)
Workers
g
3.58E-18
9.28E-35
Length of pupal forewing (mm)
h
2.11E-35
3.89E-37
Area of pupal mesosoma (mm²)
i
5.92E-09
8.86E-10
Head width across eyes (μm)
j
2.66E-11
1.31E-13
Antennal scape length (μm)
k
1.06E-33
9.56E-31
Area of each pupal compound eye (mm²)
l
3.46E-35
1.67E-34
Total area of pupal ocelli (mm²)
m
1.12E-19
2.84E-20
Area of pupal spermatheca (mm²)
n
Prepupal stage
Larval stages
Embryonic stages
Developmental time
torch knockdown
Soma
Germline
Monomorium pharaonis zygote
Worker-canalized
Gyne-canalized
Window between late-larval torch knockdown and the inferred onset of somatic organ differentiation
Window between late-larval torch knockdown and the inferred onset of germline (ovary) differentiation
o
Anterior view
Deeper Z-axis
Posterior view
Control gynes
torch-knockdown gynes
q
P = 5.04E-14
Area of the medullary serotonergic neuron cluster (μm²)
Control gynes
torch-knockdown gynes
Control workers
p
Whole brains
Optic lobes
Anti-Serotonin
Phalloidin
Control gynes
Control workers
JH-treated workers
torch-knockdown gynes

**Extended Data Fig. 2. Vivo-MO-mediated *torch* knockdown redirects gyne development toward worker-like somatic and neural phenotypes. (a)** Targeted junctions of splice-blocking vivo-MOs in *torch* pre-mRNA. Detailed design information is provided in the section "Design and validation of splice-blocking vivo-morpholinos (vivo-MOs) targeting the *torch* pre-mRNA" in the Supplementary Information. PTC: premature termination codon; NMD: nonsense-mediated decay. **(b-c)** ddPCR 24 h after vivo-MO injection (b; late third-instar larvae) and 96h after vivo-MO injection (c; prepupae) showing strong depletion of *torch* in splice-blocking vivo-morpholino-treated gynes (E2I2- and I3E4-targeting vivo-MOs) relative to control gynes. Statistics: one-way ANOVA followed by Dunnett's multiple comparisons tests. (b) $n$ = 20, 17 and 16 samples for control, *torch*-E2I2 and *torch*-I3E4, respectively; one-way ANOVA: $F(2, 50) = 357.21$, $P$ = 2.46E-30. (c) $n$ = 17, 16 and 13 samples for control, *torch*-E2I2 and *torch*-I3E4, respectively; one-way ANOVA: $F(2, 43) = 26.928$, $P$ = 2.62E-8. Dunnett-adjusted $P$ values are indicated in the figure. **(d-e)** Knockdown of *torch* accelerates the onset of metamorphosis: (d) Cumulative progression to metamorphosis in gyne larvae from 24 to 96 h after vivo-MO injection. Pale lines and points represent individual independent experiments, whereas darker lines and points represent group means with SEM; *torch*-knockdown larvae entered metamorphosis earlier than controls, showing a faster increase in cumulative proportion of prepupae across the observation window (five independent experiments per group). (e) Replicate-level T50 values, defined as the interpolated time at which 50% of larvae in each independent experiment had entered the prepupal stage; knockdown of *torch* significantly reduced T50 compared with controls (44.0 ± 2.9 h versus 65.7 ± 8.9 h; mean ± SD; $n$ = 5 independent experiments per group; exact permutation test; two-sided $P$ value is indicated in the figure). **(f–m)** Knockdown of *torch* in gynes reduced body size and produced worker-like growth of imaginal-disc-derived organs in day-12 pupae: (f) body length; (g) forewing length; (h) mesosoma area; (i) head width across the eyes; (j) antennal scape length; (k) compound-eye area per eye; (l) total ocellar area; (m) spermathecal area; *torch*-E2I2 and *torch*-I3E4 indicate *torch*-knockdown gynes injected with E2I2- and I3E4-targeting vivo-MOs, respectively, relative to controls. Workers are included as a reference to illustrate the worker-biased reduction in organ and body size observed in *torch*-knockdown gynes; worker values in panels g, l and m were always zero because forewings, ocelli and the spermatheca are absent in workers [12,17]. Statistics: one-way ANOVA followed by Dunnett's multiple-comparisons tests. Dunnett-adjusted $P$ values are indicated in the figure. Sample sizes are given in the order control, *torch*-E2I2 and *torch*-I3E4: (f) $n$ = 53, 43 and 28; $F(2, 121) = 60.82$, $P$ = 5.24E−19. (g) $n$ = 53, 43 and 28; $F(2, 121) = 161.26$, $P$ = 7.41E−35. (h) $n$ = 53, 43 and 28; $F(2, 121) = 239.63$, $P$ = 8.30E−43. (i) $n$ = 24, 12 and 13; $F(2, 46) = 43.19$, $P$ = 2.76E−11. (j) $n$ = 24, 12 and 13; $F(2, 46) = 71.90$, $P$ = 6.96E−15. (k) $n$ = 33, 14 and 28; $F(2, 72) = 327.19$, $P$ = 7.28E−37. (l) $n$ = 32, 18 and 28; $F(2, 75) = 360.85$, $P$ = 3.28E−39. (m) $n$ = 21, 14 and 10; $F(2, 42) = 208.52$, $P$ = 1.55E−22. **(n)** Model for the relative preservation of ovaries after late knockdown of *torch* in *M. pharaonis*. Previous work in this ant demonstrated that germline and somatic caste differentiation are temporally asymmetric: worker-destined embryos completely lose primordial germ cells during early embryogenesis, and late worker larval JH treatment induces multiple gyne-like somatic traits but without restoring ovary development [12,17]. We therefore hypothesize that, by the time knockdown of *torch* was initiated in late third-instar gynes, ovarian developmental potential was already more strongly canalized than that of somatic organs. This asymmetry is illustrated by the longer interval between late-larval knockdown of *torch* and the inferred onset of germline (ovary) differentiation than between knockdown of *torch* and the inferred onset of somatic organ differentiation. Accordingly, late depletion of *torch* strongly disrupted gyne-canalized somatic development, reducing organ size and redirecting these organs toward worker-like trajectories, while leaving ovaries relatively preserved. Because many ants retain worker reproductive plasticity [18,19], this interpretation may reflect a derived condition in *M. pharaonis* rather than a universal rule across ant lineages so that *torch* may still contribute to ovary development from the larval stage in species whose workers retain a germline. Figure adapted from previous work [12]. **(o)** Representative confocal optical sections at different z-depths through annotated brain compartments: MB, mushroom bodies; CA, calyx; ST, stalk; AL, antennal lobes; CB, central body; SPZ, supraesophageal zone; OL, optic lobes; LO, lobula; ME, medulla; LA, lamina. The MB comprise the CA and ST, whereas the OL comprise the LO, ME, and LA. These image stacks were used for 3D reconstruction and neuropil-volume quantification using Amira software, as shown in Fig. 2g-h. **(p)** Day-12 pupal brains stained with an anti-serotonin antibody to label serotonergic neurons (red) and counterstained with Alexa Fluor™ 546–conjugated phalloidin to label F-actin (yellow). Left, whole brain; right, enlarged view of the optic lobe. In each optic-lobe enlargement, the medulla is outlined by a white dashed line, and the red immunofluorescence signal within this region indicates the medullary serotonergic neuron cluster. This cluster is normally present in the medulla of pupal gynes but absent in workers (top two rows). Larval JH treatment restores the cluster in workers (third row), whereas the cluster is markedly reduced in *torch*-knockdown gynes (bottom row). **(q)** Quantification of the anti-serotonin-positive area showing the reduced size of the medullary serotonergic neuron cluster in *torch*-knockdown gynes relative to control gynes. Worker values are consistently zero because this neuronal cluster is absent in workers. Each dot represents one medulla. Statistical significance of differences between control gynes and *torch*-knockdown gynes was assessed using a two-sided Student's unpaired $t$-test: $n = 27$ medullae from 14 control gynes and $n = 46$ medullae from 23 *torch*-knockdown gynes; $t = 9.362$, df = 71. Scale bars, 100 μm (o), 100 μm (whole-brain images in p) and 30 μm (optic-lobe enlargements in p).

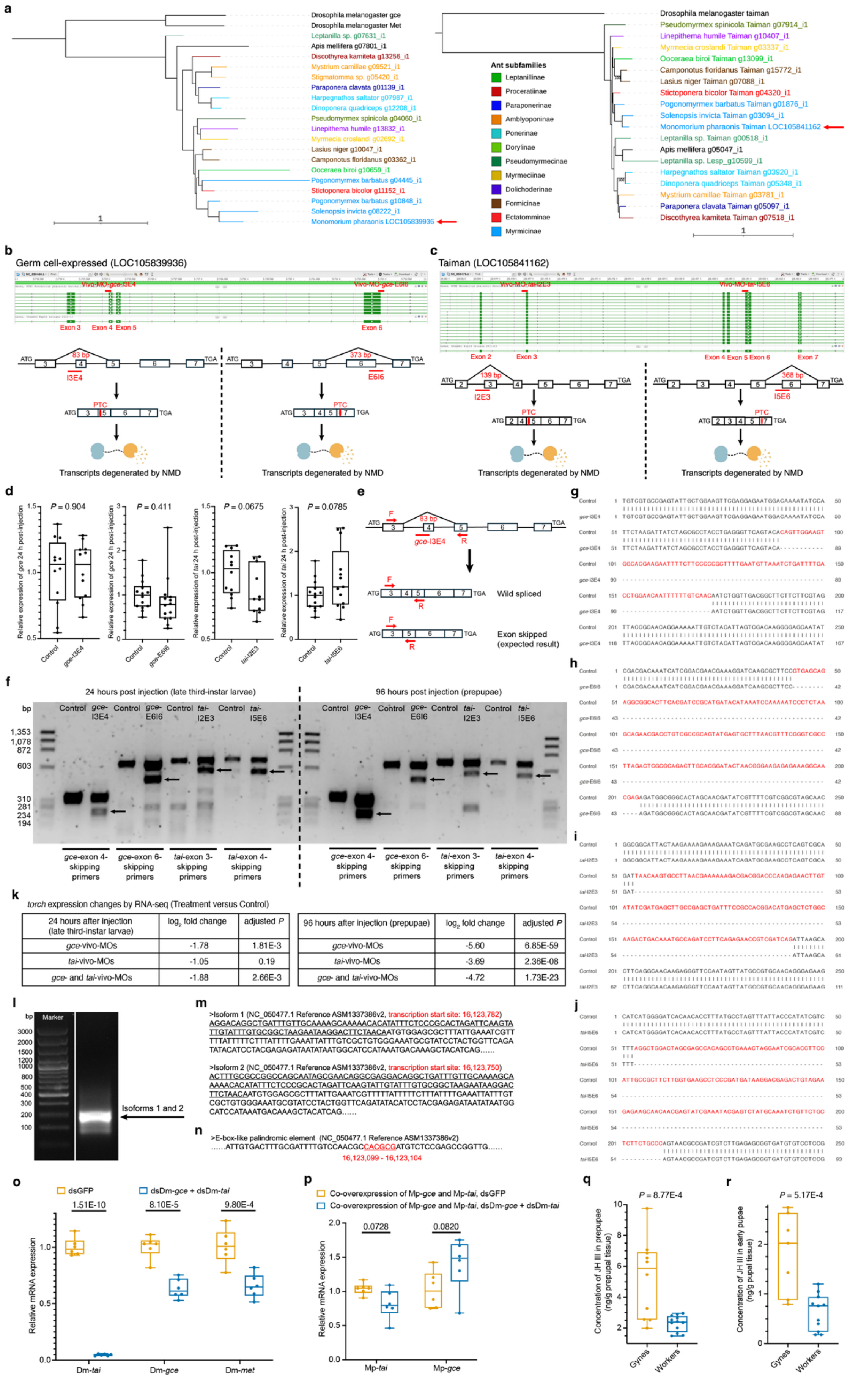


| 24 hours after injection (late third-instar larvae) | $log_2$ fold change | adjusted P |
|---|---|---|
| gce-vivo-MOs | -1.78 | 1.81E-3 |
| tai-vivo-MOs | -1.05 | 0.19 |
| gce- and tai-vivo-MOs | -1.88 | 2.66E-3 |

| 96 hours after injection (prepupae) | $log_2$ fold change | adjusted P |
|---|---|---|
| gce-vivo-MOs | -5.60 | 6.85E-59 |
| tai-vivo-MOs | -3.69 | 2.36E-08 |
| gce- and tai-vivo-MOs | -4.72 | 1.73E-23 |

**Extended Data Fig. 3. Gce–Tai perturbation and promoter assays establish direct JH-dependent activation of *torch*. (a)** Phylogenetic gene trees of Gce/Met (left) and Tai (right) with the ant branches colored by subfamily and the outgroups *Drosophila melanogaster* and *Apis mellifera* having black branches. The orthologs in *Monomorium pharaonis* are marked with red arrows. **(b)** The *gce* locus (LOC105839936) with genome-browser tracks indicating junctions targeted by Vivo-MO-*gce*-I3E4 (intron 3-exon 4) and Vivo-MO-*gce*-E6I6 (exon 6-intron 6), and diagrams illustrating predicted consequence-frameshifts or a premature termination codon (PTC) and nonsense-mediated decay (NMD) for I3E4 and E6I6. **(c)** The *tai* locus (LOC105841162) targeted by Vivo-MO-*tai*-I2E3 (intron 2-exon 3) and Vivo-MO-*tai*-I5E6 (intron 5-exon 6) and thus predicted to induce exon skipping, frameshifts, and PTCs leading to NMD. **(d-j)** Splice-blocking vivo-MOs induce exon skipping in *gce* and *tai* without triggering NMD. **(d)** ddPCR 24 h post-injection of late third instar gyne larvae showed no significant reduction of total *gce* or *tai* mRNA when quantified with primers outside the targeted exons, indicating that the splice-blocking vivo-MOs did not detectably induce transcript-wide NMD. Sample sizes were $n$ = 12/12 for *gce*-I3E4, $n$ = 12/11 for *tai*-I2E3, $n$ = 14/14 for *gce*-E6I6 and $n$ = 14/13 for *tai*-I5E6, with control/experimental samples listed in that order. Statistical significance was assessed using two-sided Student's unpaired $t$-tests: *gce*-I3E4, $t = -0.122$, df = 22; *tai*-I2E3, $t = 1.928$, df = 21; *gce*-E6I6, $t = 0.836$, df = 26; *tai*-I5E6, $t = -1.835$, df = 25. $P$ values are indicated in the figure. **(e)** Primer design for exon-spanning PCR (example: *gce*-I3E4), used to obtain direct evidence for exon skipping. **(f)** Agarose gels 24 h (late third instar gyne larvae) and 96 h (prepupae) post-injection revealed shorter amplicons in MO-treated samples (arrows), consistent with exon skipping. **(g–j)** Sanger sequencing confirmed complete or partial exon skipping in the *gce*-I3E4, *gce*-E6I6, *tai*-I2E3 and *tai*-I5E6 vivo-MO treatment groups, validating the predicted junction-specific splice alterations. Nucleotide segments absent from the vivo-MO–induced splice products relative to the wild-type spliced transcripts are highlighted in red. The corresponding exon identities of these skipped segments are described in the section "Design and validation of splice-blocking vivo-MOs targeting the *gce* pre-mRNA and *tai* pre-mRNA of the JH receptor complex" in the Supplementary Information. Raw Sanger sequencing files are provided in Supplementary Dataset 5. **(k-p)** JH–Gce–Tai signaling directly targets *torch* via a promoter E-box–like element. **(k)** RNA-seq–based expression changes of *torch* following loss of Gce, Tai, or Gce+Tai function using values extracted from Supplementary Dataset 6. **(l)** 5'-RACE products from gyne broods revealed two similarly sized *torch* 5'-UTR isoforms. **(m)** Sanger sequencing identified two transcription start sites (TSSs) for *torch* at chr11:16,123,782 and chr11:16,123,750 on NC_050477.1, thereby defining the precise transcription initiation positions of *torch*. Underlined sequences indicate the 5' UTR region, and the raw Sanger sequencing data are provided in Supplementary Dataset 7. **(n)** Inspection of the promoter window (-3 kb to +1 kb) revealed a single E-box-like palindromic element, CACGCG, at 16,123,099-16,123,104 (~650 bp upstream of both TSSs), consistent with a Gce-Tai recognition site and supporting direct JH receptor regulation of *torch*. **(o)** dsRNA knockdown efficiency for endogenous *D. melanogaster met*, *gce*, and *tai* in S2 cells, used to reduce background JH-receptor activity. Because *gce* and *met* are paralogs and thus share substantial sequence similarity [20], dsRNAs designed against *gce* and *tai* efficiently reduced the expression of all three genes simultaneously. Statistics: multiple two-sided Student's unpaired $t$-tests followed by Holm–Šídák correction across the three comparisons. $n$ = 6 samples per condition for each readout. Dm-*met*, $t = 4.600$, df = 10; Dm-*gce*, $t = 6.929$, df = 10; Dm-*tai*, $t = 29.291$, df = 10. **(p)** Relative expression of co-transfected *M. pharaonis gce* and *tai* in

S2 cells, with or without dsRNAs targeting *D. melanogaster gce* and *tai*, showing that these dsRNAs appear to not alter *M. pharaonis gce* or *tai* expression. Statistics: multiple two-sided Student's unpaired $t$-tests followed by Holm–Šídák correction across the two comparisons. $n = 6$ samples per condition for each readout. Mp-*tai*, $t = 2.404$, df = 10; Mp-*gce*, $t = -1.933$, df = 10. Holm–Šídák-adjusted $P$ values are indicated in panels o-p. Dm, *D. melanogaster*; Mp, *M. pharaonis*. **(q–r)** JH III concentrations in whole prepupae (q) and early pupae (r), measured by targeted LC–MS. At both stages, gynes exhibited significantly higher JH III levels than workers, consistent with the pattern observed in third-instar larvae (Fig. 3c). Notably, JH III concentrations were higher in prepupae than in third-instar larvae, consistent with reports of a prepupal rise in JH titers in the superorganismally caste-differentiated honeybees, bumble bees, and stingless bees [21-24]. Sample sizes were: (q) prepupae, $n = 10$ gynes and $n = 12$ workers; (r) early pupae, $n = 7$ gynes and $n = 11$ workers. Statistical significances were assessed using two-sided Student's unpaired $t$-tests: (q) $t = 3.906$, df = 20; (r) $t = 4.330$, df = 16.

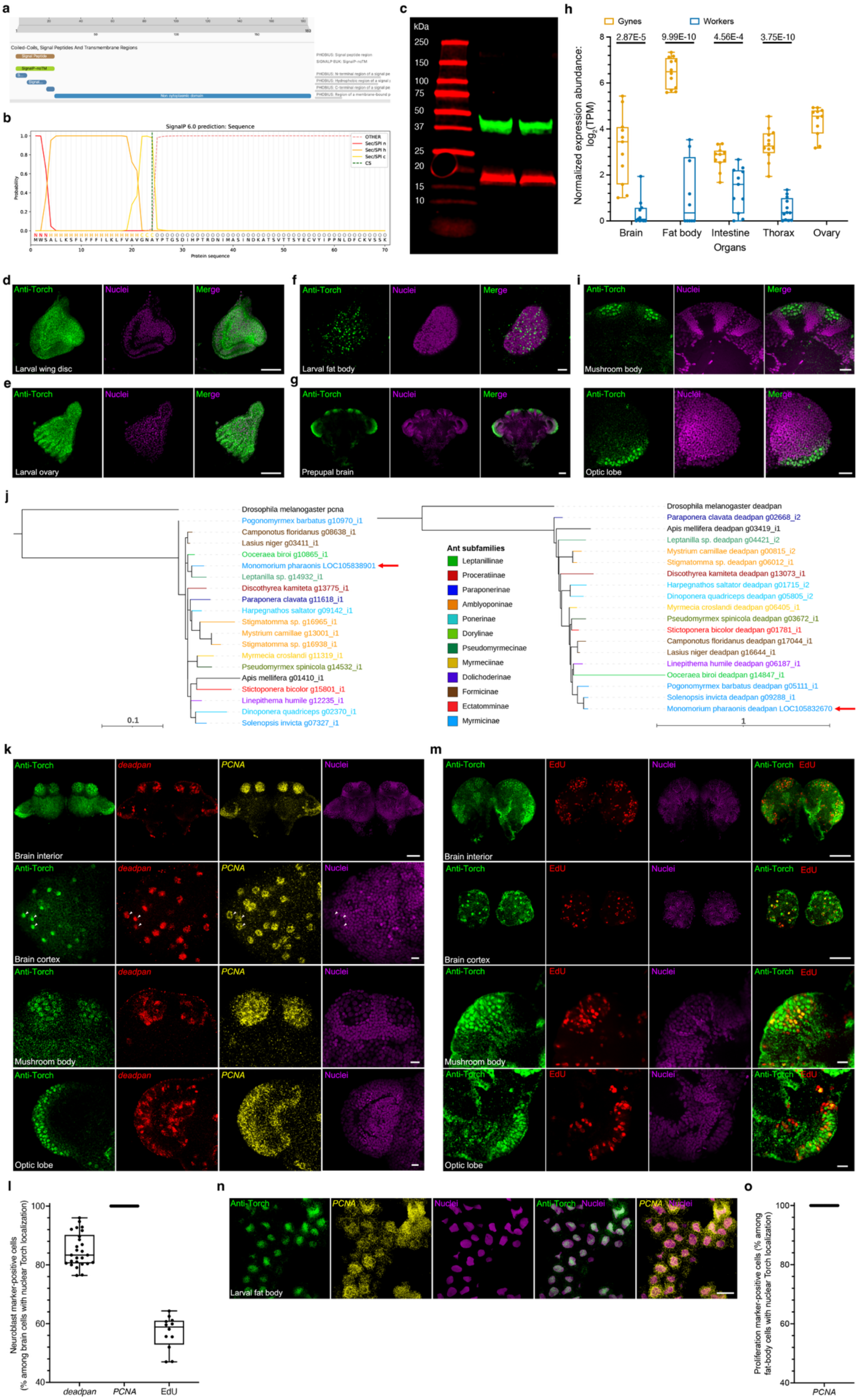
a
Coiled-Coils, Signal Peptides And Transmembrane Regions
b
SignalP 6.0 prediction: Sequence
OTHER
Sec/SPI n
Sec/SPI h
Sec/SPI c
CS
Probability
Protein sequence
c
kDa
250
150
100
75
50
37
25
20
15
10
h
Gynes
Workers
2.87E-5
9.99E-10
4.56E-4
3.75E-10
Normalized expression abundance: log2(TPM)
Brain
Fat body
Intestine
Thorax
Ovary
Organs
d
Anti-Torch
Nuclei
Merge
Larval wing disc
e
Larval ovary
f
Larval fat body
g
Prepupal brain
i
Mushroom body
Optic lobe
j
Drosophila melanogaster pcna
Pogonomyrmex barbatus g10970_i1
Camponotus floridanus g08638_i1
Lasius niger g03411_i1
Ooceraea biroi g10865_i1
Monomorium pharaonis LOC105838901
Leptanilla sp. g14932_i1
Discothyrea kamiteta g13775_i1
Paraponera clavata g11618_i1
Harpegnathos saltator g09142_i1
Stigmatomma sp. g16965_i1
Mystrium camillae g13001_i1
Stigmatomma sp. g16938_i1
Myrmecia croslandi g11319_i1
Pseudomyrmex spinicola g14532_i1
Apis mellifera g01410_i1
Stictoponera bicolor g15801_i1
Linepithema humile g12235_i1
Dinoponera quadriceps g02370_i1
Solenopsis invicta g07327_i1
0.1
Ant subfamilies
Leptanillinae
Proceratiinae
Paraponerinae
Amblyoponinae
Ponerinae
Dorylinae
Pseudomyrmecinae
Myrmeciinae
Dolichoderinae
Formicinae
Ectatomminae
Myrmicinae
Drosophila melanogaster deadpan
Paraponera clavata deadpan g02668_i2
Apis mellifera deadpan g03419_i1
Leptanilla sp. deadpan g04421_i2
Mystrium camillae deadpan g00815_i2
Stigmatomma sp. deadpan g06012_i1
Discothyrea kamiteta deadpan g13073_i1
Harpegnathos saltator deadpan g01715_i2
Dinoponera quadriceps deadpan g05805_i2
Myrmecia croslandi deadpan g06405_i1
Pseudomyrmex spinicola deadpan g03672_i1
Stictoponera bicolor deadpan g01781_i1
Camponotus floridanus deadpan g17044_i1
Lasius niger deadpan g16644_i1
Linepithema humile deadpan g06187_i1
Ooceraea biroi deadpan g14847_i1
Pogonomyrmex barbatus deadpan g05111_i1
Solenopsis invicta deadpan g09288_i1
Monomorium pharaonis deadpan LOC105832670
1
k
Anti-Torch
deadpan
PCNA
Nuclei
Brain interior
Brain cortex
Mushroom body
Optic lobe
m
Anti-Torch
EdU
Nuclei
Anti-Torch EdU
Brain interior
Brain cortex
Mushroom body
Optic lobe
l
Neuroblast marker-positive cells (% among brain cells with nuclear Torch localization)
100
80
60
40
deadpan
PCNA
EdU
n
Anti-Torch
PCNA
Nuclei
Anti-Torch Nuclei
PCNA Nuclei
Larval fat body
o
Proliferation marker-positive cells (% among fat-body cells with nuclear Torch localization)
100
80
60
40
PCNA

**Extended Data Fig. 4. Torch accumulates in nuclei of proliferating progenitors across developing gyne tissues. (a, b)** Predicted features of the *M. pharaonis* Torch protein, including (a) InterPro annotations and (b) the N-terminal signal peptide predicted by SignalP 6.0. An N-terminal Sec/SPI signal peptide, predicted to span residues 1–24, is the only recognizable structural feature of the 183-aa Torch protein. **(c)** Western blot analysis of gyne brood lysates probed with rabbit anti-Torch antibody detected a single band at ~18 kDa (red), corresponding to the predicted molecular mass of mature Torch after signal-peptide cleavage, supporting antibody specificity. Mouse anti-β-actin was co-probed as a loading control (~42 kDa; green). Two biological replicates of gyne brood lysates are shown. **(d, e)** Immunofluorescence for Torch (green) in third-instar wing imaginal discs of gyne larvae (d) and ovaries (e), with nuclei counterstained in magenta. **(f)** High-magnification anti-Torch immunofluorescence in fat bodies of third-instar gyne larvae, with anti-Torch signal in green and nuclei counterstained in magenta, revealing nuclear enrichment of endogenous Torch and its organization in discrete condensates, consistent with the liquid–liquid phase–separated condensates described for many nuclear transcription factors. **(g)** Anti-Torch immunofluorescence in brains of gyne prepupae in green and nuclei are counterstained in magenta, showing strong Torch expression throughout the brain, particularly in the mushroom bodies and optic lobes. **(h)** RNA-seq–based tissue expression of *torch* in day-1 adults, showing *torch* transcript abundance is gyne-biased across brain, fat body, intestine and thorax; ovary expression is shown only for gynes because *M. pharaonis* workers lack ovaries [12]. Sample sizes were: brain, $n = 11$ gynes and $n = 11$ workers; fat body, $n = 12$ gynes and $n = 10$ workers; intestine, $n = 11$ gynes and $n = 11$ workers; thorax, $n = 12$ gynes and $n = 12$ workers. Statistics: multiple two-sided Student's unpaired *t*-tests followed by Holm–Šídák correction across the four gyne–worker tissue comparisons: brain, $t = 5.692$, df = 20; fat body, $t = 11.403$, df = 20; intestine, $t = 4.186$, df = 20; thorax, $t = 11.473$, df = 22. Holm–Šídák-adjusted $P$ values are indicated in the figure. **(i)** Higher-magnification anti-Torch immunofluorescence of mushroom bodies (upper panels) and optic lobes (lower panels) of gyne prepupae in green and nuclei counterstained in magenta. Cells with prominent nuclear Torch accumulation displayed enlarged, weakly DNA-stained nuclei relative to neighboring cells, a classic neuroblast morphology consistent with high cell proliferation [25]. **(j)** Phylogenetic gene trees of the neuroblast markers *PCNA* (left) and *deadpan* (right). Additional legend details are provided in Extended Data Fig. 3a. **(k)** Immunofluorescence for Torch (green) in brains of gyne prepupae combined with RNA-FISH for *PCNA* (yellow) and *deadpan* (red) with nuclei counterstained in magenta. Top row: cross-sections showing all three signals concentrated in mushroom bodies and optic lobes. Second to fourth rows: higher-magnification views of the superficial brain cortex, mushroom bodies, and optic lobes, showing that nuclei with Torch accumulation are largely *PCNA*- and *deadpan*-positive, consistent with neuroblast identity. White arrowheads mark three representative neuroblasts in the superficial brain cortex with co-localized nuclear Torch, *deadpan* and *PCNA* signals. **(l)** Quantification of the percentage of Torch-positive brain nuclei that were also positive for *deadpan*, *PCNA* or EdU. All Torch-positive nuclei were *PCNA*-positive (100%), whereas 83% were *deadpan*-positive and 60% incorporated EdU, indicating DNA synthesis during the S phase. For *deadpan* and *PCNA*, 1,385 Torch-positive nuclei from 26 confocal fields across gyne brains from four third-instar larvae and four prepupae were analyzed. For EdU, 1,232 Torch-positive nuclei from 12 confocal fields across five third-instar larval brains of gynes were analyzed. Each dot represents the percentage calculated from one confocal field. **(m)** EdU incorporation (red) combined with Torch immunofluorescence (green) in brains of third-instar gyne larvae, with nuclei counterstained in magenta. Top row: cross-sections showing EdU and Torch signals concentrated in mushroom bodies and optic lobes. Second to fourth rows: higher-magnification views of the superficial brain cortex, mushroom bodies, and optic lobes, indicating that a subset of Torch-positive brain cells is actively synthesizing DNA. **(n)** Anti-Torch immunofluorescence (green) combined with RNA-FISH for *PCNA* (yellow) in fat bodies of third-instar gyne larvae, with nuclei counterstained in magenta. All fat-body cells showed nuclear Torch accumulation and *PCNA* signal. **(o)** Quantification of the percentage of Torch-positive fat-body nuclei that were *PCNA*-positive in third-instar gyne larvae. All Torch-positive fat-body nuclei were *PCNA*-positive (100%; n = 867 Torch-positive nuclei quantified from 22 confocal fields across five third-instar gyne larvae). Each dot represents the percentage calculated from one confocal field. Scale bars: 50 µm in d, e, g, k top row, m first and second rows, and n; 20 µm in i; 10 µm in k second to fourth rows, and in m third and fourth rows; 5 µm in f.

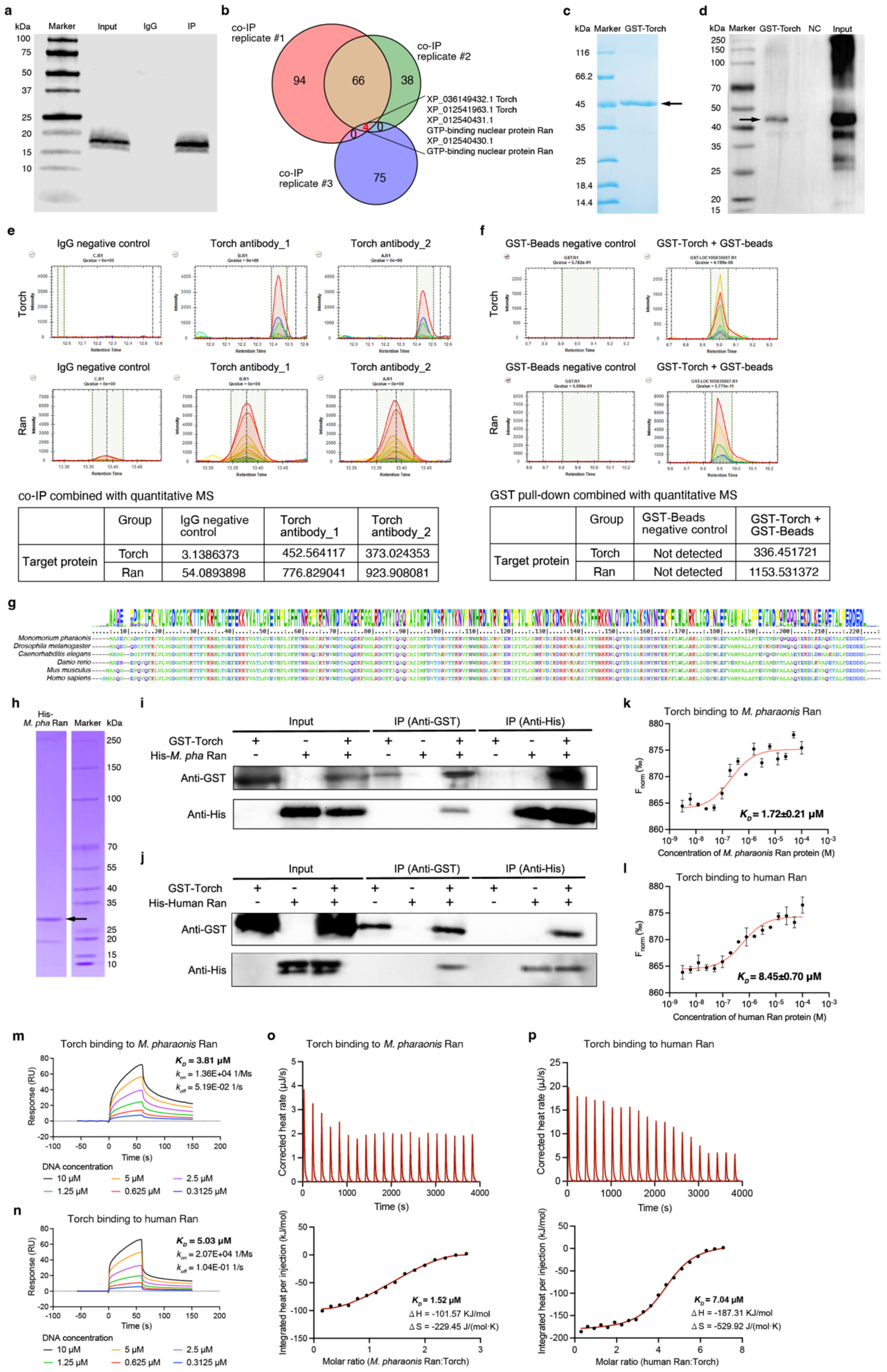


co-IP combined with quantitative MS

| | Group | IgG negative control | Torch antibody_1 | Torch antibody_2 |
|---|---|---|---|---|
| Target protein | Torch | 3.1386373 | 452.564117 | 373.024353 |
| | Ran | 54.0893898 | 776.829041 | 923.908081 |

GST pull-down combined with quantitative MS

| | Group | GST-Beads negative control | GST-Torch + GST-Beads |
|---|---|---|---|
| Target protein | Torch | Not detected | 336.451721 |
| | Ran | Not detected | 1153.531372 |

**Extended Data Fig. 5. Torch directly binds the deeply conserved Ran GTPase. (a)** Western blot verification of the co-immunoprecipitation (co-IP) assay using whole-body lysates from gyne brood of *M. pharaonis*, immunoprecipitated with anti-Torch or control IgG, with an aliquot of the starting lysate collected before antibody incubation as the input control. Blotting with anti-Torch detected a single ~18 kDa band (predicted mass of mature Torch, 18.7 kDa) in the anti-Torch IP and input lanes, but not in the IgG control, confirming antibody specificity and successful enrichment of Torch. **(b)** Venn diagram of proteins uniquely identified in anti-Torch IPs (i.e., absent from IgG controls) across three independent co-immunoprecipitation (co-IP) replicates by qualitative mass spectrometry (MS). Aside from Torch (LOC105839887) itself, GTP-binding nuclear protein Ran was the only candidate protein recovered in all three replicates. Complete lists of proteins uniquely detected in each anti-Torch IP replicate are provided in Supplementary Dataset 8. **(c)** Production of recombinant GST-tagged Torch verified by SDS-PAGE (predicted molecular mass: 46.3 kDa; band highlighted). **(d)** Western blot verification of the GST-Torch pull-down, incubating gyne brood lysates with GST-Torch (lane "GST-Torch") or GST beads alone (negative control, "NC") while retaining an aliquot as input. Blots probed with anti-GST (Catalog number: Proteintech 10000-0-AP) revealed a prominent ~45 kDa GST-Torch band (highlighted by an arrow) in the bait and input lanes but not in the NC lane, confirming bait capture and assay specificity. **(e-f)** Parallel-reaction-monitoring (PRM) MS quantification showed pronounced Ran enrichment in Torch pulldowns relative to controls in the co-IP (e) and the GST pull-down (f) samples. Upper panel: PRM extracted-ion chromatograms (XICs) for representative peptides of Torch in the experimental pull-down (bait) and the negative control. Middle panel: PRM XICs for representative peptides of Ran in the same samples. Lower panel: quantitative protein abundances for Torch and Ran across sample categories – computed by aggregating the intensities of all quantified peptides per protein – illustrating enrichment in the experimental pull-down relative to the control. Two independent co-IP replicates of the Torch pull-downs were performed, indicated as "Torch antibody_1" and "Torch antibody_2". **(g)** Multiple sequence alignment of Ran proteins from *Monomorium pharaonis* and other model organisms illustrated strong conservation across metazoans. **(h)** SDS-PAGE confirming production of recombinant His-tagged *M. pharaonis* Ran (predicted molecular mass: 26 kDa; major band highlighted). **(i-j)** Reciprocal IPs of purified proteins, showing direct association between GST-Torch and His-*M. pharaonis* Ran (i) and human (j) Ran, established a direct Torch-Ran interaction that is conserved between ants and humans. Complexes were pulled down with anti-GST or anti-His and immunoblotted with the reciprocal antibody, with inputs shown for reference. **(k-l)** Microscale thermophoresis (MST) assays corroborated these interactions, yielding dissociation constants $K_D$ = 1.72 ± 0.21 µM for *M. pharaonis* Ran (k) and $K_D$ = 8.45 ± 0.70 µM for human Ran (l). **(m-n)** Surface plasmon resonance (SPR) assays revealed binding between Torch and *M. pharaonis* Ran (m; $K_D$ = 3.81 µM) and human Ran (n; $K_D$ = 5.03 µM), as specified in resonance units (RU). **(o-p)** Isothermal titration calorimetry (ITC) assays demonstrated direct binding of Torch to *M. pharaonis* Ran (o; $K_D$ = 1.52 µM) and human Ran (p; $K_D$ = 7.04 µM).

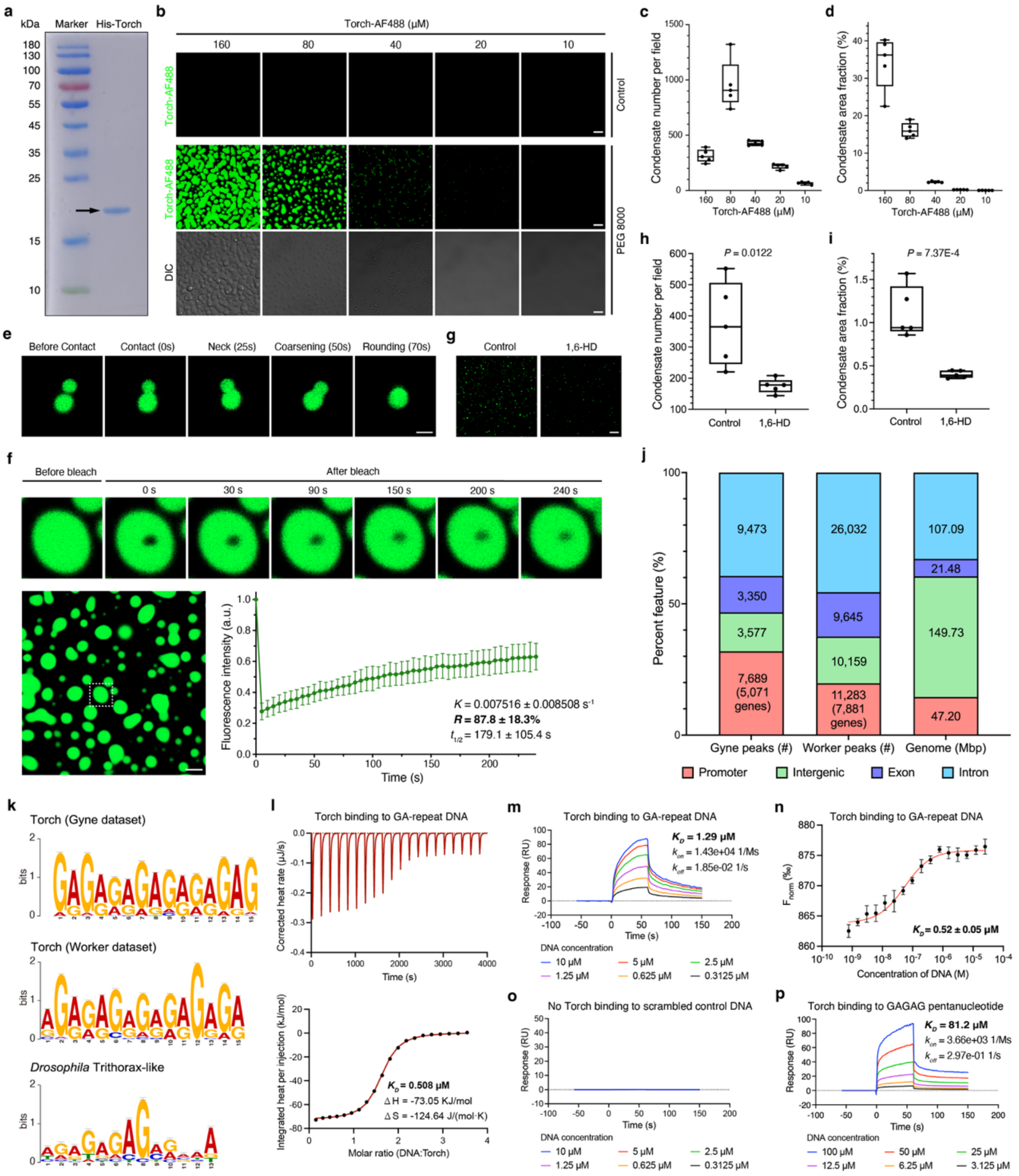

**Extended Data Fig. 6. Torch forms dynamic condensates and directly recognizes GA-repeat DNA. (a)** SDS-PAGE verification of purified *M. pharaonis* His-Torch recombinant protein (predicted molecular mass, 19.6 kDa; arrow), used for the *in vitro* liquid-liquid phase separation (LLPS), ITC, SPR, and MST assays. **(b)** Representative fluorescence and differential interference contrast (DIC) images of Alexa Fluor 488-labeled His-Torch (Torch-AF488) at the indicated concentrations (160, 80, 40, 20, and 10 μM). Samples were incubated at 4°C overnight either without PEG 8000 (Control) or with 10% PEG 8000. No condensates were detected in the no-PEG control, whereas 10% PEG 8000 induced concentration-dependent Torch condensate formation. **(c-d)** Quantification of condensate number per field (c) and condensate area fraction (d) from images as in (b). Condensate number peaked at 80 μM and declined at 160 μM, consistent with extensive coalescence at the highest Torch concentration; numbers also declined at lower concentrations where nucleation became limiting. Condensate area fraction, defined as total area divided by the imaging field, increased with Torch concentration, supporting concentration-driven condensate growth and coarsening; n = 5 independent fields per concentration. **(e)** Time-lapse images of condensate fusion in 160 μM Torch-AF488 with 10% PEG 8000 after 0.5–1 h incubation. Two contacting droplets formed a neck, coarsened, and rounded into a single spherical condensate within 70 s, indicating liquid-like surface behavior. **(f)** FRAP analysis of Torch-alone condensates formed with 160 μM Torch-AF488 and 10% PEG 8000 after 0.5–1 h incubation. The white dashed box marks the condensate enlarged for bleaching. Fluorescence intensity of the bleached ROI was corrected to an unbleached reference ROI after background subtraction and normalized to the mean pre-bleach intensity (See Methods). Torch condensates showed partial recovery over 240 s (n = 10 independent FRAP measurements; mean ± SD), indicating internal molecular exchange. **(g-i)** Representative images (g) and quantification of condensate number per field (h) and condensate area fraction (i) for Torch-AF488 condensates following 1,6-hexanediol (1,6-HD) treatment. Torch-AF488 condensates assembled from 80 μM Torch-AF488 in 10% PEG 8000 for 0.5–1 h were treated with either 0% 1,6-HD (control) or 10% 1,6-HD for 20 min before imaging. 1,6-HD significantly reduced Torch condensate number and size, supporting the hydrophobic-interaction-dependent, phase-separated nature of Torch condensates; $n$ = 5 independent fields per condition. Statistical significances were assessed using two-sided Student's unpaired $t$-tests: (h) $t = 3.222$, df = 8; (i) $t = 5.291$, df = 8. Scale bars, 10 μm (b and g), 2 μm (e), and 5 μm (f). **(j)** Torch CUT&Tag peak distribution across genomic features in gynes and workers. Called peaks were annotated to genomic features, revealing widespread Torch occupancy with enrichment at promoter-proximal regions (−3 kb to +1 kb relative to transcription start sites). Gynes harbor 24,089 peaks with 32% of them promoter-proximal (5,071 genes), whereas workers harbor 57,119 peaks with ~20% being promoter-proximal (7,881 genes) (Supplementary Dataset 9). **(k)** De novo motif discovery identified GA-repeat consensus motifs as the most significantly enriched motifs within Torch CUT&Tag peaks: CTCTCYCTCTCTCTC (reverse complement GAGAGAGAGRGAGAG; *E*-value = 4.2E-281) in gyne broods and TCTCTCYCYCTCTCT (reverse complement AGAGAGRGRGAGAGA; *E*-value = 7.5E-277) in worker broods. Both motifs were significantly enriched at peak centers (CentriMo *E*-value = 1.5E-8 and 1.3E-22, respectively) and most closely matched the *D. melanogaster* Trithorax-like motif (Tomtom *E*-value = 0.0109 and 0.0141, respectively; Supplementary Dataset 9). **(l-n)** ITC (l), SPR (m), and MST (n) assays demonstrating direct binding of purified *M. pharaonis* Torch to the GA-dinucleotide–rich DNA oligonucleotide GAGAGAGAGAGAGAG (5'–3'), with dissociation constants $K_D$ of 0.508 μM (ITC), 1.29 μM

(SPR), and 0.52 ± 0.05 μM (MST). **(o)** SPR assay showing no detectable binding of Torch to the scrambled control DNA GGAAGGAAGGAAGGA (5'–3'). **(p)** SPR assay demonstrating direct binding of Torch to the GAGAG pentanucleotide with dissociation constant $K_D$ of 81.2 μM. RU, resonance units (m, o, p).

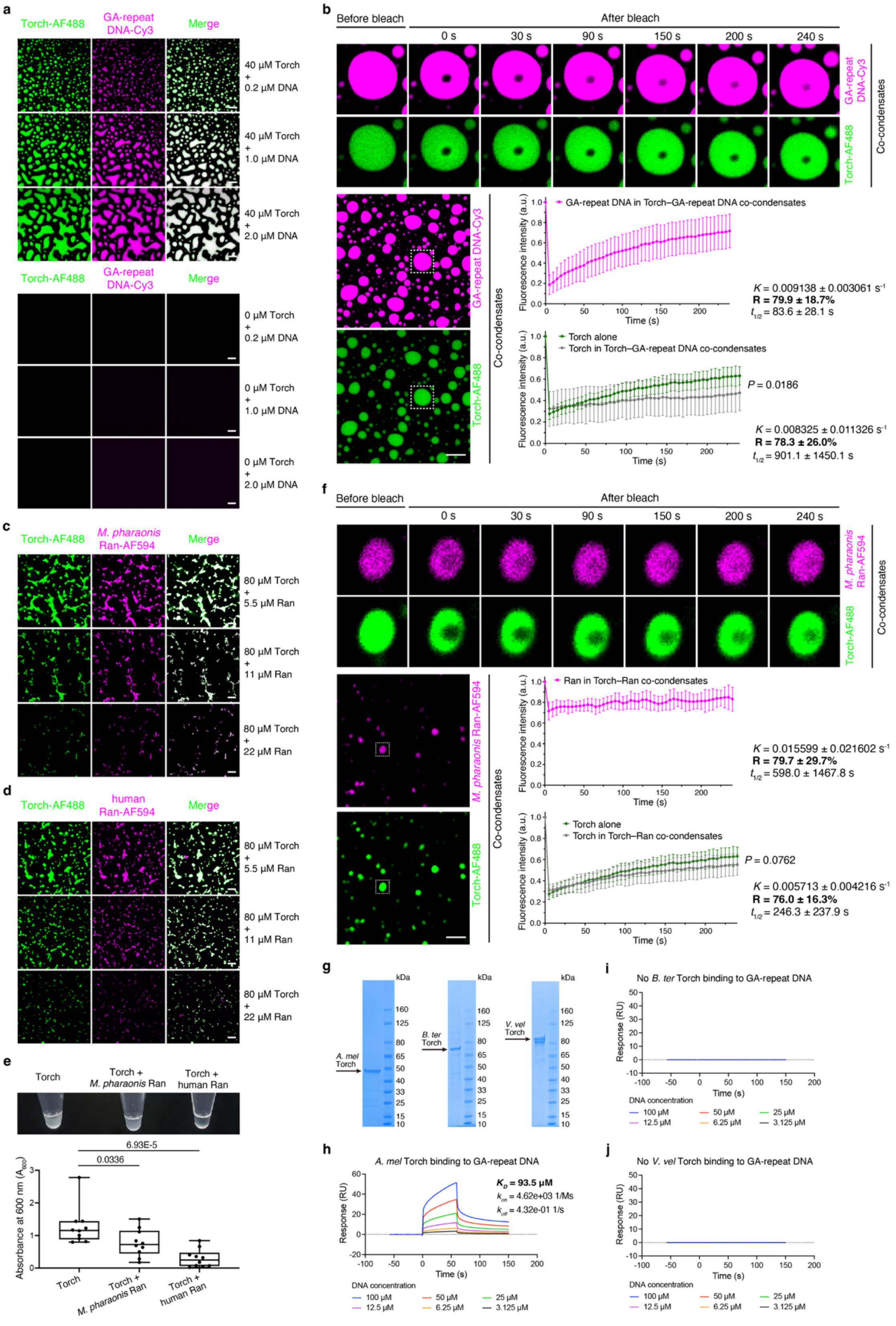
a
Torch-AF488
GA-repeat DNA-Cy3
Merge
40 μM Torch + 0.2 μM DNA
40 μM Torch + 1.0 μM DNA
40 μM Torch + 2.0 μM DNA
0 μM Torch + 0.2 μM DNA
0 μM Torch + 1.0 μM DNA
0 μM Torch + 2.0 μM DNA
b
Before bleach
After bleach
0 s
30 s
90 s
150 s
200 s
240 s
Co-condensates
GA-repeat DNA in Torch–GA-repeat DNA co-condensates
Fluorescence intensity (a.u.)
Time (s)
K = 0.009138 ± 0.003061 s-1
R = 79.9 ± 18.7%
t1/2 = 83.6 ± 28.1 s
Torch alone
Torch in Torch–GA-repeat DNA co-condensates
P = 0.0186
K = 0.008325 ± 0.011326 s-1
R = 78.3 ± 26.0%
t1/2 = 901.1 ± 1450.1 s
c
M. pharaonis Ran-AF594
80 μM Torch + 5.5 μM Ran
80 μM Torch + 11 μM Ran
80 μM Torch + 22 μM Ran
d
human Ran-AF594
e
Torch
Torch + M. pharaonis Ran
Torch + human Ran
6.93E-5
0.0336
Absorbance at 600 nm (A600)
f
Ran in Torch–Ran co-condensates
K = 0.015599 ± 0.021602 s-1
R = 79.7 ± 29.7%
t1/2 = 598.0 ± 1467.8 s
Torch in Torch–Ran co-condensates
P = 0.0762
K = 0.005713 ± 0.004216 s-1
R = 76.0 ± 16.3%
t1/2 = 246.3 ± 237.9 s
g
kDa
A. mel Torch
B. ter Torch
V. vel Torch
h
A. mel Torch binding to GA-repeat DNA
KD = 93.5 μM
kon = 4.62e+03 1/Ms
koff = 4.32e-01 1/s
Response (RU)
DNA concentration
100 μM
50 μM
25 μM
12.5 μM
6.25 μM
3.125 μM
i
No B. ter Torch binding to GA-repeat DNA
j
No V. vel Torch binding to GA-repeat DNA

**Extended Data Fig. 7. Differential tuning of Torch condensates by GA-repeat DNA and Ran and enhanced GA-repeat recognition by the ant Torch ortholog. (a)** Representative confocal fluorescence images of Alexa Fluor 488–labeled Torch (Torch-AF488; green) mixed with Cy3-labeled GA-repeat DNA (GA-repeat DNA-Cy3; magenta). In the presence of 40 μM Torch, GA-repeat DNA partitioned into Torch condensates and promoted the formation of enlarged Torch–GA-repeat DNA co-condensates in a DNA concentration-dependent manner. In contrast, GA-repeat DNA alone, tested at the same concentrations in the absence of Torch, did not form detectable condensates, indicating that GA-repeat DNA enhances Torch condensation but insufficiently to induce phase separation by itself. **(b)** Dual-color FRAP analysis of Torch–GA-repeat DNA co-condensates. Representative time-lapse images show selective photobleaching and recovery of GA-repeat DNA-Cy3 and Torch-AF488 within the same co-condensate. GA-repeat DNA recovered after photobleaching within Torch-containing condensates, indicating that GA-repeat DNA dynamically participates in the molecular exchange of Torch condensates. Torch also recovered within Torch–GA-repeat DNA co-condensates but showed reduced endpoint recovery compared with Torch-only condensates, indicating that GA-repeat DNA stabilizes Torch condensates and decreases Torch mobility. $n$ = 10 independent FRAP measurements per group; means ± SD. Endpoint recovery at 235 s was 0.631 ± 0.086 for Torch-only condensates and 0.472 ± 0.167 for Torch in Torch–GA-repeat DNA co-condensates (two-sided Welch's unpaired $t$-test, $t$ = 2.678, df = 13.41). One-phase association fitting gave $K$ = 0.009138 ± 0.003061 $s^{-1}$, recovery fraction $R$ = 79.9 ± 18.7%, and $t_{1/2}$ = 83.6 ± 28.1 s for GA-repeat DNA in co-condensates; for Torch in Torch–GA-repeat DNA co-condensates, $K$ = 0.008325 ± 0.011326 $s^{-1}$, $R$ = 78.3 ± 26.0%, and $t_{1/2}$ = 901.1 ± 1450.1 s. **(c–d)** Representative confocal fluorescence images of Torch-AF488 mixed with increasing concentrations of Alexa Fluor 594–labeled *M. pharaonis* Ran (c) or human Ran (d). Both *M. pharaonis* Ran and human Ran co-partitioned with Torch and progressively reduced the apparent size of Torch condensates, indicating that Ran constrains Torch condensate formation. **(e)** Turbidity assay measuring Torch condensation in the absence or presence of *M. pharaonis* Ran or human Ran. Representative tubes are shown above the quantification. Lower absorbance at 600 nm ($A_{600}$) indicates a clearer solution and reduced macroscopic Torch condensation. Addition of Ran decreased turbidity relative to Torch alone, with human Ran producing the strongest reduction, consistent with the suppression of Torch condensate formation observed by fluorescence microscopy; $n$ = 10 measurements per group. Statistical significance was assessed using one-way ANOVA followed by Dunnett's multiple comparisons tests for each group with the Torch-only control: $F(2, 27)$ = 12.126, $P$ = 1.75E-4. Dunnett-adjusted $P$ values are indicated in the figure. **(f)** Dual-color FRAP analysis of Torch–*M. pharaonis* Ran co-condensates. Selective photobleaching showed recovery of *M. pharaonis* Ran-AF594 within Torch–Ran co-condensates, indicating that Ran is dynamically incorporated into Torch condensates and co-exchanged with Torch. Torch-AF488 was also recovered after photobleaching, but Torch endpoint recovery was not significantly different between Torch-only condensates and Torch–*M. pharaonis* Ran co-condensates (0.631 ± 0.086 versus 0.553 ± 0.100 at 235 s). Thus, *M. pharaonis* Ran appears to reduce condensate scale without significantly altering internal Torch mobility. One-phase association fitting gave $K$ = 0.015599 ± 0.021602 $s^{-1}$, $R$ = 79.7 ± 29.7%, and $t_{1/2}$ = 598.0 ± 1467.8 s for *M. pharaonis* Ran in co-condensates; for Torch in Torch–*M. pharaonis* Ran co-condensates, we obtained $K$ = 0.005713 ± 0.004216 $s^{-1}$, $R$ = 76.0 ± 16.3%, and $t_{1/2}$ = 246.3 ± 237.9 s. Confocal LLPS samples in (a, c, and d) were incubated at 4°C overnight with 10% PEG 8000, as described for Extended Data Fig. 6b.

FRAP procedures were performed as described for Extended Data Fig. 6f, using 40 μM Torch plus 2 μM GA-repeat DNA in (b) and 80 μM Torch plus 22 μM *M. pharaonis* Ran in (f). FRAP curves are shown as means ± SD; endpoint recovery at 235 s was compared with Torch-only condensates by two-sided Welch's unpaired *t*-test; $t = 1.884$, $df = 17.56$; $n = 10$ independent FRAP measurements per group. Scale bars, 10 μm (a, c, d) and 5 μm (b, f). **(g–j)** Reduced GA-repeat binding by non-ant hymenopteran Torch orthologs. **(g)** SDS–PAGE validation of purified recombinant His-tagged Torch proteins used for the SPR assays. Predicted molecular masses are 42.6 kDa for *Apis mellifera* Torch, 66.7 kDa for *Bombus terrestris* Torch, and 89.4 kDa for *Vespa velutina* Torch; highlighted bands indicate the expected products. The same His–Torch proteins were used for His-pull-down assays (Supplementary Dataset 10). **(h-j)** SPR assays measuring binding of Torch orthologs from the honeybee *Apis mellifera* (h), bumble bee *Bombus terrestris* (i), and hornet *Vespa velutina* (j) to a GA-dinucleotide–rich DNA oligonucleotide, GAGAGAGAGAGAGAG, oriented 5' to 3'. Under these assay conditions, *B. terrestris* and *V. velutina* Torch showed no detectable binding, whereas *A. mellifera* Torch bound weakly, with an equilibrium dissociation constant of $K_D = 93.5$ μM. This affinity is approximately two orders of magnitude lower than that of *M. pharaonis* Torch shown in Extended Data Fig. 6l-n, supporting enhanced GA-repeat recognition by the ant Torch ortholog relative to the non-ant hymenopteran orthologs tested; RU, resonance units.

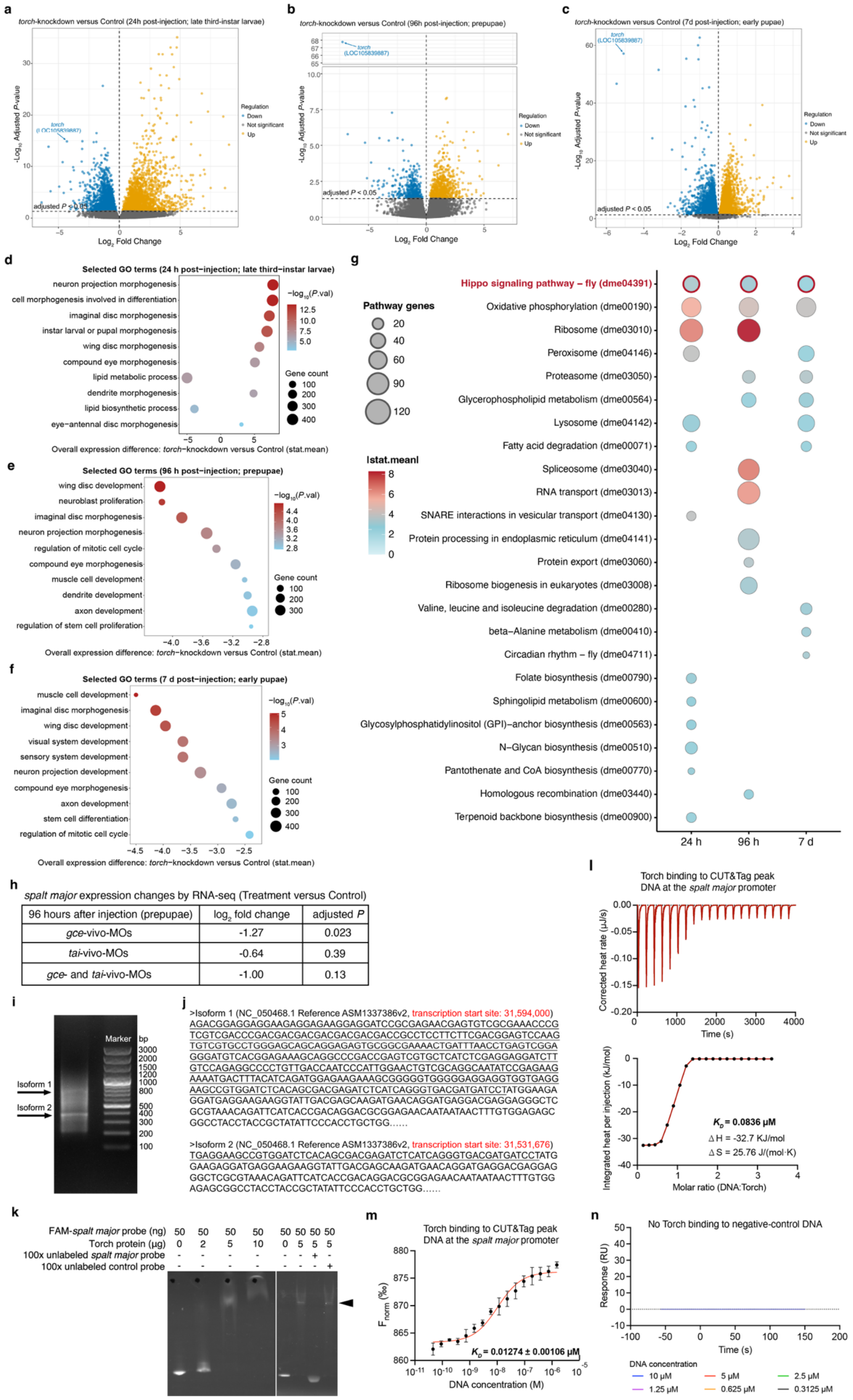

a
torch-knockdown versus Control (24h post-injection; late third-instar larvae)
b
torch-knockdown versus Control (96h post-injection; prepupae)
c
torch-knockdown versus Control (7d post-injection; early pupae)
torch (LOC105839887)
Regulation
Down
Not significant
Up
adjusted P < 0.05
-Log10 Adjusted P-value
Log2 Fold Change
d
Selected GO terms (24 h post−injection; late third−instar larvae)
neuron projection morphogenesis
cell morphogenesis involved in differentiation
imaginal disc morphogenesis
instar larval or pupal morphogenesis
wing disc morphogenesis
compound eye morphogenesis
lipid metabolic process
dendrite morphogenesis
lipid biosynthetic process
eye−antennal disc morphogenesis
−log10(P.val)
Gene count
Overall expression difference: torch−knockdown versus Control (stat.mean)
e
Selected GO terms (96 h post−injection; prepupae)
wing disc development
neuroblast proliferation
imaginal disc morphogenesis
neuron projection morphogenesis
regulation of mitotic cell cycle
compound eye morphogenesis
muscle cell development
dendrite development
axon development
regulation of stem cell proliferation
Overall expression difference: torch−knockdown versus Control (stat.mean)
f
Selected GO terms (7 d post−injection; early pupae)
muscle cell development
imaginal disc morphogenesis
wing disc development
visual system development
sensory system development
neuron projection development
compound eye morphogenesis
axon development
stem cell differentiation
regulation of mitotic cell cycle
Overall expression difference: torch−knockdown versus Control (stat.mean)
g
Pathway genes
|stat.mean|
Hippo signaling pathway – fly (dme04391)
Oxidative phosphorylation (dme00190)
Ribosome (dme03010)
Peroxisome (dme04146)
Proteasome (dme03050)
Glycerophospholipid metabolism (dme00564)
Lysosome (dme04142)
Fatty acid degradation (dme00071)
Spliceosome (dme03040)
RNA transport (dme03013)
SNARE interactions in vesicular transport (dme04130)
Protein processing in endoplasmic reticulum (dme04141)
Protein export (dme03060)
Ribosome biogenesis in eukaryotes (dme03008)
Valine, leucine and isoleucine degradation (dme00280)
beta−Alanine metabolism (dme00410)
Circadian rhythm – fly (dme04711)
Folate biosynthesis (dme00790)
Sphingolipid metabolism (dme00600)
Glycosylphosphatidylinositol (GPI)−anchor biosynthesis (dme00563)
N−Glycan biosynthesis (dme00510)
Pantothenate and CoA biosynthesis (dme00770)
Homologous recombination (dme03440)
Terpenoid backbone biosynthesis (dme00900)
24 h
96 h
7 d
h
spalt major expression changes by RNA-seq (Treatment versus Control)
96 hours after injection (prepupae) | log2 fold change | adjusted P
gce-vivo-MOs | -1.27 | 0.023
tai-vivo-MOs | -0.64 | 0.39
gce- and tai-vivo-MOs | -1.00 | 0.13
i
Marker
bp
3000
2000
1500
1200
1000
800
500
400
300
200
100
Isoform 1
Isoform 2
j
>Isoform 1 (NC_050468.1 Reference ASM1337386v2, transcription start site: 31,594,000)
AGACGGAGGAGGAAGAGGAGAAGGAGGATCCGCGAGAACGAGTGTCGCGAAACCCG
TCGTCGACCCGACGACGACGACGACGACGACCGCCTCCTTCTTCGACGGAGTCCAAG
TGTCGTGCCTGGGAGCAGCAGGAGAGTGCGGCGAAAACTGATTTAACCTGAGTCGGA
GGGATGTCACGGAGAAAGCAGGCCCGACCGAGTCGTGCTCATCTCGAGGAGGATCTT
GTCCAGAGGCCCCTGTTGACCAATCCCATTGGAACTGTCGCAGGCAATATCCGAGAAG
AAAATGACTTTACATCAGATGGAGAAGAAAGCGGGGGTGGGGGAGGAGGTGGTGAGG
AAGCCGTGGATCTCACAGCGACGAGATCTCATCAGGGTGACGATGATCCTATGGAAGA
GGATGAGGAAGAAGGTATTGACGAGCAAGATGAACAGGATGAGGACGAGGAGGGCTC
GCGTAAACAGATTCATCACCGACAGGACGCGGAGAACAATAATAACTTTGTGGAGAGC
GGCCTACCTACCGCTATATTCCCACCTGCTGG......
>Isoform 2 (NC_050468.1 Reference ASM1337386v2, transcription start site: 31,531,676)
TGAGGAAGCCGTGGATCTCACAGCGACGAGATCTCATCAGGGTGACGATGATCCTATG
GAAGAGGATGAGGAAGAAGGTATTGACGAGCAAGATGAACAGGATGAGGACGAGGAG
GGCTCGCGTAAACAGATTCATCACCGACAGGACGCGGAGAACAATAATAACTTTGTGG
AGAGCGGCCTACCTACCGCTATATTCCCACCTGCTGG......
k
FAM-spalt major probe (ng) 50 50 50 50 50 50 50 50
Torch protein (µg) 0 2 5 10 0 5 5 5
100x unlabeled spalt major probe - - - - - - + -
100x unlabeled control probe - - - - - - - +
l
Torch binding to CUT&Tag peak DNA at the spalt major promoter
Corrected heat rate (µJ/s)
Time (s)
Integrated heat per injection (kJ/mol)
Molar ratio (DNA:Torch)
KD = 0.0836 µM
ΔH = -32.7 KJ/mol
ΔS = 25.76 J/(mol·K)
m
Torch binding to CUT&Tag peak DNA at the spalt major promoter
Fnorm (‰)
KD = 0.01274 ± 0.00106 µM
DNA concentration (M)
n
No Torch binding to negative-control DNA
Response (RU)
Time (s)
DNA concentration
10 µM
5 µM
2.5 µM
1.25 µM
0.625 µM
0.3125 µM

**Extended Data Fig. 8. Stage-resolved analyses identify Hippo-associated growth programs and direct activation of *spalt major* by Torch. (a-g)** Differential expression and GO enrichment after *torch* knockdown. **(a-c)** Volcano plots of single-individual RNA-seq comparing *torch*-knockdown gynes with controls at 24 h post-injection (late third-instar larvae) (a), 96 h post-injection (prepupae) (b) and 7 d post-injection (early pupae) (c). Points are colored by regulation status (orange, up; blue, down; gray, no significant deviation). The target gene *torch* (LOC105839887) was annotated and strongly downregulated at each time points, as expected. The $\log_2$ fold-changes (adjusted *P* values) for *torch* were -4.30 (1.50E-15), -7.25 (1.71E-68) and -5.08 (7.43E-58) at 24 h, 96h and 7 d post-injection, respectively (Supplementary Dataset 11). **(d-f)** Ten selected GO term enrichments at 24 h (d), 96 h (e), and 7 d (f) post-injection, corresponding to the same time points as in (a–c). Bubble plots display the overall expression shift (*torch*-knockdown vs control; x-axis), term significance (color, $-\log_{10} p$-value) and gene count (bubble size). Significantly enriched terms (*P* value < 0.01) were related to imaginal-disc, eye, neuron, muscle development, lipid metabolism and neuroblast proliferation. These transcriptomic signatures mirror the observed phenotypes in imaginal-disc–derived organs, fat body and brain. **(g)** Recurrent enrichment of Hippo signaling across three developmental time points. The figure shows all KEGG pathways significantly enriched between *torch*-knockdown and control gynes at 24 h (late third-instar larvae), 96 h (prepupae) and 7 d (early pupae) ($P < 0.01$). The KEGG term dme04391 Hippo signaling pathway – fly was recurrently enriched across all three developmental stages (Supplementary Dataset 12). **(h)** RNA-seq–based expression changes of *spalt major* following loss of Gce, Tai, or Gce+Tai function based on values extracted from Supplementary Dataset 6. Expression of *spalt major* was significantly reduced upon Gce loss (adjusted $P < 0.05$) and showed a weaker downward expression trends upon Tai and Gce+Tai perturbation, consistent with Tai's pleiotropic co-activator role in multiple signaling pathways [26-29] that could partially buffer its impact on individual Torch targets. **(i-m)** *spalt major* 5'-RACE mapping and Torch-promoter binding assays. **(i)** 5'-RACE products from gyne broods revealed two *spalt major* 5'-UTR isoforms. **(j)** 5'-RACE product sequences with *spalt major* TSSs mapped to chromosome 2 (NC_050468.1): 31,594,000 and 31,531,676. Underlined sequences indicate the 5' UTRs, and the raw Sanger sequencing data are provided in Supplementary Dataset 7. **(k)** EMSA using FAM (carboxy-fluorescein)-labeled DNA probes spanning the Torch CUT&Tag peak in the *spalt major* promoter (chr2:31,595,843–31,595,962). Recombinant Torch induces dose-dependent mobility shifts, yielding progressively slower-migrating DNA–protein complexes at increasing protein concentrations; the shift is abolished by a 100-fold molar excess of unlabeled cognate DNA, but not by a 100-fold molar excess of unlabeled negative-control competitor DNA fragment (chr2:17,852,893–17,853,012; arrowhead), supporting sequence-specific binding. **(l)** ITC for recombinant Torch binding to the Torch CUT&Tag peak DNA fragment in the *spalt major* promoter (chr2:31,595,843–31,595,962), yielding $K_D = 0.0836$ µM. **(m)** MST for the same fragment, corroborating high-affinity binding ($K_D = 0.01274 \pm 0.00106$ µM). **(n)** SPR detected no binding of Torch to a 120-bp negative-control DNA fragment selected from our CUT&Tag dataset from a genomic region lacking Torch occupancy and GA-repeat enrichment; the fragment length matched that of the *spalt major* promoter CUT&Tag peak DNA fragment. Genomic coordinates: chr2:17,852,893–17,853,012. RU, resonance units.

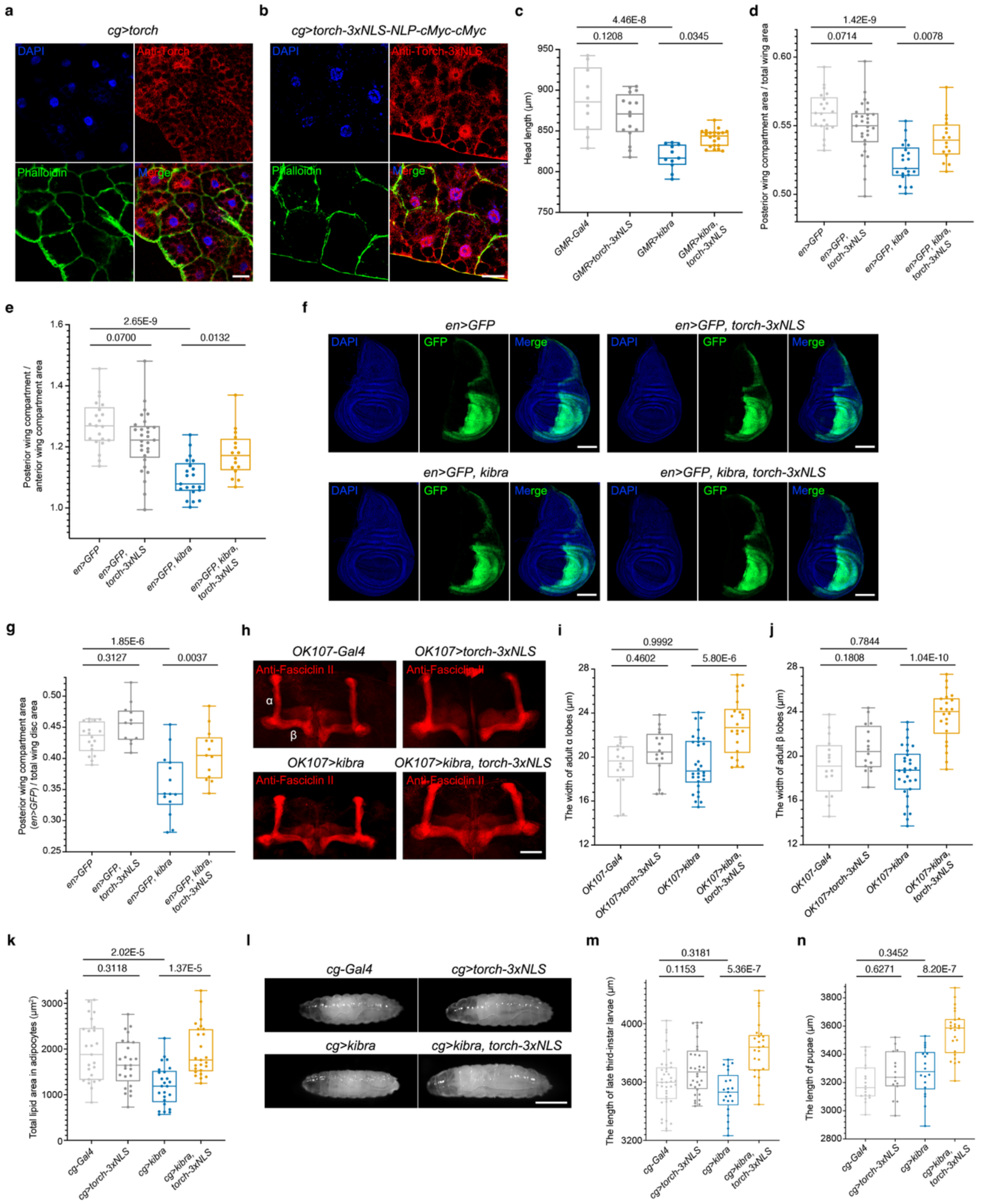

a
cg>torch
DAPI
Anti-Torch
Phalloidin
Merge
b
cg>torch-3xNLS-NLP-cMyc-cMyc
DAPI
Anti-Torch-3xNLS
Phalloidin
Merge
c
4.46E-8
0.1208
0.0345
Head length (μm)
GMR-Gal4
GMR>torch-3xNLS
GMR>kibra
GMR>kibra, torch-3xNLS
d
1.42E-9
0.0714
0.0078
Posterior wing compartment area / total wing area
en>GFP
en>GFP, torch-3xNLS
en>GFP, kibra
en>GFP, kibra, torch-3xNLS
e
2.65E-9
0.0700
0.0132
Posterior wing compartment / anterior wing compartment area
f
en>GFP
en>GFP, torch-3xNLS
en>GFP, kibra
en>GFP, kibra, torch-3xNLS
DAPI
GFP
Merge
g
1.85E-6
0.3127
0.0037
Posterior wing compartment area (en>GFP) / total wing disc area
h
OK107-Gal4
OK107>torch-3xNLS
OK107>kibra
OK107>kibra, torch-3xNLS
Anti-Fasciclin II
α
β
i
0.9992
0.4602
5.80E-6
The width of adult α lobes (μm)
OK107-Gal4
OK107>torch-3xNLS
OK107>kibra
OK107>kibra, torch-3xNLS
j
0.7844
0.1808
1.04E-10
The width of adult β lobes (μm)
k
2.02E-5
0.3118
1.37E-5
Total lipid area in adipocytes (μm²)
cg-Gal4
cg>torch-3xNLS
cg>kibra
cg>kibra, torch-3xNLS
l
cg-Gal4
cg>torch-3xNLS
cg>kibra
cg>kibra, torch-3xNLS
m
0.3181
0.1153
5.36E-7
The length of late third-instar larvae (μm)
n
0.3452
0.6271
8.20E-7
The length of pupae (μm)

**Extended Data Fig. 9. Nuclear-targeted Torch promotes Hippo-dependent growth across multiple fly tissues. (a–b)** *D. melanogaster* third-instar larval fat body (*cg-GAL4*). Immunofluorescence for Torch (red) and Alexa Fluor™ 488–conjugated phalloidin (F-actin; green) stained with DAPI (blue). Torch without an exogenous NLS is largely non-nuclear (a), whereas Torch-3×NLS is prominently nuclear (b). **(c)** Head-length quantification with eye-specific expression (*GMR-GAL4*); Torch-3×NLS partially rescues Kibra-induced head-size reduction. **(d–g)** Posterior wing compartment (*en-GAL4*). Quantification of the posterior compartment-to-total wing area ratio (d) and posterior compartment-to-anterior compartment area ratio (e) in adult wings, related to Fig. 5c–d. Wing imaginal discs from late non-wandering third-instar larvae are shown in f, with GFP marking the *en-GAL4* expression domain (green) and DAPI marking nuclei (blue). Quantification of relative posterior compartment area is shown in g. Torch-3×NLS expanded the posterior compartment only under Kibra-mediated Hippo activation. **(h–j)** Mushroom body (*OK107-Gal4*). Adult brains stained with anti-Fasciclin II (red) [30] (h) with α- and β-lobes highlighted. Quantification of α-lobe (i) and β-lobe (j) widths showed increased lobe size upon co-expression of Kibra and Torch-3×NLS. **(k)** Fat body (*cg-Gal4*) with total lipid area per adipocyte showing partial rescue after co-expression of Kibra and Torch-3×NLS. **(l–m)** Late third-instar body size (*cg-Gal4*) for representative late third-instar larvae (l), with corresponding measurements showing increased body-length upon Torch-3×NLS expression in the Kibra (Hippo-activated) background (m). **(n)** Pupal body length showing a similar trend. Statistics based on one-way ANOVA followed by Šídák-adjusted planned pairwise comparisons. Šídák-adjusted $P$ values are indicated in the figure. Each data point represents an individual animal, except in panel k, where each point represents an individual adipocyte. Sample sizes follow the left-to-right genotype order in each quantification panel: (c) $n$ = 10, 16, 11 and 21; $F(3, 54) = 18.43$, $P = 2.33\text{E-}8$; (d) $n$ = 21, 29, 21 and 16; $F(3, 83) = 17.91$, $P = 4.72\text{E-}9$; (e) $n$ = 21, 29, 21 and 16; $F(3, 83) = 17.21$, $P = 8.90\text{E-}9$; (g) $n$ = 18, 12, 14 and 13; $F(3, 53) = 17.19$, $P = 6.34\text{E-}8$; (i) $n$ = 16, 16, 30 and 22; $F(3, 80) = 10.50$, $P = 6.73\text{E-}6$; (j) $n$ = 16, 16, 30 and 22; $F(3, 80) = 21.49$, $P = 2.64\text{E-}10$; (k) $n$ = 25 per group; $F(3, 96) = 10.30$, $P = 6.13\text{E-}6$; (m) $n$ = 39, 32, 21 and 25; $F(3, 113) = 12.33$, $P = 4.88\text{E-}7$; (n) $n$ = 18, 15, 18 and 25; $F(3, 72) = 22.55$, $P = 2.12\text{E-}10$. Scale bars, 30 µm (a, b), 100 µm (f), 50 µm (h) and 1 mm (l).

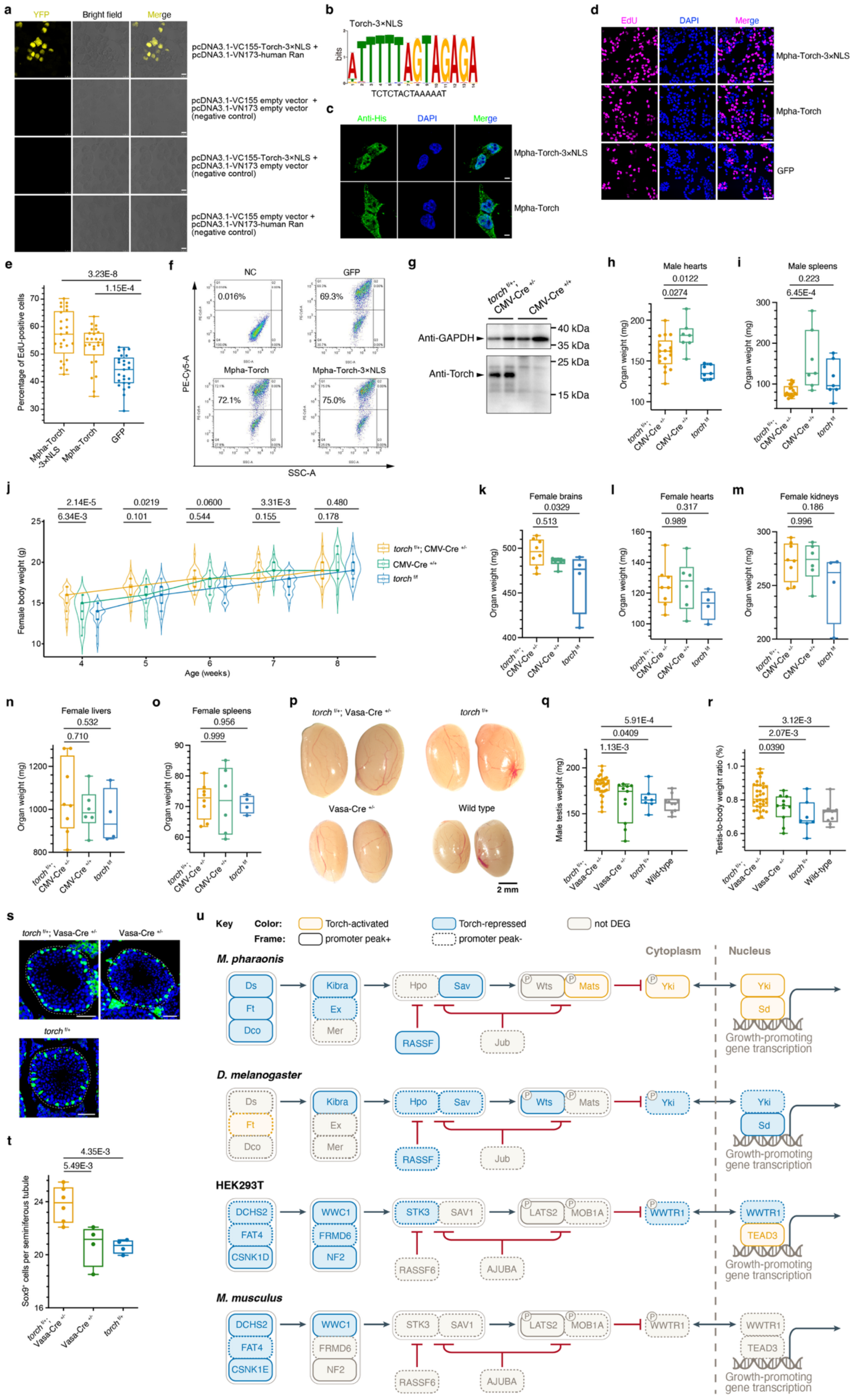
a
YFP
Bright field
Merge
pcDNA3.1-VC155-Torch-3×NLS + pcDNA3.1-VN173-human Ran
pcDNA3.1-VC155 empty vector + pcDNA3.1-VN173 empty vector (negative control)
pcDNA3.1-VC155-Torch-3×NLS + pcDNA3.1-VN173 empty vector (negative control)
pcDNA3.1-VC155 empty vector + pcDNA3.1-VN173-human Ran (negative control)
b
Torch-3×NLS
bits
TCTCTACTAAAAAT
c
Anti-His
DAPI
Merge
Mpha-Torch-3×NLS
Mpha-Torch
d
EdU
DAPI
Merge
Mpha-Torch-3×NLS
Mpha-Torch
GFP
e
Percentage of EdU-positive cells
3.23E-8
1.15E-4
Mpha-Torch -3×NLS
Mpha-Torch
GFP
f
NC
GFP
0.016%
69.3%
Mpha-Torch
Mpha-Torch-3×NLS
72.1%
75.0%
PE-Cy5-A
SSC-A
g
torch f/+; CMV-Cre +/-
CMV-Cre +/+
Anti-GAPDH
Anti-Torch
40 kDa
35 kDa
25 kDa
15 kDa
h
Male hearts
0.0122
0.0274
Organ weight (mg)
i
Male spleens
0.223
6.45E-4
Organ weight (mg)
j
Female body weight (g)
Age (weeks)
2.14E-5
6.34E-3
0.0219
0.101
0.0600
0.544
3.31E-3
0.155
0.480
0.178
torch f/+; CMV-Cre +/-
CMV-Cre +/+
torch f/f
k
Female brains
0.0329
0.513
l
Female hearts
0.317
0.989
m
Female kidneys
0.186
0.996
n
Female livers
0.532
0.710
o
Female spleens
0.956
0.999
p
torch f/+; Vasa-Cre +/-
torch f/+
Vasa-Cre +/-
Wild type
2 mm
q
Male testis weight (mg)
5.91E-4
0.0409
1.13E-3
r
Testis-to-body weight ratio (%)
3.12E-3
2.07E-3
0.0390
Wild-type
s
torch f/+; Vasa-Cre +/-
Vasa-Cre +/-
torch f/+
t
Sox9+ cells per seminiferous tubule
4.35E-3
5.49E-3
u
Key
Color:
Torch-activated
Torch-repressed
not DEG
Frame:
promoter peak+
promoter peak-
Cytoplasm
Nucleus
M. pharaonis
D. melanogaster
HEK293T
M. musculus
Ds
Ft
Dco
Kibra
Ex
Mer
Hpo
Sav
Wts
Mats
Yki
Sd
RASSF
Jub
DCHS2
FAT4
CSNK1D
CSNK1E
WWC1
FRMD6
NF2
STK3
SAV1
LATS2
MOB1A
WWTR1
TEAD3
RASSF6
AJUBA
Growth-promoting gene transcription

**Extended Data Fig. 10. Torch promotes mammalian cell proliferation and organ growth. (a-f)** Torch retains nuclear regulatory properties and promotes S-phase entry in mammalian cells. **(a)** Bimolecular fluorescence complementation assay showing a direct nuclear Torch–human Ran interaction in HEK293T cells. Torch–3×NLS was fused to the C-terminal YFP Venus fragment VC155 (residues 155–238), and human Ran was fused to the N-terminal YFP Venus fragment VN173 (residues 1–173). Nuclear YFP fluorescence indicated complementation upon Torch–Ran interaction [31]. Representative YFP, bright-field and merged images are shown for pcDNA3.1 empty-vector controls in combination with pcDNA3.1-based expression constructs. Scale bars, 10 µm. **(b)** De novo motif discovery in Torch–3×NLS-bound peaks from HEK293T cells, showing TCTCTACTAAAAAT, whose reverse complement is ATTTTTAGTAGAGA, as the second-ranked most enriched consensus motif by *E*-value ($E$ = 3.8E-817; Supplementary Dataset 13). The motif was strongly centrally enriched within Torch-bound peaks ($E$ = 8.1E-820) and contains the pentanucleotide AGAGA, consistent with Torch's GA-repeat binding preference in ant chromatin. **(c)** Stable HEK293T cell lines carrying genome-integrated constructs encoding His-tagged *M. pharaonis* Torch, either unmodified or fused to a 3×NLS, were analyzed by immunofluorescence. His-tag staining (green) with DAPI nuclear counterstaining (blue) revealed mixed nucleocytoplasmic localization of unmodified Torch, whereas Torch-3×NLS accumulated predominantly in the nucleus; scale bars: 5 µm. **(d–e)** Imaging-based analysis of EdU incorporation in genome-integrated, UbC promoter–driven HEK293T cell lines expressing GFP, *M. pharaonis* Torch, either without an exogenous NLS or fused to 3×NLS. **(d)** Representative confocal images showing EdU incorporation (magenta) and DAPI nuclear counterstaining (blue); scale bars, 50 µm. **(e)** Quantification of the percentage of EdU-positive cells from panel d. Each data point represents the percentage of EdU-positive cells quantified from one confocal field derived from one biological replicate ($n$ = 25 confocal fields per group). Statistical significances were assessed using one-way ANOVA followed by Dunnett's multiple-comparisons test against the GFP control ($F(2, 72) = 21.03$, $P$ = 6.40E-8). **(f)** Representative flow cytometry plots showing EdU incorporation in HEK293T cells, corresponding to the quantification in Fig. 6b. The percentage of EdU-positive cells is indicated in each plot. NC indicates the no-EdU negative control used to set the EdU-positive gate, which was then applied uniformly to all samples. Mpha, *M. pharaonis*. **(g-t)** Sex- and organ-specific growth outputs of heterologous Torch expression in mice. **(g)** Western blot analysis of Torch expression in male liver tissue lysates from *torch* $^{f/+}$; CMV-Cre $^{+/-}$ mice and CMV-Cre $^{+/+}$ controls. Two biological replicates are shown for each genotype. Anti-Torch detected Torch in *torch* $^{f/+}$;CMV-Cre $^{+/-}$ samples but not in CMV-Cre $^{+/+}$ controls, with GAPDH used as a loading control. **(h–i)** Heart (h) and spleen (i) weights in two-month-old male mice; neither organ showed a consistent increase in CMV-Cre;R26-LSL-*torch* mice relative to both CMV-Cre-only and R26-LSL-*torch*-only controls. Each dot represents the mass of the indicated organ from an individual mouse. Data were analyzed separately for each organ by one-way ANOVA followed by Dunnett's multiple-comparisons tests: heart, $F(2, 28) = 10.99$, $P$ = 3.00E-4; spleen, $F(2, 28) = 8.471$, $P$ = 1.33E-3. For both panels, sample sizes in the left-to-right order shown are $n$ = 17, 7 and 7. **(j)** Body weights of female CMV-Cre;R26-LSL-*torch* mice from 4 to 8 weeks of age compared with control genotypes, showing no increase in body weight throughout development apart from a transient elevation at one month of age. Each point represents one mouse measured at the indicated age; lines connect group medians. Statistics: one-way ANOVA followed by Dunnett's multiple comparisons tests (adjustment applied separately within each age): 4 weeks, $F(2, 65) = 12.10$, $P$ = 3.42E-5; 5

weeks, $F(2, 65) = 3.86$, $P = 0.0260$; 6 weeks, $F(2, 65) = 2.40$, $P = 0.0984$; 7 weeks, $F(2, 67) = 11.07$, $P = 7.04E\text{-}5$; and 8 weeks, $F(2, 65) = 1.53$, $P = 0.223$. Sample sizes for CMV-Cre;R26-LSL-*torch*, CMV-Cre-only and R26-LSL-*torch*-only mice at 4–8 weeks were 28/17/23, 28/17/23, 28/17/23, 29/17/24 and 32/17/19, respectively. **(k–o)** Female organ weights at two months of age, showing no consistent increase in brain (k), heart (l), kidney (m), liver (n), or spleen (o) mass in CMV-Cre;R26-LSL-*torch* mice relative to both control genotypes. Each dot represents the mass of the indicated organ from an individual mouse. Data were analyzed for each organ by one-way ANOVA followed by Dunnett's multiple-comparisons tests: brain, $F(2, 15) = 3.568$, $P = 0.0540$; heart, $F(2, 15) = 1.212$, $P = 0.325$; kidney, $F(2, 15) = 1.688$, $P = 0.218$; liver, $F(2, 15) = 0.5672$, $P = 0.579$; spleen, $F(2, 15) = 0.03720$, $P = 0.964$. For all panels, sample sizes in the left-to-right order shown are $n$ = 8, 6 and 4. **(p–r)** Germline-specific activation of *torch* driven by Vasa-Cre. Representative images of testes (p), absolute testis mass (q) and testis-to-body-weight ratio (r) in two-month-old Vasa-Cre;R26-LSL-*torch* males versus controls, demonstrating a local gonadal growth effect. Each dot in q and r represents an individual mouse. Data were analyzed by one-way ANOVA followed by Dunnett's multiple-comparisons tests comparing Vasa-Cre;R26-LSL-*torch* mice with each of the three control genotypes. (q) $F(3, 57) = 8.649$, $P = 8.04E\text{-}5$; (r) $F(3, 56) = 7.508$, $P = 2.61E\text{-}4$. Sample sizes are listed in the left-to-right order shown: (q) $n$ = 33, 11, 7 and 10; (r) $n$ = 33, 11, 7 and 9. **(s–t)** Testes from two-month-old males stained for SOX9 to mark Sertoli cells (s) and quantification of $SOX9^{+}$ Sertoli cells per seminiferous tubule (t) in Vasa-Cre;R26-LSL-*torch* mice versus controls. Each dot in t represents an individual mouse. Data were analyzed by one-way ANOVA followed by Dunnett's multiple-comparisons tests: $F(2, 11) = 10.75$, $P = 2.58E\text{-}3$. Sample sizes in the left-to-right order shown are $n$ = 6, 4 and 4. See Fig. 6j-k for additional legend details. Scale bars in (s), 50 µm. Dunnett-adjusted $P$ values are shown in panels e, h–o, q–r and t. **(u)** Conserved direct targeting of Kibra/WWC1 by Torch across metazoan model systems. Hippo-pathway genes regulated by Torch were mapped onto the KEGG Hippo signaling pathway for multiple species (map04392). The *M. pharaonis* panel summarizes the stage-integrated knockdown analysis shown in Fig. 4c; the *D. melanogaster* panel compares *GMR>kibra, torch-3×NLS* female heads with *GMR>kibra* controls; the HEK293T panel combines Torch and Torch–3×NLS data; and the *M. musculus* panel combines brain and kidney data (Supplementary Datasets 13-14). Regulatory directions are shown relative to Torch activity: orange boxes indicate Torch-activated genes, blue boxes indicate Torch-repressed genes, and gray boxes indicate non-DEGs. Solid boxes denote direct targets, defined as DEGs with promoter-proximal Torch CUT&Tag peaks, whereas dashed boxes denote indirect targets. For mammalian genes with multiple paralogues, one representative paralogue showing an expression change consistent with the observed pro-growth effect of Torch was selected for visualization; complete paralog-specific changes are provided in Supplementary Dataset 14. Kibra/WWC1 is highlighted as a direct Torch-repressed Hippo regulator recurrently identified in ants, flies, HEK293T cells, and mice.

**Supplementary Dataset 1. Caste-biased expression and phylogenetic distribution of *torch* across ant species [9].** This dataset reports $\log_2$ fold changes and adjusted *P* values for *torch* expression in adult gynes/queens versus workers across 65 of the 68 ant species represented in the published GAGA transcriptomic dataset. Positive $\log_2$ fold-change values indicate higher expression in gynes/queens. It also provides *torch* orthologue assignments for 163 GAGA ant species, together with species names and subfamily classifications. No *torch* orthologue was detected in seven species. Because these species are distributed across four subfamilies and do not form a single ancestral clade, this pattern is consistent with multiple lineage-specific secondary losses of *torch*.

**Supplementary Dataset 2. Sequence, phylogenetic and evolutionary-rate data supporting positive-selection analyses of *torch*.** This dataset contains the orthologous *torch* coding sequences across Hymenoptera; the PRANK codon alignment generated directly from these coding sequences using the -codon -F parameters; phylogenetic tree files used for the branch-site analyses; and branch-specific *d*N/*d*S (ω) estimates and aBSREL *P* values, with nominal (uncorrected) $P < 0.01$ used as the threshold for evidence of positive selection irrespective of the branch-wide ω estimate.

**Supplementary Dataset 3. Reanalysis of caste- and juvenile hormone-associated developmental transcriptomes from [12].** This dataset contains the *t* statistics for the gyne-versus-JH-treated-worker and gyne-versus-control-worker comparisons, together with the corresponding $\Delta t$ values, for all genes at five developmental stages: early third instar, mid-third instar, late third instar, prepupae and early pupae. These *t* statistics were calculated as defined in our previous study [12]. At each developmental stage, $\Delta t$ was calculated as $t_{\text{gyne x JH-treated worker}} - t_{\text{gyne x control worker}}$. The x-axis value plotted in Fig. 1e represents the mean $\Delta t$ across the five stages, whereas the y axis shows $-\log_{10}$ adjusted *P* values for the comparison between JH-treated and control workers at the early pupal stage. Genes with a mean $\Delta t > 1$ or $< -1$ and an adjusted *P* value $< 0.05$ are highlighted in Fig. 1e. The dataset also contains differential gene expression analysis of gyne-versus-control-worker and JH-treated-worker-versus-control-worker comparisons at each developmental stage. For both contrasts, positive $\log_2$ fold-change values indicate higher expression in control workers.

**Supplementary Dataset 4. Protein sequence alignments and phylogenetic trees of *germ cell-expressed* (*gce*), *taiman* (*tai*), *PCNA* and *deadpan* orthologues.** This dataset contains protein sequence alignments and corresponding phylogenetic trees across representative ant species, the honeybee (*Apis mellifera*) and the fruit fly (*Drosophila melanogaster*).

**Supplementary Dataset 5. Sanger sequencing validation of vivo-MO-induced exon skipping in *gce* and *tai* transcripts.** This dataset contains Sanger sequencing results for PCR amplicons generated from late third-instar gyne larvae collected 24 h after injection with the splice-blocking *gce*-I3E4, *gce*-E6I6, *tai*-I2E3 or *tai*-I5E6 vivo-MO. For each treatment, raw chromatograms are provided for both the wild-type and vivo-MO-induced exon-skipped amplicons. These sequencing data verify the identities and splice junctions of the corresponding wild-type and exon-skipped *gce* and *tai* transcripts.

**Supplementary Dataset 6. Differential gene expression following splice-blocking vivo-MO targeting of *gce* and *tai* in gynes.** This dataset contains differential gene expression results from single-individual RNA-seq of gynes collected 24 h post-injection (late third-instar larvae) and 96 h post-injection (prepupae) following vivo-MO treatments targeting *gce*, *tai* or both genes, compared with control vivo-MO-injected gynes at the corresponding time point. For differential expression analysis, the *gce* group combined samples injected with *gce*-I3E4 or *gce*-E6I6, and the *tai* group combined samples injected with *tai*-I2E3 or *tai*-I5E6. At 24 h, the dual-target group combined samples co-injected with *gce*-I3E4 + *tai*-I5E6 or *gce*-E6I6 + *tai*-I2E3; at 96 h, it combined samples co-injected with *gce*-I3E4 + *tai*-I2E3 or *gce*-E6I6 + *tai*-I5E6. Negative $\log_2$ fold-change values indicate reduced expression in gynes with impaired Gce and/or Tai function relative to controls. Differentially expressed genes (DEGs; adjusted $P < 0.05$) are provided in a separate sheet.

**Supplementary Dataset 7. Sanger sequencing of 5'-RACE amplicons used to map the transcription start sites (TSSs) of *torch* and *spalt major*.** This dataset contains Sanger sequencing data for two 5'-RACE amplicons per gene, corresponding to two *torch* transcript isoforms identified in whole-body third-instar gyne larvae and two *spalt major* transcript isoforms identified in whole-body gyne prepupae. The *torch* TSSs map to chr11:16,123,782 and chr11:16,123,750 (NC_050477.1), whereas the *spalt major* TSSs map to chr2:31,594,000 and chr2:31,531,676 (NC_050468.1). Raw chromatograms are provided for each amplicon.

**Supplementary Dataset 8. Candidate Torch-interacting proteins identified by co-immunoprecipitation and GST–Torch pull-down mass spectrometry.** This dataset lists proteins detected specifically in anti-Torch immunoprecipitates, but absent from IgG negative controls, across three independent biological replicates, together with proteins detected specifically in one GST–Torch pull-down but absent from the negative-control pull-down, using *M. pharaonis* gyne-brood lysates and qualitative mass spectrometry. Torch (LOC105839887) was detected in all three anti-Torch IP protein lists and the GST–Torch pull-down list, confirming successful recovery of endogenous Torch in the IPs and the GST–Torch bait in the pull-down assay.

**Supplementary Dataset 9. Genome-wide Torch CUT&Tag peaks and motif analyses in gyne and worker broods.** This dataset contains Torch CUT&Tag peak coordinates and annotations from *M. pharaonis* gyne broods (24,089 peaks) and worker broods (57,119 peaks), with promoter-proximal peaks, defined as peaks located from −3 kb to +1 kb relative to a TSS, listed in separate sheets. MEME-ChIP de novo motif discovery, CentriMo central-enrichment and Tomtom motif-similarity results generated separately from the peak sequences of each caste are also provided. FIMO results are additionally provided for occurrences of the top gyne-brood motif, GAGAGAGAGRGAGAG, within gyne-brood Torch peaks in the *spalt major* promoter, retaining matches with $P < 0.01$. Notably, the prominent promoter-proximal peak on chromosome 2 (NC_050468.1; chr2:31,595,843–31,595,962) contained a highly significant match to this motif (FIMO $P < 0.0001$).

**Supplementary Dataset 10. Candidate proteins identified by His–Torch pull-down mass spectrometry in three non-ant hymenopterans.** This dataset lists proteins detected specifically in His–Torch pull-downs, but absent from the corresponding negative-control pull-downs, using worker-brood lysates from the honeybee *Apis mellifera*, bumble bee *Bombus terrestris* and hornet *Vespa velutina*. Separate qualitative mass-spectrometry protein lists are provided for each species. The corresponding Torch orthologues—*A. mellifera* LOC100577537, *B. terrestris* LOC100645125 and *V. velutina* LOC124954358—were detected in their respective His–Torch pull-down protein lists, confirming successful recovery of the His–Torch bait in each pull-down.

**Supplementary Dataset 11. Stage-resolved single-individual RNA-seq differential expression following vivo-MO-mediated *torch* knockdown in gynes.** This dataset contains differential gene expression results for all genes comparing *torch*-knockdown and control gynes at 24 h post-injection (late third-instar larvae), 96 h post-injection (prepupae) and 7 d post-injection (early pupae), with each RNA-seq library derived from an individual gyne. Negative $\log_2$ fold-change values indicate lower expression in *torch*-knockdown gynes. For each stage, DEGs (adjusted $P < 0.05$) are provided in a separate sheet.

**Supplementary Dataset 12. GAGE-based GO and KEGG enrichment following vivo-MO-mediated *torch* knockdown in gynes.** This dataset contains all Gene Ontology (GO) terms and KEGG pathways with GAGE-derived $P < 0.01$ at each developmental stage, based on the whole-transcriptome expression changes reported in Supplementary Dataset 11.

**Supplementary Dataset 13. Torch CUT&Tag profiles in heterologous expression systems.** This dataset contains genome-wide Torch CUT&Tag peak coordinates and annotations specifically detected in Torch-expressing samples: *D. melanogaster GMR-GAL4* female heads co-expressing *kibra* and *torch-3×NLS*, HEK293T cells expressing *torch-3×NLS*, and brains and kidneys from *torch*-expressing male mice. Promoter-proximal peaks, defined as peaks located from −3 kb to +1 kb relative to a TSS, are provided in separate sheets. FIMO results are also provided for the Torch-binding GA-repeat motif (GAGAGAGAGRGAGAG) within promoter-associated Torch CUT&Tag peaks at *kibra* in *M. pharaonis* and *D. melanogaster*, human *WWC1* in HEK293T cells, and mouse *WWC1* in brain and kidney, with motif matches reported at FIMO $P < 0.01$. MEME-ChIP de novo motif-discovery summaries and CentriMo central-enrichment results are additionally provided for Torch-3×NLS-bound peaks in HEK293T cells.

**Supplementary Dataset 14. Differential gene expression following heterologous *torch* expression in *Drosophila*, HEK293T cells and mice.** This dataset contains RNA-seq differential expression results comparing *D. melanogaster GMR-GAL4* female heads co-expressing *kibra* and *torch-3×NLS* with *GMR>kibra* controls; HEK293T cells stably expressing *torch* or *torch–3×NLS* with control HEK293T cells carrying a genome-integrated empty-vector construct; and brains, kidneys, livers and testes from *torch*-expressing male mice with the corresponding CMV-Cre-only controls. Positive $\log_2$ fold-change values indicate higher expression in *torch*-overexpressing (OE) samples, and DEGs (adjusted $P < 0.05$) are provided in separate sheets. For read alignment and quantification, the *M. pharaonis torch* sequence (LOC105839887), which is absent from the native host genomes, was appended to each host reference genome and gene annotation so that *torch* reads were quantified alongside endogenous genes. *torch* was significantly upregulated in every *torch*-expressing group relative to its corresponding control, independently confirming successful transgene expression in all samples. A separate summary lists the nine conserved direct Torch targets shared across the CUT&Tag-profiled heterologous systems, all of which were downregulated in response to Torch expression.

**Supplementary Dataset 15. GAGE-based KEGG pathway enrichment following heterologous *torch* expression in *Drosophila*, HEK293T cells and mice.** This dataset contains GAGE enrichment results for all KEGG pathways tested in the heterologous RNA-seq comparisons reported in Supplementary Dataset 14.
